# Local genetic correlation analysis reveals heterogeneous etiologic sharing of complex traits

**DOI:** 10.1101/2020.05.08.084475

**Authors:** Yiliang Zhang, Qiongshi Lu, Yixuan Ye, Kunling Huang, Wei Liu, Yuchang Wu, Xiaoyuan Zhong, Boyang Li, Zhaolong Yu, Brittany G. Travers, Donna M. Werling, James J. Li, Hongyu Zhao

**Author notes:** To whom correspondence should be addressed: Dr. Hongyu Zhao, Department of Biostatistics, Yale School of Public Health, 60 College Street, New Haven, CT, 06520, USA. These authors contributed equally to this work (Yiliang Zhang, Qiongshi Lu).

## Abstract

Local genetic correlation quantifies the genetic similarity of complex traits in specific genomic regions, which could shed unique light on etiologic sharing and provide additional mechanistic insights into the genetic basis of complex traits compared to global genetic correlation. However, accurate estimation of local genetic correlation remains challenging, in part due to extensive linkage disequilibrium in local genomic regions and pervasive sample overlap across studies. We introduce SUPERGNOVA, a unified framework to estimate both global and local genetic correlations using summary statistics from genome-wide association studies. Through extensive simulations and analyses of 30 complex traits, we demonstrate that SUPERGNOVA substantially outperforms existing methods and identifies 150 trait pairs with significant local genetic correlations. In particular, we show that the positive, consistently-identified, yet paradoxical genetic correlation between autism spectrum disorder and cognitive performance could be explained by two etiologically-distinct genetic signatures with bidirectional local genetic correlations. We believe that statistically-rigorous local genetic correlation analysis could accelerate progress in complex trait genetics research.

## Introduction

Genome-wide association study (GWAS) has achieved remarkable success in the past 15 years and has identified numerous single-nucleotide polymorphisms (SNPs) associated with complex human traits and diseases^1^. Increasingly accessible summary statistics from GWAS, in conjunction with advances in analytical methods that use marginal association statistics as input, have circumvented logistical challenges in data sharing and greatly accelerated research in complex trait genetics^2^.

With these advancements, multi-trait modeling has undergone rapid developments, leading to the emergence of numerous methods that study the shared genetic basis across multiple phenotypes^3-8^. Among these methods, genetic correlation analysis is a statistically powerful and biologically interpretable approach to quantifying the overall genetic similarity of two traits^9-15^. It has gained popularity in the field, provided new insights into the shared genetics of many phenotypes^10,16^, and has a variety of downstream applications^9^. Properly modeling genetic correlation could enhance statistical power in genetic association studies^3,4^, improve risk prediction accuracy^17-19^, and facilitate causal inference and mediation analysis^5,7,20-22^. A number of methods have been developed for genetic correlation estimation. Built upon the GREML approach^14,23^, cross-trait linkage disequilibrium (LD) score regression (LDSC) was the first method that uses GWAS summary statistics alone as input^10,24^. Methods have also been developed to estimate annotation-stratified^12^ and trans-ethnic^13^ genetic correlation. Bioinformatics servers have been built to improve the computation and visualization of genetic correlations^25^.

Local genetic correlation analysis is another important approach to tackling the underlying etiological mechanisms shared by multiple complex traits^11,26^. Instead of estimating the average correlation of genetic effects across the genome, local genetic correlation quantifies the genetic similarity of two traits in specific genomic regions. This approach could reveal local, heterogenous architecture of etiological sharing and is critical for understanding the heterogeneity in pleiotropic genetic effects. Existing methods have struggled to provide statistically principled and robust results due to technical challenges including extensive LD in local chromosomal regions and pervasive sample overlap across GWASs.

Here, we introduce a novel statistical framework named SUPERGNOVA for local genetic correlation estimation. Based on the GNOVA approach which was designed for partitioning genetic correlation by functional annotation^12^, SUPERGNOVA is a principled framework for diverse types of genetic correlation analyses. Through extensive simulations, we demonstrate that SUPERGNOVA provides statistically rigorous and computationally efficient inference for both global and local genetic correlations and substantially outperforms existing methods when applied to local genomic regions. Additionally, our approach uses GWAS summary statistics alone as input, and is robust to overlapping GWAS samples even when the shared sample size is unknown. We applied SUPERGNOVA to 30 complex traits and report 150 pairs of phenotypes with significant local genetic correlations. In particular, we investigated an empirical paradox – the robust, positive genetic correlation between autism spectrum disorder (ASD) and cognitive ability, which contradicts the comorbidity between ASD and intellectual disability^27^. We demonstrate that multiple distinct etiologic pathways contribute to the shared genetics between ASD and cognitive ability which could only be revealed by genetic correlation analysis at a local scale.

## Results

### Overview of SUPERGNOVA analytical framework

Genetic covariance (correlation) is defined as the covariance (correlation) of genetic effects on two traits. It is commonly used as an informative metric to quantify the shared genetic basis between traits. Given the marginal association statistics from two GWASs (i.e. z-scores *z*_1_ and *z*_2_), genetic covariance *ρ* between two traits can be estimated by minimizing the “distance” between the empirical covariance of z scores, i.e., 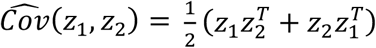, and the theoretical covariance

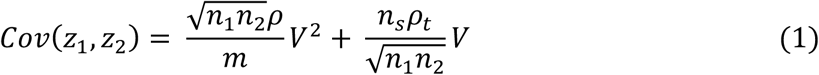

where *m* is the number of SNPs, *n*_1_ and *n*_2_ are the sample sizes of two GWASs, *n*_*s*_ is the number of individuals included in both studies, *V* is the LD matrix, and *ρ*_*t*_ = *ρ* + *ρ*_*e*_ is the sum of genetic covariance (i.e. *ρ*) and the covariance of non-genetic effects (i.e. *ρ*_*e*_) on the two traits among shared individuals. Derivation of the theoretical covariance and other statistical details are reported in the **Supplementary Note**. In the **Methods** section, we show that with different definitions of “distance”, existing methods such as LDSC^10^ and GNOVA^12^ are special cases of this unified framework.

Local genetic covariance (correlation) can be defined in a similar way by focusing only on SNPs in a pre-specified genomic region (**Methods**). Despite the conceptual similarity between global and local genetic correlation, local z-scores from each GWAS can be highly correlated due to the extensive LD in local regions. Hence, most methods developed for global genetic correlation cannot be directly applied to estimate local correlations. In addition, ubiquitous sample overlap across GWASs introduces technical correlations among association statistics from different studies, which further complicates the estimation of genetic correlation, especially in local regions. SUPERGNOVA resolves these statistical challenges by decorrelating local z-scores with eigenvectors of the local LD matrix (**Figure 1**). In practice, LD can be estimated from an external reference panel (e.g., 1000 Genomes Project^28^). Due to the noise in LD estimation, we only use the first *K*_*i*_ eigenvectors to transform and decorrelate association statistics in any given region *i* where *K*_*i*_ can be determined adaptively in SUPERGNOVA. After decorrelation, local genetic covariance *ρ*_*i*_ is estimated through a weighted least squares regression in each region. When the number of SNPs is large, SUPERGNOVA is equivalent to GNOVA (**Supplementary Note**). Another technical challenge is that numerically unstable estimates of local heritability will lead to extreme variability in the estimates of local genetic correlation. Therefore, we base our inference on local genetic covariance which is statistically equivalent. We discuss more statistical details in the **Methods** section.

**Figure 1.**
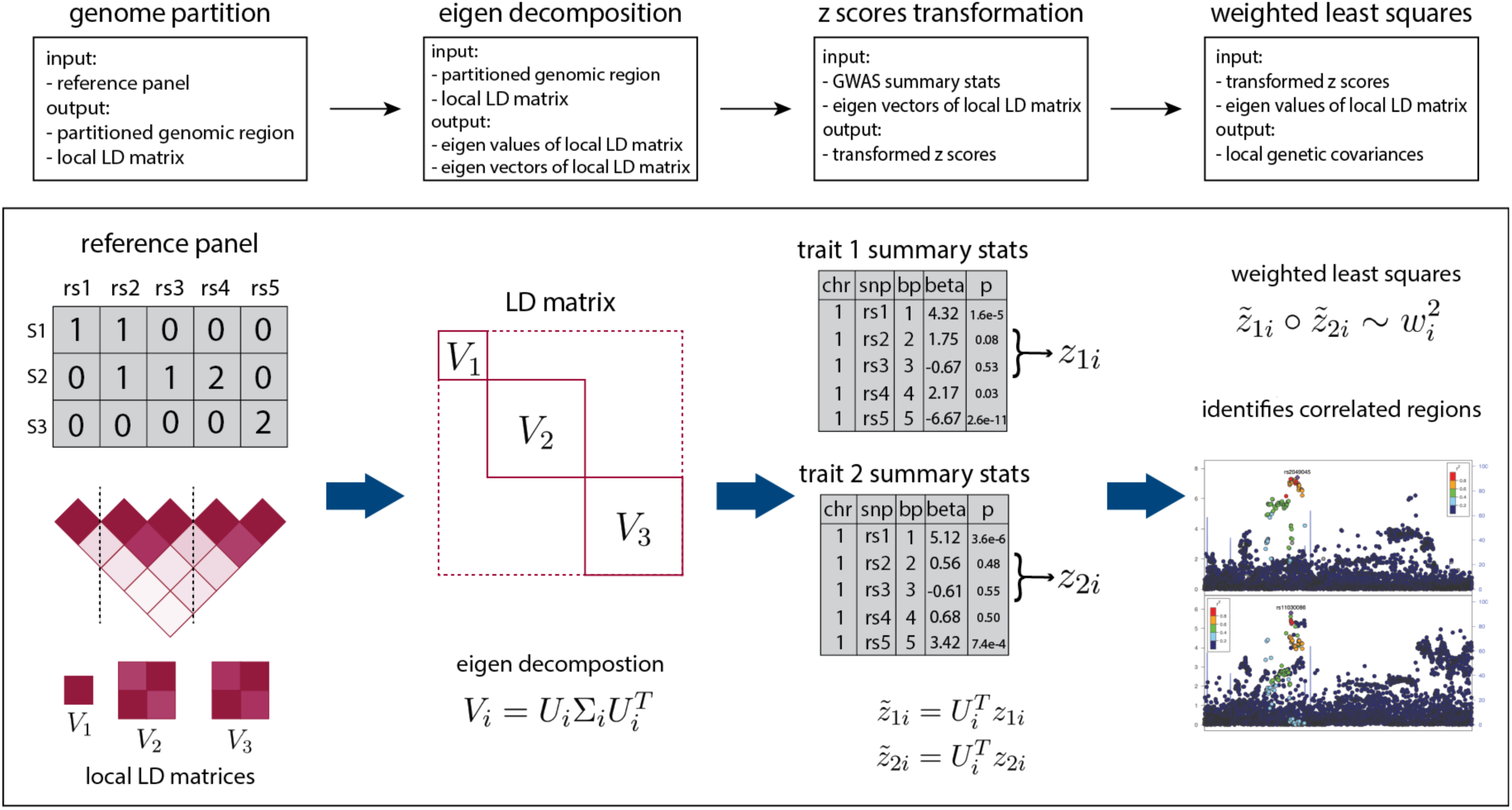
SUPERGNOVA workflow. Details on the statistical framework are described in the Methods section. *w*_*i*_ denotes the diagonal elements of Σ_*i*_, which are also the eigenvalues of each local LD matrix. Notation, ∘ in the last step indicates the element-wise product.

### Simulations

We performed simulations to assess the performance of SUPERGNOVA for both global and local genetic correlation analyses. We compared SUPERGNOVA with multiple state-of-the-art methods in six different simulation settings and repeated each setting 100 times. We used real genotype data from the Wellcome Trust Case Control Consortium (WTCCC) to simulate quantitative traits. After quality control, 15,918 samples and 287,539 SNPs remained in the dataset. We equally divided 15,918 samples into two subsets which we denote as set 1 and set 2. To assess the robustness of our approach to sample overlap between GWASs, we generated another dataset by combining 3,979 samples from set 1 and 3,980 samples from set 2. We refer to it as set 3. This results in a 50% sample overlap between set 1 and set 3. Detailed simulation settings and quality control procedures are described in the **Methods** section.

We compared the performance of LDSC, GNOVA, and SUPERGNOVA on global genetic covariance estimation. Both SUPERGNOVA and GNOVA showed superior statistical power compared to LDSC in all settings (**Figures 2A-C**). No method showed inflated type I error rates when the true covariance was 0. All three approaches provided unbiased estimates for global genetic covariance but LDSC estimates had substantially larger variance compared to GNOVA and SUPERGNOVA (**Supplementary Figures 1-3**).

**Figure 2.**
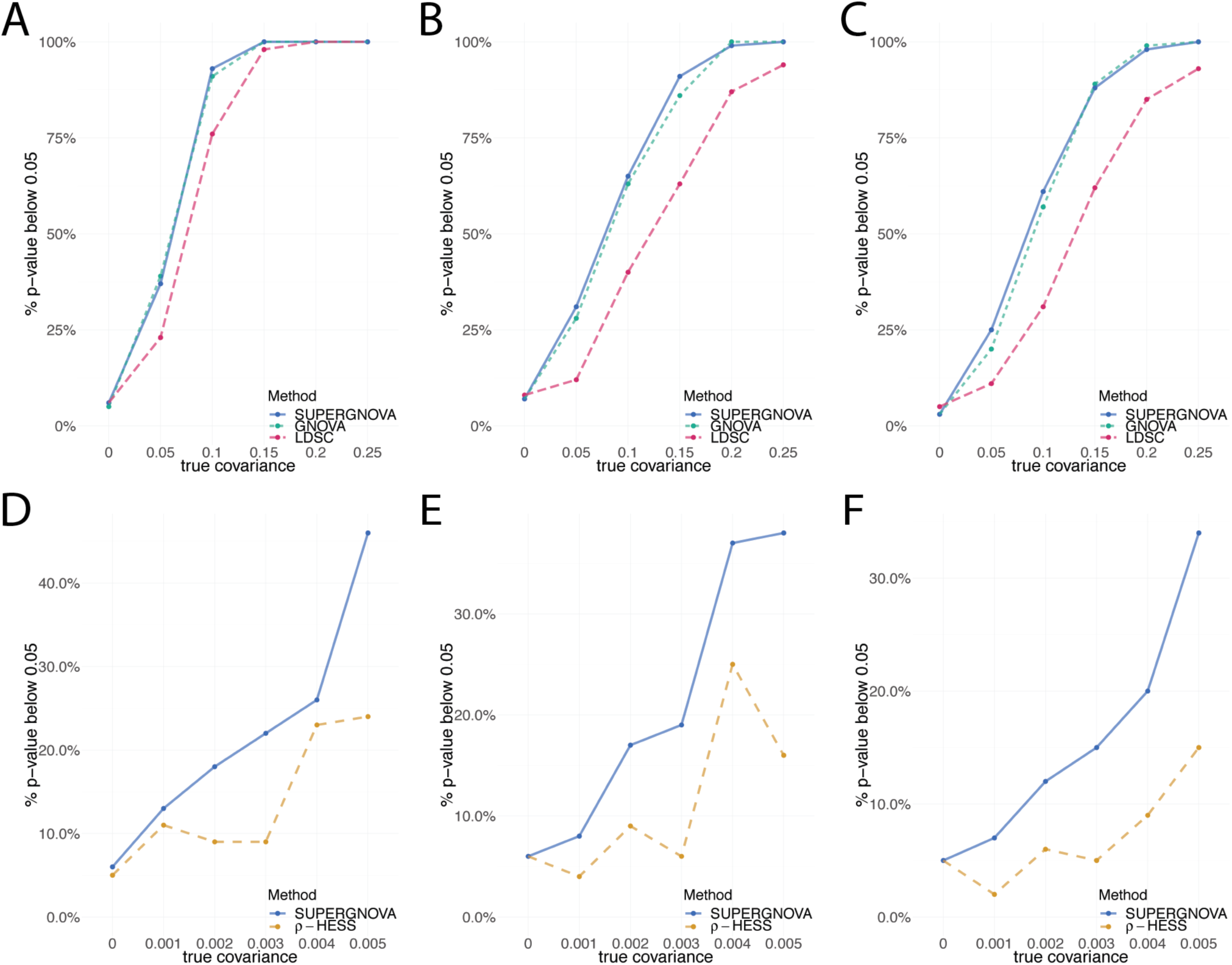
Simulation results. Panels A-C compare the type-I error and statistical power of SUPERGNOVA, GNOVA, and LDSC in global genetic covariance estimation. **(A)** Two GWASs were simulated on two non-overlapping datasets (set 1 and set 2). **(B)** GWASs were simulated on two datasets with a 50% sample overlap (set 1 and set 3). **(C)** Two GWASs were both simulated on the same dataset (set 1) with a 100% sample overlap. Panels D-F compare SUPERGNOVA and *ρ*-HESS on local genetic covariance estimation using GWASs **(D)** without sample overlap (set 1 and set 2), **(E)** with a partial sample overlap (set 1 and set 3), and **(F)** with a complete sample overlap (set 1 only).

Next, we compared *ρ*-HESS and SUPERGNOVA on their performance of estimating local genetic covariance. We used 395 SNPs from a selected genomic region of about 3.3 Mb on chromosome 2 as the local region of interest. The remaining SNPs on chromosome 2 (23,839 SNPs) were used as the “background SNPs” in the analysis. We set the covariance in the small local region to be from 0 to 0.005. Outside of this region on chromosome 2, covariance was fixed as 0. The total heritability was set to be 0.5 and was equally distributed among all SNPs on chromosome 2 (24,234 SNPs). When there is no overlapping sample between two studies, SUPERGNOVA estimates were unbiased, showing well-controlled type I error and good statistical power. On the other hand, *ρ*-HESS consistently underestimated local genetic covariance and had lower statistical power (**Figure 2D**; **Supplementary Figure 4**).

We repeated these simulations in set 1 and set 3 with a 50% sample overlap. SUPERGNOVA estimates of local genetic covariance remained unbiased with well-controlled type I error (**Supplementary Figure 5**). Compared to *ρ* -HESS, SUPERGNOVA showed superior statistical power (**Figure 2E**). *ρ*-HESS underestimated local genetic covariance even though we provided the correct sample size for shared samples (**Supplementary Figure 5**). We also performed simulations under a complete sample overlap by simulating two traits on set 1. SUPERGNOVA still achieved unbiased estimation and valid inference (**Supplementary Figure 6** and **Figure 2F**). *ρ*-HESS lacked statistical power in all settings. Additionally, *ρ*-HESS was not robust to mis-specified overlapping sample size and suffered from type I error inflation when provided with inaccurate values of the overlapping sample size and phenotypic correlation (**Supplementary Figures 7-8**; **Methods**). We note that SUPERGNOVA does not need the shared sample size or phenotypic correlation as input.

### Global and local genetic correlations among 30 complex traits

We applied SUPERGNOVA to estimate local and global genetic correlations among 30 phenotypes (**Supplementary Table 1**). We partitioned the genome into 2,353 approximately independent regions (about 1.6 centimorgan on average) using LDetect^29^, with LD estimated from the 1000 Genomes Project phase III samples of European ancestry^28^. 127 pairs of traits were globally correlated (p < 0.05/435 = 1.1e-4; **Supplementary Figure 9**) and 150 pairs of traits were locally correlated in 109 different regions under Bonferroni correction (p < 0.05/1,006,072 = 5.0e-8; **Figure 3A** and **Supplementary Tables 2-3**).

**Figure 3.**
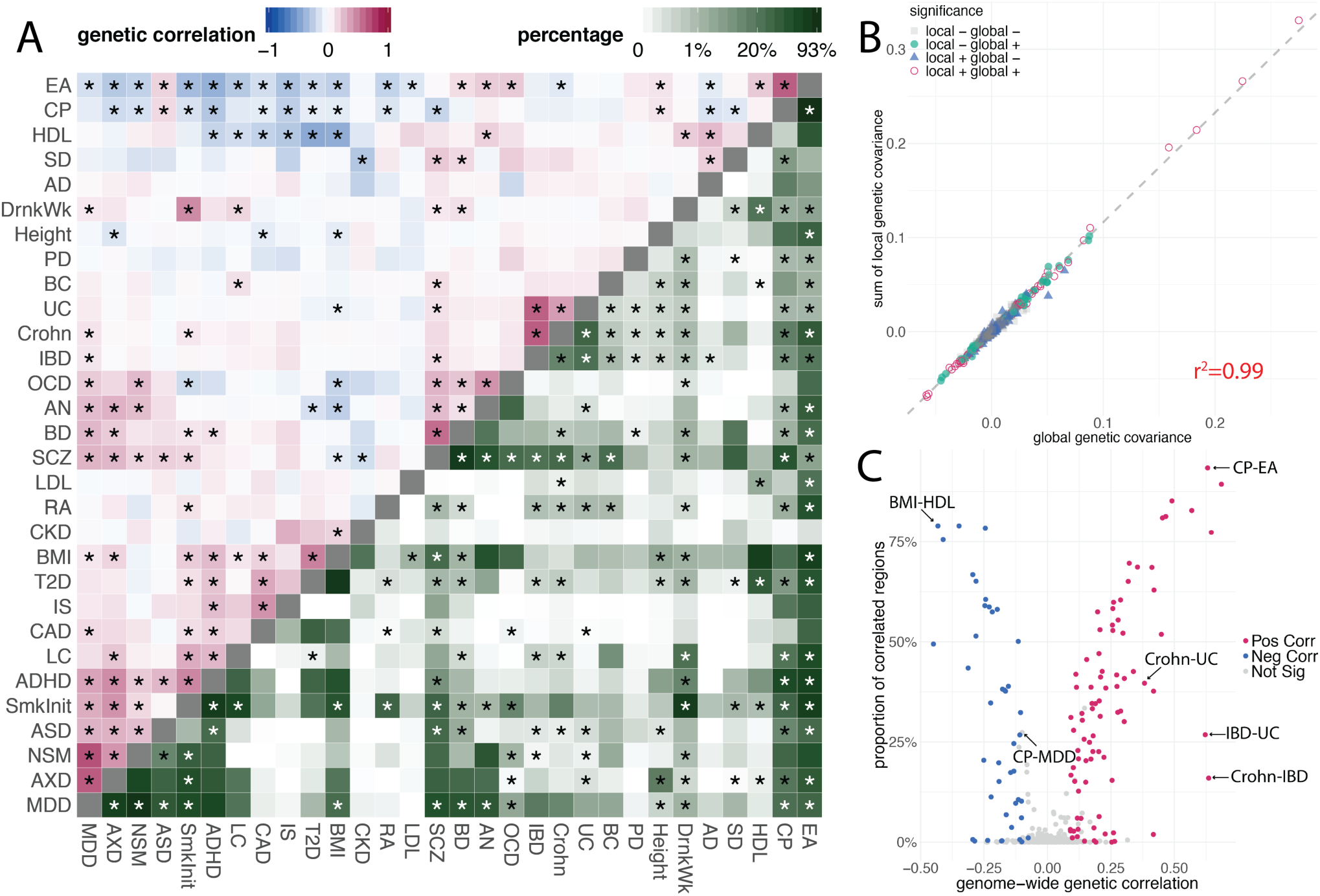
Global and local genetic correlations among 30 complex traits. **(A)** Estimates of global genetic correlations (upper triangle) and estimated proportions of correlated regions among 435 trait pairs (lower triangle). Asterisks in the upper triangle highlight significant genetic correlations after Bonferroni correction for 435 pairs. Asterisks in the lower triangle indicate at least one significantly correlated region between the traits after Bonferroni correction for all 1,006,072 regions in 435 trait pairs. We grouped traits with hierarchical clustering applied to global genetic correlations. **(B)** Global genetic covariance estimates were highly concordant with the sums of local genetic covariance. Each point represents a trait pair. Color and shape of each data point denote the significance status in global and local correlation analyses. **(C)** Volcano plot comparing the global genetic correlation and proportion of correlated local regions. Each point represents a trait pair. Color of each data point represents the significance and direction of global correlation. Trait pairs discussed in the main text are labeled in the plot.

The sums of local covariance across 2,353 regions were highly concordant with the estimated global genetic covariance (**Figure 3B**; R^2^=0.99), but local genetic covariance revealed diverse architecture of genetic sharing locally. We estimated the proportion of correlated regions for each pair of traits using ashr^30^ (**Methods**; **Figure 3A**; **Supplementary Table 4**). The proportion of correlated regions predicted global genetic correlation in general, with some notable outlier trait pairs (**Figure 3C**). Two subtypes of inflammatory bowel disease (IBD), Crohn’s disease and ulcerative colitis (UC), had strong pairwise global correlations but relatively sparse local genetic correlations (**Supplementary Figure 10**). In fact, all 8 identified regions were positively correlated among Crohn’s disease, UC, and IBD and harbored genome-wide significant loci reported in the GWAS on IBD^31^, suggesting that a limited fraction of the genome contribute to different subtypes of IBD with strong and concordant effects. In contrast, SNPs in 93% of regions had correlated effects between cognitive performance (CP) and educational attainment (EA; global genetic correlation=0.63; p=6.1e-115), the highest among all trait pairs. 79% of regions showed correlated effects between body mass index (BMI) and high-density lipoprotein (HDL) cholesterol (global genetic correlation=-0.43; p=3.6e-41), the highest among negatively correlated traits. These results suggest extensive and ‘omnigenic’ genetic sharing between these traits, which is also reflected in the substantial shift in the distribution of local genetic covariances (**Supplementary Figure 10**). Bidirectional correlations were also observed in several trait pairs, including ASD and CP^32^ (**Supplementary Figure 10**; global correlation=0.15; 15% of regions were correlated). Across all trait pairs, we observed a modest association between the sample size of traits and the proportion of correlated regions (**Supplementary Figure 11**).

We identified significant local genetic covariance for 86 trait pairs that were not significantly correlated in the global analysis (**Supplementary Table 2-3; Supplementary Figure 12**), including HDL cholesterol and low-density lipoprotein (LDL) cholesterol^11^, CP and major depressive disorder (MDD), obsessive-compulsive disorder (OCD) and anxiety disorder (AXD), and ASD and bipolar disorder (BD).

Our analyses also implicated several genomic regions showing correlated genetic effects on more than two traits. The *BDNF* locus on chromosome 11 (hg19 coordinate: 27,019,873-28,741,185) is known to control the development of neurons and synapses and is vital to learning, memory, and vulnerability to stress^33-36^. We identified significant genetic covariance at this locus between 6 trait pairs among schizophrenia (SCZ), EA, smoking initiation (SmkInit), drinks per week (DrnkWk), and ADHD (**Supplementary Figure 13-14**). Another locus on chromosome 11 (111,985,737-113,103,996) was identified among 7 neuropsychiatric traits: anorexia nervosa (AN), BD, MDD, CP, SCZ, SmkInit, and neuroticism (NSM) (**Supplementary Figure 15-16**). *NCAM1* at this locus is involved in development and maintenance of the nervous system and is associated with SCZ and comorbid alcohol and drug dependence^37-39^. These hub regions with pervasive correlations among psychiatric disorders hint at key regulators in the nervous system and provide guidance to functional genomic studies that interrogate the mechanisms of pleiotropic effects^40^.

Local genetic covariance that did not achieve statistical significance may still be worth follow-up investigations. Despite evidence on phenotypic correlations, previous studies have suggested that Alzheimer’s disease (AD) is not genetically correlated with neuropsychiatric traits except education and cognition^16^. We identified suggestive local correlations of AD with 7 neuropsychiatric traits: NSM (p=1.2e-6), OCD (p=3.0e-6), CP (p=3.4e-6), DrnkWk (p=2.0e-5), MDD (p=2.7e-5), and AN (p=4.2e-4), at the *SPI1* locus (chr11: 46,876,411-48,200,127). We replicated the local correlations with DrnkWk (p=7.5e-2) and NSM (p=1.4e-2) using an independent GWAS of AD family history (**Methods**; **Supplementary Tables 5-6**). The estimates for local genetic covariance were highly consistent between two analyses (R^2^=0.84; **Supplementary Figure 17**). The *SPI1* locus has been consistently identified in AD GWASs^41,42^. A recent genome-wide survival study of AD onset convincingly demonstrated that transcription factor (TF) PU.1 encoded by *SPI1* is a key regulator for the development and function of myeloid cells and lower *SPI1* expression delays the onset of AD by regulating gene expression in myeloid cells^43^. However, genetic covariance of AD with DrnkWk and NSM was not statistically significant in TF binding sites of PU.1 in macrophages and monocytes (**Methods**; **Supplementary Table 7**). Transcriptome-wide association study (TWAS) identified a number of PU.1-regulated genes associated with these phenotypes in macrophages and monocytes (**Methods**; **Supplementary Tables 8-9**), but all the genes shared by multiple traits are located at the *SPI1* locus (**Supplementary Figure 17**). These results suggest that although the *SPI1* locus may have correlated roles in multiple psychiatric and neurodegenerative diseases, PU.1 may modulate the risk of these diseases through regulating the transcription of distinct susceptible genes in myeloid cells.

### Dissecting the shared genetic basis of ASD and cognitive ability

We further demonstrate the power of SUPERGNOVA through an in-depth case study of the shared genetics between ASD and cognitive ability. Paradoxically, previous studies based on multiple different approaches have found a positive genetic correlation between ASD and CP^16,44,45^. We also identified significant positive global genetic correlations between ASD and measures of cognitive ability (**Figure 3**), e.g., CP (standardized score on neuropsychological tests; correlation=0.15, p=3.2e-8) and EA (years of schooling; correlation=0.18, p=3.8e-14). Cognitive phenotypes in these GWASs have been previously described in detail^46^. However, such a positive correlation contradicts the known comorbidity of intellectual disability and ASD with regard to *de novo* variants of high penetrance^27,47^. In addition, other neurodevelopmental disorders such as ADHD showed negative genetic correlations with cognitive measures (correlation=-0.29 and p=2.9e-29 with CP; correlation=-0.41 and p=2.0e-59 with EA), but the genetic correlation between ASD and ADHD was positive (correlation=0.28, p=2.3e-9).

A total of 64 genomic regions with significant local genetic covariance were identified among ADHD, ASD, and CP at a false discovery rate (FDR) cutoff of 0.1 (**Supplementary Table 10**; **Supplementary Figure 18**). The local covariances of CP with ASD and ADHD were bidirectional. No region with a negative covariance between ASD and ADHD was identified. The paradox that ASD and ADHD show opposite correlations with CP was not observed in any local region (**Figure 4A**; **Supplementary Table 11**). 18 regions showed significant positive correlations between ASD and CP, among which 3 regions were also significant and positive between ADHD and ASD and 2 regions were significant and positive between ADHD and CP. Similarly, we identified 32 regions with significant negative correlations between ADHD and CP. Among these regions, 3 were positive between ADHD and ASD and 3 were negative between ASD and CP. Three regions reached statistical significance in all three trait pairs. ASD and ADHD were positively correlated in all three regions. ASD and ADHD were both positively correlated with CP in the regions on chromosomes 4 (150,634,191-153,226,998) and 14 (36,683,516-38,481,516) (**Supplementary Figures 19-20**) and were both negatively correlated with CP in the region on chromosome 7 (104,158,491-105,425,027) (**Figure 4B; Supplementary Figure 21**).

**Figure 4.**
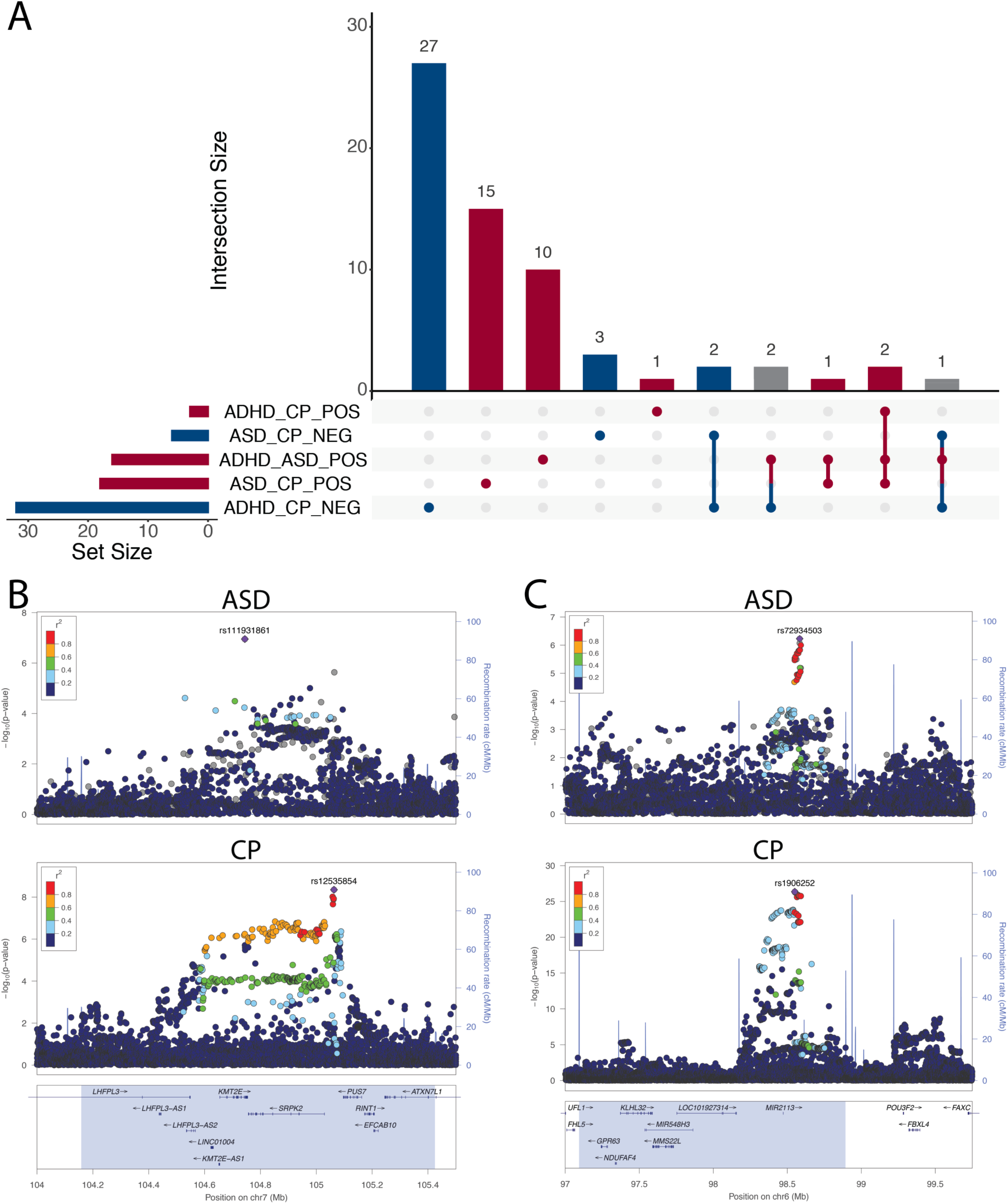
Bidirectional local genetic covariance between ASD and CP. **(A)** Regions with significant local genetic covariance among ADHD, ASD, and CP (FDR < 0.1). This plot uses bars to break down the Venn diagram of overlapped regions in different categories. The five categories shown in the lower panel are correlated regions of ADHD and CP (positive and negative), ASD and CP (positive and negative), and ASD and CP (positive only). We use different colors (red, blue, and gray) to annotate region categories of positive, negative, and mixed covariance directions. **(B)** LocusZoom plots for ASD and CP GWAS associations at the *KMT2E* locus. ASD and CP are negatively correlated in the highlighted region. **(C)** LocusZoom plots for ASD and CP at the *POU3F2* locus. ASD and CP are positively correlated in the highlighted region. *POU3F2* is 700 kb downstream of the GWAS association peak.

The locus on chromosome 7 (104,158,491-105,425,027) showed significant and negative correlations between CP and both neurodevelopmental disorders (**Supplementary Table 11).** We also identified a significant negative correlation of CP and SCZ in this region (p=1.8e-8; **Supplementary Table 12; Supplementary Figure 21**). Among genes at this locus, *PUS7* is associated with intellectual disability and neurological defects^48^. *De novo* mutations in *KMT2E* cause a spectrum of neurodevelopmental disorders including ASD^49^. An intronic SNP in *KMT2E*, rs111931861, with a minor allele frequency (MAF) of 0.034, reached genome-wide significance in a recent ASD GWAS^44^ (**Figure 4B**). *KMT2E* was also implicated by a recent exome sequencing study^50^. It is the only gene that reached genome-wide significance in both GWAS and exome sequencing studies of ASD. TWAS did not identify any genes associated with ADHD, ASD, SCZ, or CP in this region (**Supplementary Tables 13; Methods**). These results, coupled with the findings about *de novo* and ultra-rare variants in *KMT2E*, suggest that common variants in this region may be tagging protein-altering variants instead of regulatory variants for transcriptional activities. A missense SNP in *KMT2E*, rs117986340, was nominally associated with ASD (p=5.7e-2) and ADHD (p=4.4e-2) in GWAS (**Supplementary Table 14**; **Methods**) but this hypothesis needs to be investigated in the future using sequencing data.

*POU3F2* (also known as *BRN2*) is a key TF in the central nervous system and a master regulator of gene expression changes in BD and SCZ^51,52^. It is the first genome-wide significant locus identified for EA^53^. It has also been identified in a recent TWAS for ASD^54^. In our analysis, the *POU3F2* locus on chromosome 6 (97,093,295-98,893,182) showed significant positive correlations between ASD and CP (p=1.8e-5; **Figure 4C**) and among many neuropsychiatric phenotypes including AN, BD, DrnkWk, EA, and SmkInit (**Supplementary Table 12**; **Supplementary Figures 22-23**). In addition, genes in other regions showing nominal negative correlations between ASD and CP were significantly enriched for *POU3F2* protein-protein interactors (PPIs) (odds ratio=24.8; p=2.8e-3; **Methods**). This is consistent with our recent finding that genes regulated by TF *POU3F2* showed a 2.7-fold enrichment for loss-of-function *de novo* mutations in ASD probands which are known to cause comorbid intellectual disability^54^. These results hint at a pervasive, regulatory role of *POU3F2* in cognitive ability and many neuropsychiatric disorders^55,56^.

Regions showing opposite correlation directions between ASD and CP were enriched for distinct mechanistic pathways (**Methods**; **Figure 5**; **Supplementary Tables 15-18**). Genomic regions with negative correlations between ASD and CP were significantly enriched for chromatin modifier genes (enrichment=3.2; p=3.8e-4; **Supplementary Table 15**). *De novo* protein-truncating mutations in these genes are known to cause ASD, intellectual disability, and a variety of congenital anomalies^27,57,58^. Regions positively correlated between ASD and CP were significantly enriched for postsynaptic density (PSD) proteins (enrichment=1.8; p=3.5e-4; **Supplementary Table 16**). *FMRP* targets also showed a significant enrichment in positively correlated regions (enrichment=1.9; p-value=2.7e-3; **Supplementary Table 16**). The enrichment of *FMRP* targets in negatively correlated regions was comparable but did not reach statistical significance after multiple testing correction (enrichment=1.8; p-value=0.032; **Supplementary Tables 15**). PSD genes are known to be enriched for associations identified in ASD TWAS^54^. *FMRP* targets are enriched for both ASD heritability quantified using common variants^59^ and *de novo* mutations of ASD^60,61^. *FMRP* target genes showed a 12.4-fold enrichment (p=3.5e-15) in the 102 risk genes identified in the latest exome sequencing study of ASD^50^. Notably, findings from exome-sequencing studies (e.g., the 102 ASD genes^50^) and gene sets known to be enriched for ultra-rare or *de novo* protein-truncating variants in ASD probands (e.g., chromatin modifiers^27^) showed substantially stronger enrichment in the regions with negative ASD-CP correlations than the regions with positive correlations (**Supplementary Tables 15-16**).

**Figure 5.**
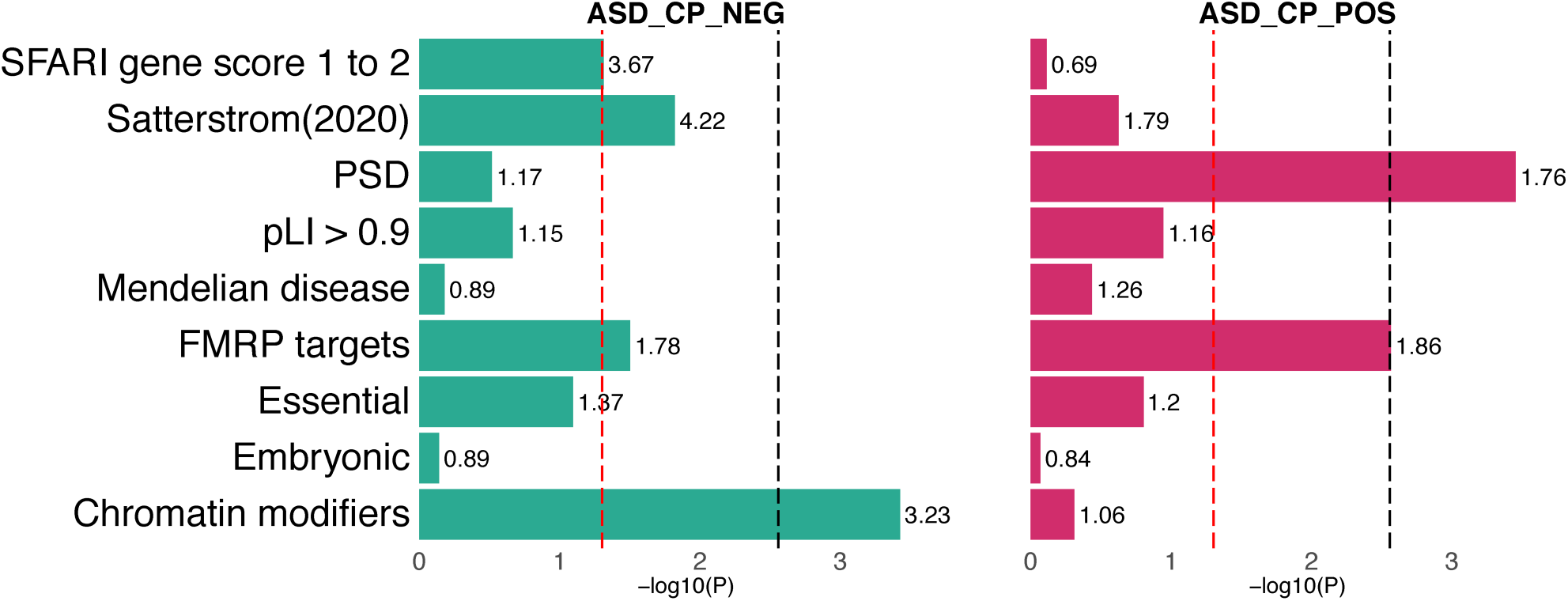
Enrichment for gene sets in correlated regions between ASD and CP. Regions with opposite correlations between ASD and CP were enriched for different mechanistic pathways. Fold enrichment values are labeled next to each bar. The red dashed lines mark the p-value cutoff of 0.05 and the black dashed lines denote the p-value thresholds after Bonferroni correction (p=2.8e-3).

We then assessed the enrichment of associations for other complex traits in genetically correlated regions between ASD and CP (**Methods**). Regions with positive correlations between ASD and CP were significantly enriched for associations for 10 traits documented in GWAS Catalog (p<0.05/664=7.5e-5), including extremely high intelligence (odds ratio=9.7; p=3.5e-9), household income (odds ratio=52.6; adjusted p=5.7e-9), and loneliness (odds ratio=5.5; p=4.1e-5) (**Supplementary Table 19; Supplementary Figure 24**). Negatively correlated regions were enriched for associations with a variety of neurodevelopmental and psychiatric disorders including SCZ (odds ratio=6.1; p=2.2e-31), BD (odds ratio=13.3; p=2.7e-24), and NSM (odds ratio=10.0; p=6.0e-12) (**Supplementary Table 20; Supplementary Figure 24**). We also estimated stratified genetic covariance of 28 other traits with ASD and CP in these identified regions (**Supplementary Figure 25**). EA, MDD, and rheumatoid arthritis (RA) showed significant stratified covariance with ASD or CP (p<0.05/112=4.5e-4) in regions positively correlated between ASD and CP (**Supplementary Table 21**). On the other hand, ADHD, EA, and AXD showed significant stratified covariance with ASD or CP in regions showing significant negative correlations between ASD and CP (**Supplementary Table 22**). Overall, traits showed consistent covariances with ASD and CP in regions with positive ASD-CP covariances, while the covariances with ASD and CP have opposite directions in regions with negative ASD-CP covariances (**Supplementary Figure 25**). In other words, no paradoxical covariances were present when we zoomed in by ASD-CP correlated regions.

Genes in positively correlated regions of ASD and CP were expressed in a substantially higher proportion of cells in fetal brains compared to background genes (p=0.012; log-rank test) (**Methods**; **Supplementary Figure 26; Supplementary Table 23**) while the elevation of gene expression rate in negatively correlated regions was not significant (p=0.15, log-rank test). We did not identify a significant difference in the expression rate between genes in the ASD-CP positively correlated regions and genes in the ASD-CP negatively correlated regions (p=0.71, log-rank test). The average expression of both gene sets was significantly higher than background genes across prenatal and postnatal stages (p=9.7e-525 and 2.5e-99 for genes in positively and negatively correlated regions, respectively) (**Methods; Supplementary Figure 27**). We also identified significantly higher expression of genes in positively correlated regions than in negatively correlated regions across developmental stages (p=1.92e-61; **Supplementary Figure 27**). We did not identify differential expression between prenatal and postnatal brains for either gene set (p=0.83 and 0.81; **Methods; Supplementary Table 24**).

We then investigated the phenotypic heterogeneity of ASD probands with different genetic signatures. We constructed two polygenic risk scores (PRSs) of ASD based on independent SNPs from genomic regions with positive and negative local correlations between ASD and CP, respectively, for 5,469 ASD probands and 2,132 healthy siblings in the Simons Foundation Powering Autism Research for Knowledge (SPARK) cohort (**Methods**). We refer to these scores as PRS+ and PRS-. Both PRS+ and PRS- are normally distributed in SPARK (**Supplementary Figure 28**). PRS+ could significantly distinguish ASD probands and healthy siblings (odds ratio= 1.08; p=0.026) while the association between PRS- and ASD status was not significant (odds ratio=1.02; p=0.71; **Methods**). 1,803 probands had both genotype data and intelligence quotient (IQ) information. Probands with high PRS+ had higher IQ compared to probands with high PRS-, with the average IQ changing sharply in the right tail of the PRS distribution, from 93.8 and 94.7 (p=0.64; two-sample t-test) in the 75% percentile to 101.7 and 84.0 (p=0.046) in the 99% percentile (**Figure 6A; Supplementary Figure 29**). At the 99% percentile, the proband subgroups with extreme PRS+ and PRS-did not have overlapping samples. 10.5% of probands with extreme PRS+ and 31.6% of probands with extreme PRS-had an IQ below 70 (**Figures 6A and 6B**). Four probands in the extreme PRS-group had relatively high PRS+ (greater than the 90% percentile). All of them had IQ>70 (**Figure 6B**). No proband in the extreme PRS+ group had high PRS- (**Figures 6C**).

**Figure 6.**
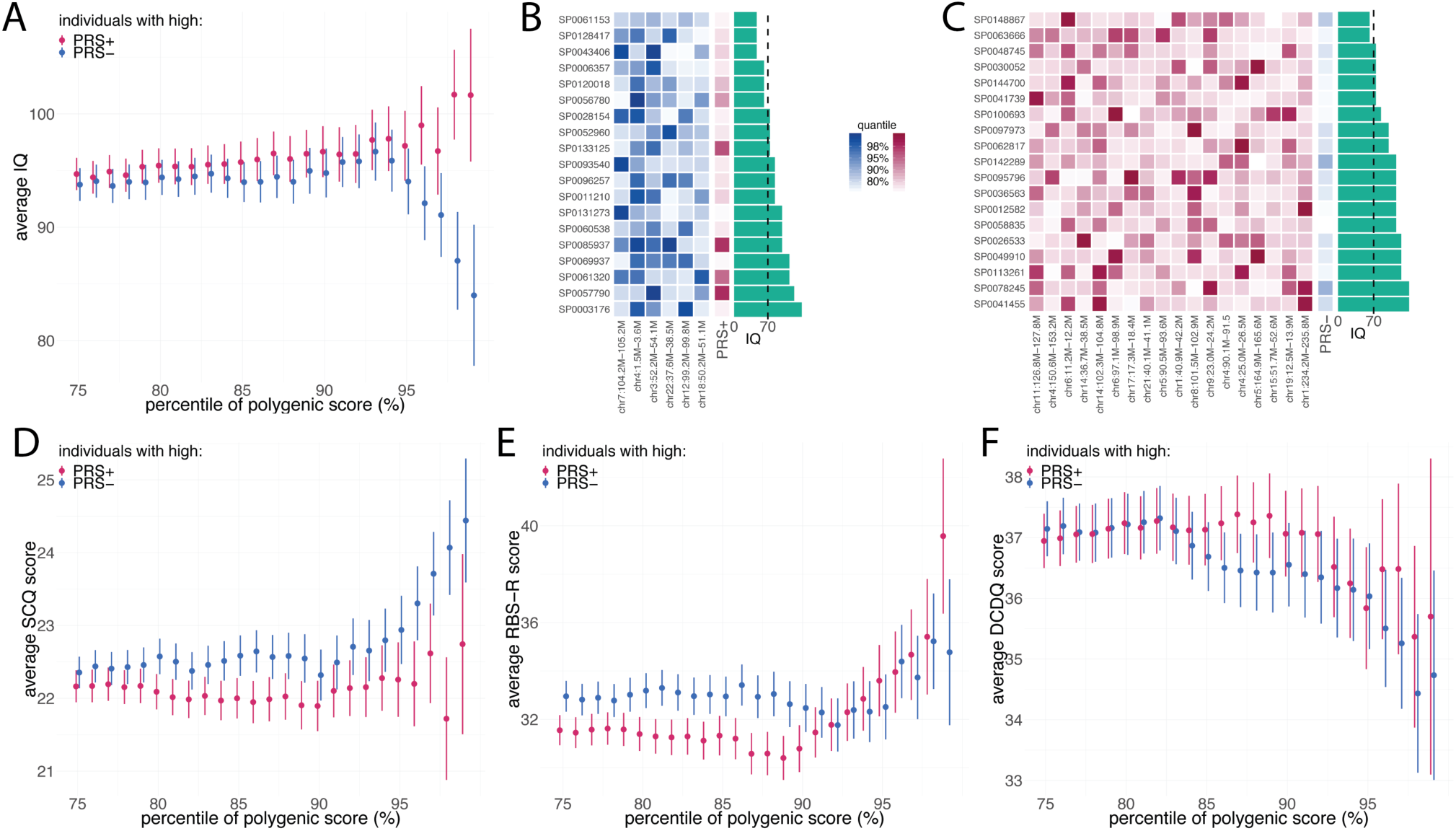
Phenotypic heterogeneity of ASD probands with high PRS+ and PRS-. **(A)** Average IQ is computed for different groups defined by PRS. Each interval indicates standard error of the estimated mean. **(B)** PRS percentiles and IQ of probands with extreme PRS-. PRS-was calculated using six negatively correlated regions between ASD and CP. The blue heatmap indicates the percentile of ASD PRS in each contributed region for each proband. The percentiles of PRS+ values are shown in the red boxes. IQ is shown as green bars. **(C)** PRS percentiles and IQ of probands with extreme PRS+. PRS+ was calculated using 18 positively correlated regions between ASD and CP and the per-locus percentiles are shown in red. The percentiles of PRS+ are shown in blue and the green bars denote IQ. In panel D-F, average phenotypic scores of ASD were computed for ASD probands at various PRS percentiles. Intervals denote standard errors for the estimated means. The phenotypic scores are **(D)** SCQ score, **(E)** RBS-R score, and **(F)** DCDQ score.

4,267 probands in SPARK had genotype data and social communication questionnaire (SCQ) scores (**Methods**). We used SCQ score as a proxy for ASD symptom severity. Probands with extreme PRS- at the 99% percentile showed significantly elevated SCQ scores compared to other probands (p=0.03; two-sample t-test) **(Supplementary Table 25**). Average SCQ score rose from 22.4 in the 75% percentile to 24.4 in the 99% percentile (**Figure 6D**). We did not identify a significant elevation in probands with extreme PRS+ (p=0.56). The repetitive behaviors scale-revised (RBS-R) questionnaire was used to quantify repetitive behaviors, including self-injuries, restricted behavior, compulsive behavior, stereotyped behavior, ritualistic behavior, and sameness behavior^62^. We observed a significant increase of RBS-R scores in probands with extreme PRS+ (p=0.016; **Figure 6E**) but not in probands with extreme PRS- (p=0.4; **Supplementary Table 25**). We also investigated motor ability quantified by the developmental coordination disorder questionnaire^63,64^ (DCDQ) in SPARK. We observed a downward trend of DCDQ score (i.e., worse motor ability) as PRS increases (**Figure 6F**) but the changes were not statistically significant (**Supplementary Table 25**). Follow-up analyses examining RBS-R and DCDQ subscales found that the pattern of results was not driven by any one of the subdomains. Finally, we assessed the enrichment of ASD subtypes in extreme PRS+ and PRS-groups. No subtype reached statistical significance (**Supplementary Table 26**), with Asperger’s disorder showing the strongest yet modest enrichment (enrichment=1.58; p=0.082) in probands with extreme PRS+ (**Supplementary Figure 30**).

## Discussion

Owing to increasingly accessible GWAS summary statistics and advances in statistical methods to directly model summary-level data, genetic correlation estimation, especially at the genome-wide scale, has become a routine procedure in post-GWAS analyses. These correlation estimates effectively summarize the complex etiologic sharing of multiple traits into concise, robust, and interpretable values, which provided novel insights into the shared genetic architecture of a spectrum of phenotypes. However, genome-wide genetic correlations only reflect the average concordance of genetic effects across the genome and often fail to reveal the local, heterogenous pleiotropic effects, especially when the underlying genetic basis involves multiple etiologic pathways. To this end, methods that partition genetic covariance by functional annotation or local genetic region have achieved some success^11,12^. These methods generally use more sophisticated statistical models and are more adaptive to diverse types of shared genetic architecture. On the downside, it is statistically more challenging to estimate all the parameters in these models using GWAS summary statistics alone. The problem is further exacerbated by technical issues such as strong LD among SNPs in local regions and sample overlap across different GWASs. Due to these challenges, stratified genetic correlation analysis has not been as popular as its genome-wide counterpart.

In this paper, we have introduced SUPERGNOVA, a unified framework for both genome-wide and stratified genetic correlation analysis. Improved upon our previous work^12^, SUPERGNOVA directly addresses the technical challenges in local genetic correlation inference while retaining the statistical optimality in analyses at the genome-wide scale. Through extensive simulations, we demonstrated that SUPERGNOVA provides statistically robust and efficient estimates and substantially outperforms other methods in estimation accuracy and statistical power. Notably, SUPERGNOVA uses GWAS summary statistics as the input and is robust to arbitrary sample overlap between GWAS datasets.

Applied to 30 complex traits, SUPERGNOVA identified 150 trait pairs with significant local genetic covariance, including 86 pairs without a significant global correlation. We identified various patterns in the shared genetic architecture between traits, with some traits (e.g., EA and CP) showing ubiquitous genetic covariance in a large fraction of the genome and other traits (e.g., Crohn’s disease and UC) showing relatively sparse genetic sharing with strong pleiotropic effects. Our analyses also implicated hub regions in the genome that are significantly correlated across numerous neuropsychiatric phenotypes. These results can guide future modeling efforts on these traits as well as functional genomic studies that interrogate key regions with pervasive regulatory roles across many phenotypes.

ASD and cognitive ability showed significant, bidirectional local genetic correlations in our analysis. We performed in-depth analyses to further dissect the shared genetics of ASD and cognition. Human genetic studies for ASD have been fruitful in the past decade. Numerous consortium-scale whole-exome and whole-genome sequencing studies have been conducted to assess the roles of *de novo* mutations and very rare transmitted variants in ASD^27,50,65,66^. These studies have convincingly identified more than 100 risk genes and a number of etiologic pathways for ASD. Additionally, overwhelming evidence suggests that rare and *de novo* pathogenic variants in pathways such as chromatin modifiers and *FMRP* target genes contribute to the comorbidity of ASD and intellectual disability^27^. In contrast, successful GWASs for ASD have just begun to emerge^44^. It was notable that risk genes implicated by common SNPs do not have an apparent overlap with ASD genes identified in rare variant studies. Interestingly, based on GWAS data, researchers have identified a positive genetic correlation between ASD and cognition^16^. A recent study further demonstrated that PRS of EA is over-transmitted from healthy parents to ASD probands, including probands who have pathogenic *de novo* mutations in known ASD genes, but not to the unaffected siblings^45^. These findings raised two important questions. Why are ASD genes affected by common SNPs different from genes harboring rare protein-altering variants? Why do common and rare variants suggest opposite genetic relationships between ASD and cognition?

We aimed to address these questions head-on using local genetic correlations. We identified significant positive correlations of ASD and CP in 18 genomic regions but also 6 regions showing significant negative correlations. Locally, we did not observe the paradoxical correlation pattern seen in the global analysis, i.e., two positively correlated neurodevelopmental disorders ASD and ADHD showing opposite correlations with cognitive measures. Regions that were significantly correlated in all three trait pairs (e.g., the *KMT2E* locus) all showed consistent local correlations between both ASD and ADHD with CP. Of note, the set of regions negatively correlated between ASD and CP had a 3.2-fold enrichment for chromatin modifier genes. Thus, a genetic signature with consistent results between common and rare variants was hidden in plain sight. These genes, affected by both rare protein-altering variants and common (possibly regulatory) SNPs, may contribute to ASD with comorbid intellectual impairment in part through dysregulating chromatin modification in the developing brain. The positive global correlation between ASD and cognition was explained by a second genetic signature driven by a different set of regions that showed positive local correlations and were significantly enriched for PSD genes. When calculating the total genetic covariance between ASD and CP in the genome, negatively correlated regions were overwhelmed by the positive covariance in regions involved in the second signature, thus showing a positive global covariance. PRS based on these two signatures (PRS+ and PRS-) showed distinct associations with ASD phenotypes in the SPARK cohort. Compared to PRS-, PRS+ could better distinguish ASD cases from healthy controls. Both PRS+ and PRS-were associated with IQ in ASD probands but with opposite directions. In addition, PRS-significantly predicted overall ASD symptom severity while PRS+ significantly predicted repetitive behaviors. We also observed an enrichment of Asperger’s disorder in probands with high PRS+ (and a slight depletion in probands with high PRS-) but the results remain to be validated using larger samples in the future.

Our method still has some limitations. Although SUPERGNOVA can effectively estimate local genetic covariance, local genetic correlation estimates are numerically unstable due to the non-negligible noise in the estimates of local heritability. Second, due to the distal regulatory nature of common genetic variations, causal genes may not always be included in the pre-defined genetic region harboring GWAS associations. We suggest researchers also investigate regions adjacent to the identified region when interpreting local correlation results from SUPERGNOVA. Our implemented software allows users to re-define their local region of interest if needed. Third, a future direction is to extend our method to estimate transethnic local genetic correlation^13^. The local correlation estimates provided by SUPERGNOVA may also improve other types of multi-trait analysis such as multi-trait association mapping^3^ and genomic structural equation modeling (GenomicSEM)^4^. We believe SUPERGNOVA may play a critical role in accelerating the development of novel statistical genetics tools in the future.

Taken together, SUPERGNOVA provides a biologically-motivated and statistically principled analytical strategy to tackle etiologic sharing of complex traits. A combination of global and local genetic correlation could provide new insights into the shared genetic basis of many phenotypes. We believe SUPERGNOVA will have wide applications in complex trait genetics research.

## Methods

### Statistical model

We start with the statistical framework for global genetic covariance. Assume there are two studies with sample sizes *n*_1_ and *n*_2_, respectively. Standardized trait values *ϕ*_1_ and *ϕ*_2_ follow the linear models below:

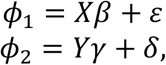

where *X* and *Y* are *n*_1_ × *m* and *n*_2_ × *m* standardized genotype matrices; *m* is the number of shared SNPs between the two studies; *ϵ* and *δ* are the noise terms; and *β* and *γ* denote the genetic effects for *ϕ*_1_ and *ϕ*_2_. We adopt a model with random effects and random design matrices^10,12,24^ to define genetic covariance *ρ*. The combined random vector of *β* and *γ* follows a multivariate normal distribution given by:

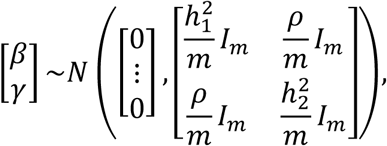

where 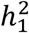 and 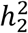 are the heritability of the two traits, respectively; *I*_*m*_ is the identity matrix of size *m*. In practice, two different GWASs may share a subset of samples. Without loss of generality, we assume the first *n*_*s*_ samples in each study are shared (*n*_*s*_ ≤ *n*_1_ and *n*_*s*_ ≤ *n*_2_). The non-genetic effects of the shared samples for the two traits are correlated:

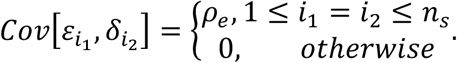

Since trait values *ϕ*_1_ and *ϕ*_2_ are standardized, we have 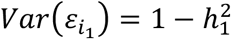 and 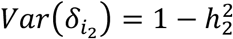 for 1 ≤ *i*_1_ ≤ *n*_1_ and 1 ≤ *i*_2_ ≤ *n*_2_.

In GWAS summary data, we can approximate z-scores of SNP *j* for trait 1 and trait 2 by 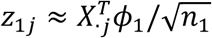 and 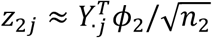. We use *z*_1_ and *z*_2_ to denote the vectors for all SNPs’ z-scores and use *V* to denote the LD matrix. Under a random design model, 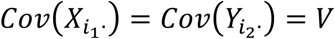 and the variance-covariance matrix of 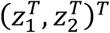 is given by

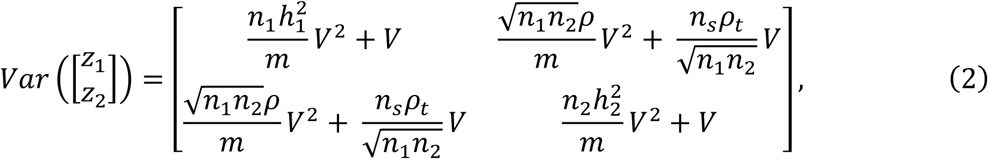

where *ρ*_*t*_ is defined as the sum of genetic covariance and non-genetic effects covariance, i.e. *ρ*_*t*_ = *ρ* + *ρ*_*e*_. We provide detailed derivations of (2) in the **Supplementary Note**.

Most existing genetic covariance methods are based on the idea of minimizing the “distance” between the empirical covariance matrix 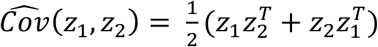 and the theoretical covariance in (1). For example, LDSC^10^ regresses the diagonal elements of empirical z-score covariance matrix on that of the theoretical covariance matrix. GNOVA^12^ applies the method of moments estimator that compares the trace of the empirical and theoretical covariance matrices. Our new approach, SUPERGNOVA, is also based on this unified framework.

The statistical framework we introduced above can be easily generalized to local genetic covariance. We assume *ϕ*_1_ and *ϕ*_2_ follow additive linear models:

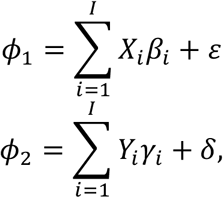

where *X*_*i*_ and *Y*_*i*_ are the genotypes and *β*_*i*_ and *γ*_*i*_ are the effect sizes of SNPs in region *i*. In practice, *I* genomic regions can be mutually independent LD blocks defined by LDetect^29^. Following the same derivations as shown above, the variance-covariance matrix of local z scores *z*_1*i*_ and *z*_2*i*_ is

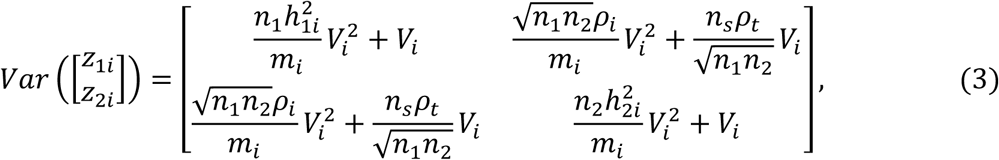

where 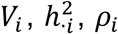 and *m*_*i*_ are LD matrix, heritability, genetic covariance and number of SNPs for region *i*, respectively. *ρ*_*t*_ here is defined as the sum of local genetic covariance and non-genetic effects covariance, i.e. 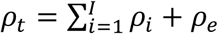. Details about the construction of statistical model for local genetic covariance are provided in **Supplementary Note**.

### Local genetic covariance estimation

Following (3), the covariance of *z*_1*i*_ and *z*_2*i*_ (i.e. z-scores of trait 1 and trait 2 in region *i*) is

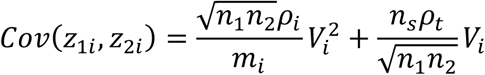

Assume eigen decomposition of *V*_*i*_ is 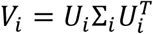, then we have

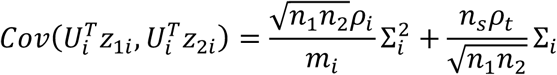

where 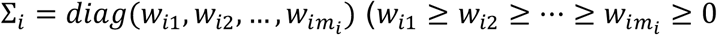 are the eigenvalues of Σ_*i*_) and *U*_*i*_ is the corresponding orthogonal matrix of eigenvectors. Denote 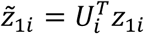 and 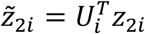. For *j* = 1, 2, …, *m*_*i*_, the expected value and variance of 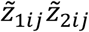 for the *j*th eigenvalue *w*_*ij*_ are

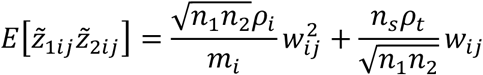

and

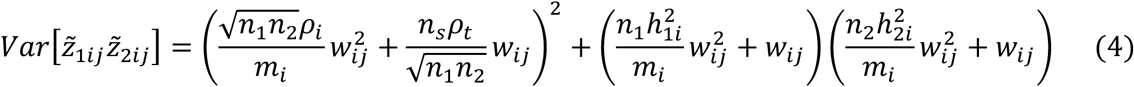

where 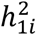 and 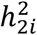 can be estimated by the method of moments^67^. Derivations of (3) and (4) are in the **Supplementary Note**. Due to the noise in LD estimation, we only use the first *K*_*i*_ eigenvalues to estimate *ρ*_*i*_. The procedure to adaptively determine *K*_*i*_ is described in the following section. In practice, the LD matrices are estimated from an external reference panel (e.g., the 1000 Genomes Project^28^) and the intercept of cross-trait LDSC^10^ provides an estimate of 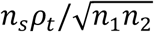, denoted as 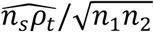. For each genomic region, we can estimate local genetic covariance and test the significance of 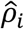 using the weighted regression of 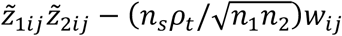, denoted by *η*_*ij*_, on the square of eigenvalue weighted by the reciprocal of the variance in (4) which is approximated by 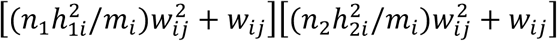. Since 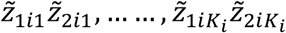are independent for any region *i*, the theoretical variance of 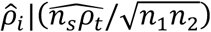 is analytically given by

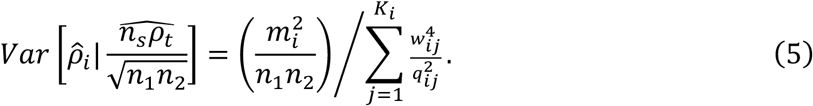

Here, we denote 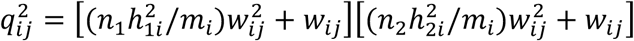. In weighted regression, 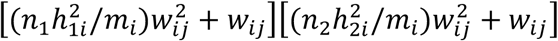 is treated as the reciprocal of the weight’s square. So, the empirical variance of 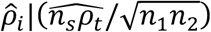 is analytically given by

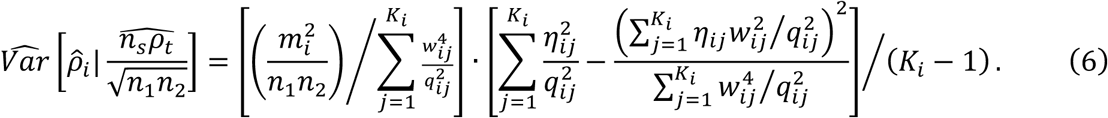

Derivations of (5) and (6) are in the **Supplementary Note.** To compensate for the variance introduced by LDSC in the estimation of 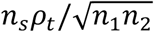, we approximate 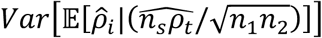 by

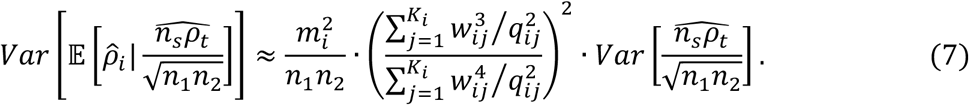

The derivation of (7) is in the **Supplementary Note**. The estimation of the last term in (7) 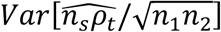 is from LDSC. We use (6) to approximate 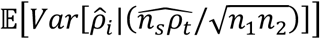. By the law of total variance, we combine the results in (6) and (7) to obtain *Var*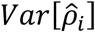:

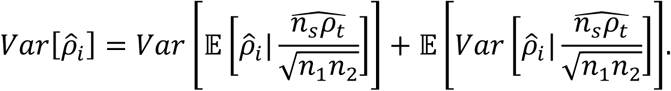

Then, local genetic correlation is estimated by 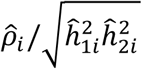. We approximate the standard error and the confidence interval of local genetic correlation estimation by the Delta method. However, local genetic correlation estimates might be numerically unstable due to the noise in the estimates of local heritability, which is in the denominator of the estimator of local genetic correlation. So, the estimates of genetic covariance of SUPERGNOVA are more reliable.

### An adaptive procedure to determine *K*_*i*_

Sample LD information is rarely available for published GWASs. Therefore, we use an external reference panel to estimate LD. In practice, the number of SNPs is far greater than the number of individuals in the reference panel. For example, in this paper, we used 503 samples of European ancestry from the 1000 Genomes Project phase III as the reference panel. The average number of SNPs in a local region is about 2,000. To achieve robust inference, we apply factor selection and only use the first *K*_*i*_ eigenvectors and eigenvalues for region *i*. There are several existing methods to perform factor selection^68-73^. Here, we propose an adaptive procedure to determine the value of optimal *K*_*i*_. Under the optimal *K*_*i*_, theoretical variance in (5) and empirical variance in (6) should be close. We know from (5) that theoretical variance decreases with the increase of *K*_*i*_. However, the value of empirical variance rapidly increases when the cutoff for the eigenvalues approaches towards zero (**Supplementary Figure 31**). We adaptively determine the optimal *K*_*i*_ as follows. First, we set an upper bound for *K*_*i*_. In our paper, the upper bound is 503 which is the number of samples in the reference panel. Then, for region *i*, we compute the value of theoretical variance and empirical variance for *K*_*i*_ taking values from 10 to the upper bound. We denote the maximum of theoretical variance and empirical variance for each *K*_*i*_ as

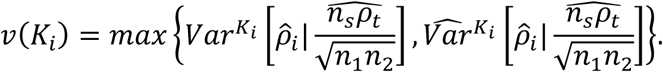

The optimal *K*_*i*_ is determined by 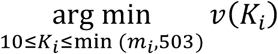

### Simulation settings

We used genotype data from the Wellcome Trust Case Control Consortium (WTCCC) to conduct simulations. Samples were randomly divided into two equal subgroups with 7,959 individuals. We denote them as set 1 and set 2. We randomly sampled 3,979 individuals from set 1 and 3,980 individuals from set 2 to create set 3 which has a 50% sample overlap with set 1. Samples with European ancestry from the 1000 Genomes Project phase III^28^ were used as the LD reference in our simulations. We kept common SNPs with MAFs greater than 5% and removed all SNPs with ambiguous alleles. After quality control, 287,539 SNPs remained in both WTCCC and 1000 Genomes Project data.

We used LDetect^29^ to partition the genome into 2,197 LD blocks (∼1.6 centimorgan in width on average). To estimate local genetic covariance, we selected the largest region partitioned by LDetect on chromosome 2 (176,998,822-180,334,969) to be the local region of interest. There are 395 SNPs in this region in the genotype data from WTCCC. The effects of SNPs on two simulated traits were only correlated in the local region. The remaining 23,839 SNPs on chromosome 2 were used as background SNPs whose genetic covariance was set as 0. The effect sizes of SNPs were generated by a multivariate normal distribution and we applied Genome-wide Complex Trait Analysis (GCTA)^74^ to simulate *ϕ*_1_ and *ϕ*_2_. We used PLINK^75^ to run GWAS and obtain summary statistics of the two simulated traits. We repeated each simulation setting 100 times. Detailed simulation settings are summarized below.

For simulations of global genetic covariance analysis, we set the heritability of two traits to be 0.5 and set the genetic covariance to be 0, 0.05, 0.1, 0.15, 0.2, and 0.25, respectively. We conducted three simulations, corresponding to different levels of sample overlap.

1. Independent studies: we used set 1 and set 2 to simulate *ϕ*_1_ and *ϕ*_2_, respectively.
2. Partial sample overlap: *ϕ*_1_ and *ϕ*_2_ were simulated on set 1 and set 3. The covariance of non-genetic effects on shared samples was set to be 0.2.
3. Complete sample overlap: *ϕ*_1_ and *ϕ*_2_ were both simulated on set 1. The covariance of non-genetic effects on shared samples was set to be 0.2.

For simulations of local genetic covariance analysis, we set the heritability of two traits to be 0.5. The total heritability is evenly distributed to all SNPs. The covariance of the local genetic effects was set to be 0, 0.001, 0.002, 0.003, 0.004, and 0.005, respectively. Similar to global analyses, we conducted three sets of simulations.

1. Independent studies: *ϕ*_1_ and *ϕ*_2_ were simulated on set 1 and set 2, respectively.
2. Partial sample overlap: we simulated *ϕ*_1_ and *ϕ*_2_ on set 1 and set 3. The covariance of non-genetic effects was set to be 0.2.
3. Complete sample overlap: we simulated both *ϕ*_1_ and *ϕ*_2_ on set 1. The covariance of non-genetic effects was set to be 0.2.

To evaluate the robustness of *ρ*-HESS^11^ against mis-specified overlapping sample size, we provided the method with an overlapping sample size of 1,000 and a phenotypic correlation of 0.05 as input in partial sample overlap and complete sample overlap scenarios to estimate local genetic covariance.

### GWAS data

GWAS summary statistics of 29 complex traits included in our analyses are publicly available. We obtained the summary statistics of a recent lung cancer GWAS directly from the authors^76^. Details of the 30 GWASs are summarized in **Supplementary Table 1**. We used munge_sumstats.py script in LDSC to reformat these data and removed strand-ambiguous SNPs from each dataset. For each trait pair, we took the intersection of SNPs in two GWAS and the 1000 Genomes Project. We matched the effect alleles after removing SNPs with MAF lower than 5%. We only included the SNPs in autosomes and excluded the MHC region in all analyses.

We accessed samples from the SPARK study through the Simons Foundation Autism Research Initiative (SFARI). Samples in the SPARK study were genotyped by the Illumina Infinium Global Screening Array. Details on these samples have been previously reported and are available on the SFARI website^77^. Following data processing procedure in Huang et al.^54^, we performed pre-imputation quality control (QC) using PLINK. The genotype data were phased and imputed to the HRC reference panel version r1.1 2016 using the Michigan Imputation server^78^.

### Estimation of the proportion of correlated regions

We estimated the proportion of correlated regions with an R package called *ashr*^*30*^ after the estimation of local genetic covariance among the 30 phenotypes. The inputs were estimates of local genetic covariance and its standard error. The unimodal prior distribution was set to be “halfnormal” for all the results of pairs of traits. The method applied a Bayesian framework to compute FDR for each genomic region. To estimate the numbers of correlated regions for each pair of traits, we computed the sum of (1 – FDR) given by ashr for each region.

### Follow-up analyses in the *SPI1* locus for AD and other neuropsychiatric traits

To replicate local genetic covariance identified at the *SPI1* locus, we defined a new genomic region centered at *SPI1* with a 1-Mb span. We estimated the local genetic covariance between AD (IGAP2019^41^) and the other 29 traits for this region. For replication, we implemented a GWAS for AD family history in the UK Biobank and estimated the local genetic covariance of this GWAS with other traits. Details on the AD-proxy GWAS have been previously reported^41,79^.

We obtained PU.1 binding sites as ChIP-seq peaks from the ReMap datasets^80^ (GEO: GSE31621; *SPI1*, blood monocyte and macrophage datasets^81^). Following Huang et al.^43^, we expanded each ChIP-seq peak by 150 kb up- and downstream to define the transcription factor binding site annotation. We applied GNOVA^12^ to estimate the genetic covariance between AD and 29 other traits in the PU.1 binding sites. We trained elastic net gene expression imputation models^82,83^ using expression profiles adjusted by peer factors^84^ and PCs and matched genotypes from 758 monocyte and 599 macrophage samples in the Cardiogenics Consortium^85^ imputed by Michigan Imputation Server^78^. We downloaded Cardiogenics resources from European Genome-phenome Archive (EGA) platform. To investigate the regulatory relationship between PU.1 and the identified genes in myeloid cells, we used GREAT^86^ to map PU.1 each binding peak in macrophages and monocytes to the nearest gene.

### Cross-tissue transcriptome-wide association analysis

To identify genes associated with ADHD, ASD, CP and SCZ in brain tissues in the *KMT2E* region (chr7: 104158491-105425027), we implemented cross-tissue transcriptome-wide association analysis using UTMOST^87^. We used gene expression imputation models trained by genotype and normalized gene expression data from the GTEx project^88-91^ (version V8). We considered 13 brain tissues. For individual expression data, we regressed out the effects of confounding covariates including first five genotype PCs, PEER factors optimized by sample sizes as in the GTEx V8 paper^91^, sequencing platforms, library construction protocol and donor sex. Cis-genotype data was extracted for SNPs located within 1MB distance from the transcription starting sites of all protein coding genes. Then we trained expression imputation models based on cis-genotypes for each gene in each tissue using 10-fold elastic net with alpha being 0.5. Models with credible imputation performances (FDR<0.05) were used in later analysis.

### Functional annotation for variants in GWAS data

We used bedtools^92^ to extract sequence from the *KMT2E* region. We then performed gene annotations on each of the variants using ANNOVAR^93^. For exonic and splicing variants, missense variants were represented by nonsynonymous single nucleotide variants (SNVs) and loss-of-function variants were annotated as frameshift, stopgain, or stoploss mutations by ANNOVAR. We took overlapped SNPs and matched the effect alleles between ANNOVAR annotations and GWAS summary data of ADHD, ASD, CP and SCZ respectively.

### Gene set enrichment analysis

We used R package *TxDb.Hsapiens.UCSC.hg19.knownGene* to identify genes in the correlated regions between ASD and CP with nominal significant covariances (p<0.01). We only included protein-coding genes in our analysis, resulting in 317 positively correlated genes and 179 negatively correlated genes. We applied *Enrichr*^*94,95*^ to implement enrichment analysis on GWAS catalog 2019^96^ (**Supplementary Tables 18-19**), and TF PPI^94^. We identified *FMRP* target genes, genes encoding PSD proteins, gene preferentially expressed in human embryonic brains, essential genes, chromatin modifier genes, genes with probability of loss-of-function intolerance (pLI) > 0.9, and SFARI evidence score based on previous literature^54^. We obtained a list of 102 genes identified by the refined transmitted and *de novo* association (TADA) model^97^ (FDR<0.1) in the recent exome sequencing study on ASD^50^. We performed hypergeometric test to assess the enrichment of ASD-CP positively and negatively correlated genes in these gene sets.

### Analysis of spatio-temporal RNA-seq data in brain tissues

We used single-cell RNA-seq data generated by the PsychENCODE Consortium^98^ in fetal brains to test the elevation of gene expression of ASD-CP correlated genes in brain development. There were 762 cells collected from neocortical regions of eight fetal brains from 5 to 20 PCW. We kept only protein-coding genes which included 18,134 genes in this analysis. 310 of positively correlated and 175 of negatively correlated genes were overlapped with these genes. Following Satterstrom et al.^50^, for each time point, a gene was considered expressed if at least one transcript mapped to this gene in 25% or more of cells for at least one PCW period before. By definition, gene expression rate increased with fetal development. We performed log-rank test to test the difference of gene expression rate in developmental brain between positively or negatively correlated genes and other genes.

We downloaded developmental bulk RNA-seq data from BrainSpan. Gene-level RPKMs were used across 524 samples from 42 individuals in 26 brain regions^98^. We kept protein-coding genes in our analysis. Following Satterstrom et al.^50^, we removed samples with RNA integrity number (RIN) ≤ 7 and only used neocortical regions – dorsolateral prefrontal cortex (DFC), ventrolateral prefrontal cortex (VFC), medial prefrontal cortex (MFC), orbitofrontal cortex (OFC), primary motor cortex (M1C), primary somatosensory cortex (S1C), primary association cortex (A1C), inferior parietal cortex (IPC), superior temporal cortex (STC), inferior temporal cortex (ITC), and primary visual cortex (V1C). Genes were defined as expressed if their RPKMs were at least 0.5 in 80% samples from at least one neocortical region at one major temporal epoch. Consequently, 14,803 genes were defined as expressed in 325 samples from 8 post-conceptual weeks (PCW) to 40 years of age. We then log-transformed RKPM (log_2_[RKPM+1]). We followed the definition of developmental stages in Li et al^98^. We performed t-test to determine the differential expression among ASD-CP positively correlated genes, negatively correlated genes, and background genes.

To study the relative prenatal and postnatal bias, we performed linear regression for the transformed RKPM of each gene on a binary ‘prenatal’ stage variable. Sex was included as an adjustment variable. Genes were defined as prenatally (or postnatally) biased if log_2_ fold change> 0.1 (or <-0.1) and q-value<0.05 resulting in 5,562 prenatally biased genes and 5,361 postnatally biased genes. We followed the definition of ASD-CP positively and negatively correlated genes from gene set enrichment analysis. Chi-squared test was performed to test if the distributions of prenatally and postnatally biased genes in ASD-CP positively and negatively correlated regions were significantly different from background genes.

### PRS analysis in SPARK

We used the 18 positively correlated regions and 6 negatively corelated regions (FDR<0.1) between ASD and CP to construct PRS+ and PRS- of ASD. We clumped the SNPs by PLINK^75^. We set the significance threshold for index SNPs as 1, LD threshold for clumping as 0.1, and physical distance threshold for clumping as 250 kb. We generated scores for 5,469 ASD probands and 2,132 healthy siblings in the SPARK cohort. We assessed associations between two PRSs and ASD using logistic regression. We then investigated the association between the two PRSs and ASD phenotypes in probands, including IQ, SCQ score, RBS-R score, DCDQ score, and subtypes of ASD. For each phenotype, we used the maximum sample with both genotype and phenotype data. Sample sizes for these phenotypes in SPARK are summarized in **Supplementary Table 25**. We performed two-sample t-test for quantitative phenotypes between probands with extreme PRS (top 1%) and other probands. We performed hypergeometric test to test enrichment of subtypes in the extreme PRS group.

### URLs

LDetect (https://bitbucket.org/nygcresearch/ldetect/src/master/);

LDSC (https://github.com/bulik/ldsc);

GNOVA (https://github.com/xtonyjiang/GNOVA);

*ρ*-HESS (https://huwenboshi.github.io/hess/);

ReMap database (http://pedagogix-tagc.univ-mrs.fr/remap/);

Cardiogenics dataset (https://ega-archive.org/studies/EGAS00001000411);

Enrichr (https://amp.pharm.mssm.edu/Enrichr/);

BrainSpan (http://brainspan.org/static/home);

PsychENCODE (http://www.psychencode.org);

SPARK (https://www.sfari.org/resource/spark/);

### Data and code availability

SUPERGNOVA software is publicly available at https://github.com/qlu-lab/SUPERGNOVA.

## Acknowledgements

This study was Supported in part by NIH grants 3P30AG021342-16S2 and 1R01GM122078 and NSF grants DMS 1713120 and DMS 1902903. Q.L is supported by the Clinical and Translational Science Award (CTSA) program, through the NIH National Center for Advancing Translational Sciences (NCATS), grant UL1TR000427. We also acknowledge research support from the University of Wisconsin-Madison Office of the Chancellor and the Vice Chancellor for Research and Graduate Education with funding from the Wisconsin Alumni Research Foundation and the Waisman Center pilot grant program at the University of Wisconsin-Madison.

This study makes use of data generated by the Wellcome Trust Case-Control Consortium. We conducted the research using the UK Biobank resource under approved data requests (refs: 29900 and 42148). This study makes use of summary statistics from many GWAS consortia. We thank the investigators in these GWAS consortia for generously sharing their data. We thank the CARDIOGENICS project for providing expression data of macrophages and monocytes and expression data. CARDIOGENICS resources was funded by the European Union FP6 program (LSHM-CT-2006-037593). We thank the PsychENCODE Consortium for providing the single-cell RNA-seq data in fetal brain. We are grateful to all the families participating in the Simons Foundation Powering Autism Research for Knowledge (SPARK) study.

## Author contributions

Y.Z. and Q.L. developed the statistical framework.

Y.Z. performed statistical analysis.

Y.Y. assisted in analyzing GWAS data.

K.H. curated gene list of ASD pathways.

Y.W and K.H. processed SPARK data.

W.L. and Z.Y. assisted in training expression imputation model.

Y.W and X.Z implemented the GWAS on AD family history.

B.L. performed ANNOVAR analysis.

B.T., D.W., and J.L. advised on the biology of ASD and ADHD.

Q.L. and H.Z. advised on statistical and genetics issues.

Y.Z. implemented the software.

Y.Z. and Q.L. wrote the manuscript.

All authors contributed to manuscript editing and approved the manuscript.

## Notes

https://github.com/qlu-lab/SUPERGNOVA

### Competing Interest Statement

The authors have declared no competing interest.

