## Supplementary Note for "Local genetic correlation analysis reveals heterogeneous etiologic sharing of complex traits"

### 1 Properties for the SUPERGNOVA framework

#### 1.1 Statistical model for global genetic covariance

We follow the random design and random effects model to construct phenotypes. Suppose we sample two cohorts of different phenotypes, in which sample sizes are $n_1$ and $n_2$, respectively. Assume the two GWAS studies share the same set of SNPs and there are $m$ SNPs in total. We measure phenotype 1 in cohort 1 and phenotype 2 in cohort 2. We assume all the $m$ SNPs are associated with both traits. We model phenotype vectors for each cohort as

$$\begin{aligned}\phi_1 &= X\beta + \epsilon \\ \phi_2 &= Y\gamma + \delta,\end{aligned}$$

where $X$ and $Y$ are standardized random matrices of genotypes, with demensions $n_1 \times m$ and $n_2 \times m$; $\beta$ and $\gamma$ are vectors of standardized genotype effect sizes, and $\delta$ and $\epsilon$ are vectors of residuals, representing environmental effects and non-additive genetic effects.

Each row of $X$ and $Y$ represents the standardized genotypes of an individual in the corresponding GWAS study. By standardized genotypes, we mean the genotype of each SNP is normalized to mean zero and variance one. We assume the genotypes of different samples are independent from each other. Due to linkage disequilibrium(LD), genotypes of different SNPs are correlated. We denote the LD matrix as $V$. That is, $cov(X_{i_1\cdot}) = V = cov(Y_{i_2\cdot})$, for any $1 \leq i_1 \leq n_1$ and $1 \leq i_2 \leq n_2$. We define the LD score of a variant $j$ as $l_j := \sum_k V_{jk}^2$. We assume we can bound $l_j$ by generic constant $M$, i.e. $M > l_j$, for $j \in \{1, 2, \ldots, p\}$. We suppose that $(\beta^T, \gamma^T)^T$ is subject to multivariate normal distribution and has mean zero and covariance matrix

$$\mathrm{Var}\left[\begin{pmatrix}\beta \\ \gamma\end{pmatrix}\right] = \frac{1}{m}\begin{pmatrix} h_1^2 I_m & \rho I_m \\ \rho I_m & h_2^2 I_m\end{pmatrix}.$$

We define $h_1^2$ and $h_2^2$ as the heritability of trait 1 and trait 2, respectively. $\rho$ is defined as genetic covariance between trait 1 and trait 2. In addition, genetic correlation $r$ is defined by $\rho/\sqrt{h_1^2 h_2^2}$.

In practice, two different GWASs often share a subset of samples. Without loss of generality, we assume $0 \leq n_s \leq \min\{n_1, n_2\}$ is the number of samples shared by the two GWASs and the first $n_s$ samples in each study are shared, i.e. the first $n_s$ rows of $X$ and $Y$ are the same. To account for the non-genetic correlation introduced by sample overlapping, we assume $\epsilon$ and $\delta$ are subject to multivariate normal distribution with covariance:

$$\mathrm{Cov}\left[\epsilon_i, \delta_j\right] = \begin{cases}\rho_e, 1 \leq i = j \leq n_s \\ 0, \text{otherwise}\end{cases}.$$

The variance of $\epsilon$ and $\delta$ are:

$$\mathrm{Var}\left[\epsilon\right] = \left(1 - h_1^2\right) I_{n_1},\ \mathrm{Var}\left[\delta\right] = \left(1 - h_2^2\right) I_{n_2},$$

so that

$$\begin{aligned}\text{Var}\left[\phi_1\right] &= \text{Var}\left[X\beta\right] + \text{Var}[\epsilon] \\ &= \mathbb{E}\left[X\beta\beta^T X^T\right] + \left(1-h_1^2\right) I_{n_1} \\ &= \frac{h_1^2}{m}\mathbb{E}\left[XX^T\right] + \left(1-h_1^2\right) I_{n_1} \\ &= h_1^2 I_{n_1} + \left(1-h_1^2\right) I_{n_1} = I_{n_1}\end{aligned}$$

and similarly, $\text{Var}\left[\phi_2\right] = I_{n_2}$. We assume genotype, effect size, and environmental effect are independent to each other.

Genetic covariance $\rho$ is the covariance of genetic components. For an individual with standarized genotype $G$, denoting $g_1$ and $g_2$ as the genetic components for trait 1 and trait 2, respectively, we have:

$$\begin{aligned}\text{Cov}\left[g_1, g_2\right] &= \mathbb{E}\left[G^T\beta\gamma^T G\right] \\ &= \mathbb{E}\left[\mathbb{E}\left[G^T\beta\gamma^T G \middle| G\right]\right] \\ &= \mathbb{E}\left[G^T\mathbb{E}\left[\beta\gamma^T\right] G\right] \\ &= \frac{\rho}{m}\mathbb{E}\left[G^T G\right] \\ &= \rho.\end{aligned}$$

We define genetic correlation as the genetic covariance normalized by the heritability:

$$r_g = \frac{\text{Cov}\left[g_1, g_2\right]}{\sqrt{\text{Var}\left[g_1\right]\text{Var}\left[g_2\right]}} = \frac{\rho}{\sqrt{h_1^2 h_2^2}}.$$

### 1.2 Statistical model for local genetic covariance

In this section, we generalize the statistical framework above for local genetic covariance. We assume $\phi_1$ and $\phi_2$ follow additive linear models:

$$\begin{aligned}\phi_1 &= \sum_{i=1}^{I} X_i\beta_i + \epsilon \\ \phi_2 &= \sum_{i=1}^{I} Y_i\gamma_i + \delta,\end{aligned}$$

where $X_i$ and $Y_i$ are the standarized genotypes and $\beta_i$ and $\gamma_i$ are the effect sizes of SNPs in regions $i$. We assume SNPs from different regions are independent and we use $V_i$ to denote the LD matrix in region $i$. We assume effect size $(\beta_i^T, \gamma_i^T)^T$ is subject to multivariate normal distribution:

$$\begin{pmatrix}\beta_i \\ \gamma_i\end{pmatrix} \sim N\left(0, \frac{1}{m_i}\begin{bmatrix} h_{1i}^2 I_{m_i} & \rho_i I_{m_i} \\ \rho_i I_{m_i} & h_{2i}^2 I_{m_i}\end{bmatrix}\right)$$

and effect sizes of SNPs from different regions are independent. Environmental error covariance $\rho_e$ in local genetic covariance is defined the same as it is in global genetic covariance. The variance of $\epsilon$ and $\delta$ are $Var\left[\epsilon\right] = \left(1 - \sum_{i=1}^{I} h_{1i}^2\right) I_{n_1}$and $Var\left[\delta\right] = \left(1 - \sum_{i=1}^{I} h_{2i}^2\right) I_{n_2}$ so that $Var\left(\phi_1\right) = 1$ and $Var\left(\phi_2\right) = 1$.

The covariance of the genetic components is the sum of local genetic covariance. Denote $G_i$

as the standardized genotype in region $i$ for an individual. We have

$$\begin{aligned}\operatorname{Cov}[g_1, g_2] &= \mathbb{E}\left[\left(\sum_{i=1}^{I} G_i^T \beta_i\right)\left(\sum_{i=1}^{I} \gamma_i^T G_i\right)\right] \\ &= \sum_{i=1}^{I} \mathbb{E}\left[\left(G_i^T \beta_i \gamma_i^T G_i\right)\right] \\ &= \sum_{i=1}^{I} \mathbb{E}\left[\mathbb{E}\left[G_i^T \beta_i \gamma_i^T G_i \middle| G_i\right]\right] \\ &= \sum_{i=1}^{I} \mathbb{E}\left[G_i^T \mathbb{E}\left[\beta_i \gamma_i^T\right] G_i\right] \\ &= \sum_{i=1}^{I} \frac{\rho_i}{m_i} \mathbb{E}\left[G_i^T G_i\right] \\ &= \sum_{i=1}^{I} \rho_i.\end{aligned}$$

Local genetic correlation is defined by:

$$r_{ig} = \frac{\rho_i}{\sqrt{h_{1i}^2 h_{2i}^2}}.$$

### 1.3 Covariance of z scores

In genome-wide association studies (GWAS), summary statistics are more accessible than individual-level genotype data due to potential privacy and data sharing security concerns. For a GWAS with quantitative trait, the z score of a single SNP $j$ is given by:

$$z_j = \frac{\hat{\beta}_j}{se(\hat{\beta}_j)}$$

where $\hat{\beta}_j$ is the estimated coefficient from marginal linear regression between the trait and the SNP $j$, and $se(\hat{\beta}_j)$ is the corresponding standard error. In practice, we approximate z scores by $z_{1j} = X_{\cdot j}^T \phi_1 / \sqrt{n_1}$ and $z_{2j} = Y_{\cdot j}^T \phi_2 / \sqrt{n_2}$.

#### 1.3.1 Global

We derive the variance-covariance matrix of $\left(z_1^T, z_2^T\right)^T$ in this section. It's sufficient to show that

$$Cov\left(z_1, z_2\right) = \frac{\sqrt{n_1 n_2}\rho}{m} V^2 + \frac{n_s \rho_t}{\sqrt{n_1 n_2}} V, \tag{1}$$

where $\rho_t = \rho + \rho_e$. Because $Var(z_1)$ and $Var(z_2)$ can be derived from (1). In fact, to compute $Var(z_1)$, we can assume trait 1 and trait 2 are from the same study and hence we have

$$Var(z_1) = Cov(z_1, z_1) = \frac{\sqrt{n_1 n_1} h_1^2}{m} V^2 + \frac{n_1\left[h_1^2 + \left(1 - h_1^2\right)\right]}{\sqrt{n_1 n_1}} V = \frac{n_1 h_1^2}{m} V^2 + V.$$

Similarly, $Cov(z_2) = \left(n_2 h_2^2 / m\right) V^2 + V$. To prove (1), we begin with the following proposition.

**Proposition 1** *Assume* $\Gamma_1, \Gamma_2 \overset{i.i.d.}{\sim} B(1, p_\Gamma)$; $\Pi_1, \Pi_2 \overset{i.i.d.}{\sim} B(1, p_\Pi)$; $Cor(\Gamma_1, \Pi_1) = Cor(\Gamma_2, \Pi_2) = r_{\Gamma\Pi}$; $\Gamma_1$ *and* $\Pi_2$ *are independent.* $\Gamma_2$ *and* $\Pi_1$ *are independent. We have:*

*(1).*

$$
\begin{aligned}
&\mathbb{E}\left[\left(\Gamma_1+\Gamma_2-2p_\Gamma\right)^2\left(\Pi_1+\Pi_2-2p_\Pi\right)^2\right]\\
=&4\left(r_{\Gamma\Pi}^2+1\right)p_\Pi\left(1-p_\Pi\right)p_\Gamma\left(1-p_\Gamma\right)+2r_{\Gamma\Pi}\sqrt{p_\Pi\left(1-p_\Pi\right)}\sqrt{p_\Gamma\left(1-p_\Gamma\right)}\left(1-2p_\Pi\right)\left(1-2p_\Gamma\right).
\end{aligned}
$$

*(2). Additionally, if* $\Lambda_1,\Lambda_2 \overset{i.i.d.}{\sim} B(1,p_\Lambda)$*;* $Cor(\Gamma_1,\Lambda_1)=Cor(\Gamma_2,\Lambda_2)=r_{\Gamma\Lambda}$*;* $Cor(\Pi_1,\Lambda_1)=Cor(\Pi_2,\Lambda_2)=r_{\Pi\Lambda}$*;* $\mathbb{E}\left[\left(\Pi_1-p_\Pi\right)\left(\Gamma_1-p_\Gamma\right)\left(\Lambda_1-p_\Lambda\right)\right]=\mathbb{E}\left[\left(\Pi_2-p_\Pi\right)\left(\Gamma_2-p_\Gamma\right)\left(\Lambda_2-p_\Lambda\right)\right]=\Delta$*;* $\Lambda_1$ *is independent with* $\Gamma_2$ *and* $\Pi_2$*;* $\Lambda_2$ *is independent with* $\Gamma_1$ *and* $\Pi_1$*. We have:*

$$
\begin{aligned}
&\mathbb{E}\left[\left(\Gamma_1+\Gamma_2-2p_\Gamma\right)\left(\Lambda_1+\Lambda_2-2p_\Lambda\right)\left(\Pi_1+\Pi_2-2p_\Pi\right)^2\right]\\
=&4\left(r_{\Gamma\Pi}r_{\Pi\Lambda}+r_{\Gamma\Lambda}\right)p_\Pi\left(1-p_\Pi\right)\sqrt{p_\Gamma\left(1-p_\Gamma\right)}\sqrt{p_\Lambda\left(1-p_\Lambda\right)}+2\left(1-2p_\Pi\right)\Delta.
\end{aligned}
$$

We note the genotype of SNP $j$ before standardization is the sum of two independent idential binomial distribution with $p$ equal to minor allele frequency, $0.05 < f_j < 0.5$, of SNP $j$. We only consider common variants. With Hardy-Weinberg equilibrium, for any $1 \le \tau \le n_1$, we approximate $X_{\tau j}$ by

$$X_{\tau j} := \frac{S_{1\tau j}+S_{2\tau j}-2f_j}{\sqrt{2f_j\left(1-f_j\right)}},$$

where $S_{1\tau j}$ and $S_{2\tau j}$ represent allelic dosage for SNP $j$ on each chromosomethe and $f_j$ is the minor allele frequency of SNP $j$. The approximate of genotype in $Y$ is defined in the same way. By proposition 1, for different SNP, SNP $j$, SNP $\zeta$, we have

$$\mathbb{E}\left[X_{\tau j}^2 X_{\tau\zeta}^2\right]=V_{j\zeta}^2+1+\frac{V_{j\zeta}\left(1-2f_j\right)\left(1-2f_\zeta\right)}{2\sqrt{f_j\left(1-f_j\right)}\sqrt{f_\zeta\left(1-f_\zeta\right)}}.$$

For SNP $k$, $k \neq j,\zeta$, denoting $\Delta_{jk\zeta}=\mathbb{E}\left[\left(S_{\alpha\tau j}-f_j\right)\left(S_{\alpha\tau k}-f_k\right)\left(S_{\alpha\tau\zeta}-f_\zeta\right)\right]$, $\alpha=1,2$, we have

$$\mathbb{E}\left[X_{\tau j}X_{\tau k}X_{\tau\zeta}^2\right]=V_{j\zeta}V_{k\zeta}+V_{jk}+\frac{\left(1-2f_\zeta\right)\Delta_{jk\zeta}}{2f_\zeta\left(1-f_\zeta\right)\sqrt{f_j\left(1-f_j\right)}\sqrt{f_k\left(1-f_k\right)}},$$

where $V_{j\zeta}$ represents the $j$-th row, $\zeta$-th column of LD matrix $V$. In other words, $V_{j\zeta}$ is the correlation of genotypes between SNP $j$ and SNP $\zeta$. $V_{jk}$ and $V_{k\zeta}$ are defined in the same way.

Now, we prove (1). We have

$$
\begin{aligned}
\mathbb{E}\left[z_1z_2^T\right]&=\mathbb{E}\left[\frac{X^T\phi_1}{\sqrt{n_1}}\cdot\frac{\phi_2^TY}{\sqrt{n_2}}\right]=\mathbb{E}\left[\frac{X^T\left(X\beta+\epsilon\right)}{\sqrt{n_1}}\cdot\frac{\left(Y\gamma+\delta\right)^TY}{\sqrt{n_2}}\right]\\
&=\mathbb{E}\left[\frac{X^TX\beta\gamma^TY^TY}{\sqrt{n_1n_2}}\right]+\mathbb{E}\left[\frac{X^T\epsilon\delta^TY}{\sqrt{n_1n_2}}\right]\\
&=\mathbb{E}\left\{\mathbb{E}\left[\frac{X^TX\beta\gamma^TY^TY}{\sqrt{n_1n_2}}\middle|X,Y\right]\right\}+\mathbb{E}\left\{\mathbb{E}\left[\frac{X^T\epsilon\delta^TY}{\sqrt{n_1n_2}}\middle|X,Y\right]\right\}\\
&=\mathbb{E}\left\{\frac{X^TX\mathbb{E}\left[\beta\gamma^T\right]Y^TY}{\sqrt{n_1n_2}}\right\}+\mathbb{E}\left\{\frac{X^T\mathbb{E}\left[\epsilon\delta^T\right]Y}{\sqrt{n_1n_2}}\right\}\\
&=\frac{\rho}{m\sqrt{n_1n_2}}\mathbb{E}\left[X^TXY^TY\right]+\frac{\rho_e}{\sqrt{n_1n_2}}\mathbb{E}\left[\sum_{i=1}^{n_s}X_{i\cdot}^TY_{i\cdot}\right]\\
&=\frac{\rho}{m\sqrt{n_1n_2}}\mathbb{E}\left[X^TXY^TY\right]+\frac{\rho_en_s}{\sqrt{n_1n_2}}V. \qquad (2)
\end{aligned}
$$

To complete the calculation, we next compute $\mathbb{E}\left[X_{\cdot j}^T X Y^T Y_{\cdot k}\right]$. First, we denote $\mathcal{N} = \left\{(\tau,\eta) \,\middle|\, \tau=\eta=1,2,\ldots,n_s\right\}$. For $j=k$, we have

$$\begin{aligned}
\mathbb{E}\left[X_{\cdot j}^T X Y^T Y_{\cdot j}\right] &= \mathbb{E}\left[\sum_{\zeta=1}^{p}\left(\sum_{\tau=1}^{n_1} X_{\tau j}X_{\tau\zeta}\right)\cdot\left(\sum_{\eta=1}^{n_2} Y_{\eta j}Y_{\eta\zeta}\right)\right] \\
&= \sum_{\zeta=1}^{p}\sum_{\tau=1}^{n_1}\sum_{\eta=1}^{n_2}\mathbb{E}\left[X_{\tau j}X_{\tau\zeta}Y_{\eta j}Y_{\eta\zeta}\right] \\
&= \sum_{\zeta=1}^{p}\left(\sum_{(\tau,\eta)\in\mathcal{N}}\mathbb{E}\left[X_{\tau j}X_{\tau\zeta}Y_{\eta j}Y_{\eta\zeta}\right] + \sum_{(\tau,\eta)\notin\mathcal{N}}\mathbb{E}\left[X_{\tau j}X_{\tau\zeta}\right]\mathbb{E}\left[Y_{\eta j}Y_{\eta\zeta}\right]\right) \\
&= \sum_{\zeta=1}^{p}\left(\sum_{(\tau,\eta)\in\mathcal{N}}\mathbb{E}\left[X_{\tau j}^2 X_{\tau\zeta}^2\right] + (n_1n_2-n_s)\,V_{j\zeta}^2\right) \\
&= \sum_{\zeta=1}^{p}\left(n_s\left[V_{j\zeta}^2 + 1 + \frac{V_{j\zeta}(1-2f_j)(1-2f_\zeta)}{2\sqrt{f_j(1-f_j)}\sqrt{f_\zeta(1-f_\zeta)}}\right] + (n_1n_2-n_s)\,V_{j\zeta}^2\right) \\
&= n_1n_2\sum_{\zeta=1}^{p}V_{j\zeta}^2 + \sum_{\zeta=1}^{p}n_s + \sum_{\zeta=1}^{p}\frac{n_sV_{j\zeta}(1-2f_j)(1-2f_\zeta)}{2\sqrt{f_j(1-f_j)}\sqrt{f_\zeta(1-f_\zeta)}} \\
&= n_1n_2\sum_{\zeta=1}^{p}V_{j\zeta}^2 + pn_s + n_s\sum_{\zeta=1}^{p}\frac{V_{j\zeta}(1-2f_j)(1-2f_\zeta)}{2\sqrt{f_j(1-f_j)}\sqrt{f_\zeta(1-f_\zeta)}} \\
&= n_1n_2\sum_{\zeta=1}^{p}V_{j\zeta}^2 + pn_s + o(n_sp) \\
&\approx n_1n_2\sum_{\zeta=1}^{p}V_{j\zeta}^2 + pn_s. \qquad (3)
\end{aligned}$$

We explain the approximation in (3) here. In the deivation below, $C$ represents a generic constant, whose value may be different at different places. As mentioned above, we only consider common variants in our supplementary note, which means $0.05 < f_j < 0.5$ for any SNP $j$. So, for any SNP $j$ and $\zeta$, there exists a constant $C$ so that

$$\frac{(1-2f_j)(1-2f_\zeta)}{2\sqrt{f_j(1-f_j)}\sqrt{f_\zeta(1-f_\zeta)}} < C.$$

By Cauchy-Schwarz inequality, we have

$$\sum_{\zeta=1}^{p}\frac{V_{j\zeta}(1-2f_j)(1-2f_\zeta)}{2\sqrt{f_j(1-f_j)}\sqrt{f_\zeta(1-f_\zeta)}} < C\sum_{\zeta=1}^{p}V_{j\zeta} \le C\sqrt{p\sum_{\zeta=1}^{p}V_{j\zeta}^2} = C\sqrt{pl_j}. \qquad (4)$$

So, we have

$$n_s\sum_{\zeta=1}^{p}\frac{V_{j\zeta}(1-2f_j)(1-2f_\zeta)}{2\sqrt{f_j(1-f_j)}\sqrt{f_\zeta(1-f_\zeta)}} \le C\cdot l_j\cdot n_s\sqrt{p} = o(n_sp)$$

For $j \neq k$, we have

$$\begin{aligned}
\mathbb{E}\left[X_{\cdot j}^T X Y^T Y_{\cdot k}\right] &= \mathbb{E}\left[\sum_{\zeta=1}^{p}\left(\sum_{\tau=1}^{n_1} X_{\tau j} X_{\tau\zeta}\right)\cdot\left(\sum_{\eta=1}^{n_2} Y_{\eta k} Y_{\eta\zeta}\right)\right] \\
&= \sum_{\zeta=1}^{p}\sum_{\tau=1}^{n_1}\sum_{\eta=1}^{n_2}\mathbb{E}\left[X_{\tau j} X_{\tau\zeta} Y_{\eta k} Y_{\eta\zeta}\right] \\
&= \sum_{\zeta=1}^{p}\left(\sum_{(\tau,\eta)\in\mathcal{N}}\mathbb{E}\left[X_{\tau j} X_{\tau\zeta} Y_{\eta k} Y_{\eta\zeta}\right] + \sum_{(\tau,\eta)\notin\mathcal{N}}\mathbb{E}\left[X_{\tau j} X_{\tau\zeta}\right]\mathbb{E}\left[Y_{\eta k} Y_{\eta\zeta}\right]\right) \\
&= \sum_{\zeta=1}^{p}\left(\sum_{(\tau,\eta)\in\mathcal{N}}\mathbb{E}\left[X_{\tau j} X_{\tau k} X_{\tau\zeta}^2\right] + \left(n_1 n_2 - n_s\right) V_{j\zeta} V_{k\zeta}\right) \\
&= \sum_{\zeta=1}^{p}\left(n_s\left[V_{j\zeta} V_{k\zeta} + V_{jk} + \frac{\left(1-2f_\zeta\right)\Delta_{jk\zeta}}{2f_\zeta\left(1-f_\zeta\right)\sqrt{f_j\left(1-f_j\right)}\sqrt{f_k\left(1-f_k\right)}}\right] + \left(n_1 n_2 - n_s\right) V_{j\zeta} V_{k\zeta}\right) \\
&= n_1 n_2 \sum_{\zeta=1}^{p} V_{j\zeta} V_{k\zeta} + \sum_{\zeta=1}^{p} n_s V_{jk} + \sum_{\zeta=1}^{p}\frac{n_s\left(1-2f_\zeta\right)\Delta_{jk\zeta}}{2f_\zeta\left(1-f_\zeta\right)\sqrt{f_j\left(1-f_j\right)}\sqrt{f_k\left(1-f_k\right)}} \\
&= n_1 n_2 \sum_{\zeta=1}^{p} V_{j\zeta} V_{k\zeta} + p n_s V_{jk} + n_s\sum_{\zeta=1}^{p}\frac{\left(1-2f_\zeta\right)\Delta_{jk\zeta}}{2f_\zeta\left(1-f_\zeta\right)\sqrt{f_j\left(1-f_j\right)}\sqrt{f_k\left(1-f_k\right)}} \\
&= n_1 n_2 \sum_{\zeta=1}^{p} V_{j\zeta} V_{k\zeta} + p n_s V_{jk} + o(n_s p) \\
&\approx n_1 n_2 \sum_{\zeta=1}^{p} V_{j\zeta} V_{k\zeta} + p n_s V_{jk}. \qquad (5)
\end{aligned}$$

We note that $\Delta_{jk\zeta} \leq V_{j\zeta}$ by definition. So, the approximation in (5) is derived from the same argument as (4).

Combine (3) and (5) we have

$$\mathbb{E}\left[X^T X Y^T Y\right] \approx n_1 n_2 V^2 + p n_s V \qquad (6)$$

Substituting (6) into (2), we finish the calculation for $Cov\left(z_1, z_2\right)$.

#### 1.3.2 Local

We assume SNPs from different local regions are indepedent. In other words, we assume LD matrix takes the block structure

$$V = \begin{pmatrix} V_1 & & & \\ & V_2 & & \\ & & \ddots & \\ & & & V_I \end{pmatrix},$$

where $I$ is the number of locla regions.

Based on the derivations for global genetic covariance above, we generalize the results to calculate covariance of local z scores, $Cov(z_{1i}, z_{2i})$. We have

$$
\begin{aligned}
\mathbb{E}\left[z_{1i}z_{2i}^T\right] &= \mathbb{E}\left[\frac{X_i^T\phi_1}{\sqrt{n_1}}\cdot\frac{\phi_2^TY_i}{\sqrt{n_2}}\right] = \mathbb{E}\left[\frac{X_i^T\left(\sum_{\nu=1}^I X_\nu\beta_\nu+\epsilon\right)}{\sqrt{n_1}}\cdot\frac{\left(\sum_{\nu=1}^I Y_\nu\gamma_\nu+\delta\right)^T Y_i}{\sqrt{n_2}}\right] \\
&= \sum_{\nu=1}^{I}\mathbb{E}\left[\frac{X_i^TX_\nu\beta_\nu\gamma_\nu^TY_\nu^TY_i}{\sqrt{n_1n_2}}\right]+\mathbb{E}\left[\frac{X_i^T\epsilon\delta^TY_i}{\sqrt{n_1n_2}}\right] \\
&= \mathbb{E}\left[\frac{X_i^TX_i\beta_i\gamma_i^TY_i^TY_i}{\sqrt{n_1n_2}}\right]+\sum_{\nu\neq i}\mathbb{E}\left[\frac{X_i^TX_\nu\beta_\nu\gamma_\nu^TY_\nu^TY_i}{\sqrt{n_1n_2}}\right]+\frac{n_s\rho_e}{\sqrt{n_1n_2}}V_i \\
&= \frac{\rho_i}{m_i}\mathbb{E}\left[\frac{X_i^TX_iY_i^TY_i}{\sqrt{n_1n_2}}\right]+\sum_{\nu\neq i}\frac{\rho_\nu}{m_\nu}\mathbb{E}\left[\frac{X_i^T\mathbb{E}\left(X_\nu Y_\nu^T\right)Y_i}{\sqrt{n_1n_2}}\right]+\frac{n_s\rho_e}{\sqrt{n_1n_2}}V_i \\
&\approx \frac{\sqrt{n_1n_2}\rho_i}{m_i}V_i^2+\frac{n_s\rho_i}{\sqrt{n_1n_2}}V_i+\sum_{\nu\neq i}\frac{n_s\rho_\nu}{\sqrt{n_1n_2}}V_i+\frac{n_s\rho_e}{\sqrt{n_1n_2}}V_i \qquad (7)\\
&= \frac{\sqrt{n_1n_2}\rho_i}{m_i}V_i^2+\frac{n_s\rho_t}{\sqrt{n_1n_2}}V_i.
\end{aligned}
$$

Here, $\rho_t = \sum_{i=1}^{I}\rho_i + \rho_e$. Approximation in (7) follows from (6).

#### 1.4 Variance of $\tilde{z}_{1ij}\tilde{z}_{2ij}$

Assume eigen decomposition of $V_i$ is $V_i = U_i\Sigma_iU_i^T$. Denote $\tilde{z}_{1i} = U_i^Tz_{1i}$ and $\tilde{z}_{2i} = U_i^Tz_{2i}$. Then, we have

$$
\begin{pmatrix}\tilde{z}_{1i}\\ \tilde{z}_{2i}\end{pmatrix} \sim N\left(0, \begin{pmatrix}\frac{n_1h_{1i}^2}{m_i}\Sigma_i^2+\Sigma_i & \frac{\sqrt{n_1n_2}\rho_i}{m_i}\Sigma_i^2+\frac{n_s\rho_t}{\sqrt{n_1n_2}}\Sigma_i\\ \frac{\sqrt{n_1n_2}\rho_i}{m_i}\Sigma_i^2+\frac{n_s\rho_t}{\sqrt{n_1n_2}}\Sigma_i & \frac{n_2h_{2i}^2}{m_i}\Sigma_i^2+\Sigma_i\end{pmatrix}\right). \qquad (8)
$$

We order the eigenvalues in $\Sigma_i$ by their values. We denote the eigenvalues by $w_{i1} \leq w_{i2} \leq \ldots \leq w_{im_i}$. $\tilde{z}_{1ij}$ and $\tilde{z}_{2ij}$ denote the $j$-th element of $\tilde{z}_{1i}$ and $\tilde{z}_{2i}$, respectively. We use the following proposition to obtain $Var\left(\tilde{z}_{1ij}\tilde{z}_{2ij}\right)$.

**Proposition 2** *Assume*

$$
\begin{pmatrix}\xi_1\\ \xi_2\end{pmatrix} \sim N\left(0, \begin{pmatrix}\sigma_1^2 & \rho_0\\ \rho_0 & \sigma_2^2\end{pmatrix}\right)
$$

*we have* $Var\left(\xi_1\xi_2\right) = \rho_0^2 + \sigma_1^2\sigma_2^2$.

By proposition 2, we have

$$
var\left[\tilde{z}_{1ij}\tilde{z}_{2ij}\right] = \left(\frac{\sqrt{n_1n_2}\rho_i}{m_i}w_{ij}^2+\frac{n_s\rho_t}{\sqrt{n_1n_2}}w_{ij}\right)^2+\left(\frac{n_1h_{1i}^2}{m_i}w_{ij}^2+w_{ij}\right)\left(\frac{n_2h_{2i}^2}{m_i}w_{ij}^2+w_{ij}\right). \qquad (9)
$$

#### 1.5 Variance of $\hat{\rho}_i$

From (8), we have

$$
\mathbb{E}\left[\tilde{z}_{1ij}\tilde{z}_{2ij}\right] = \frac{\sqrt{n_1n_2}\rho_i}{m_i}w_{ij}^2+\frac{n_s\rho_t}{\sqrt{n_1n_2}}w_{ij}.
$$

We obtain the estimates of $\rho_i$ by weighted linear regression that regresses $\tilde{z}_{1ij}\tilde{z}_{2ij} - \frac{n_s\rho_t}{\sqrt{n_1n_2}}w_{ij}$ on $w_{ij}^2$. We use LD score regression[1] to obtain the estimates of $\frac{n_s\rho_t}{\sqrt{n_1n_2}}$, which is denoted by $\widehat{n_s\rho_t}/\sqrt{n_1n_2}$. The weights are given by the reciprocal of the variance in (9). In practice, we use $\left(\frac{n_1h_{1i}^2}{m_i}w_{ij}^2+w_{ij}\right)\left(\frac{n_2h_{2i}^2}{m_i}w_{ij}^2+w_{ij}\right)$ to approximate the variance, $\mathbb{E}\left[\tilde{z}_{1ij}\tilde{z}_{2ij}\right]$. For notation simplicity, we denote $q_{ij}^2 = \left(\frac{n_1 h_{1i}^2}{m_i} w_{ij}^2 + w_{ij}\right)\left(\frac{n_2 h_{2i}^2}{m_i} w_{ij}^2 + w_{ij}\right)$ and $\eta_{ij} = \tilde{z}_{1ij}\tilde{z}_{2ij} - \frac{n_s \rho_t}{\sqrt{n_1 n_2}} w_{ij}$. We note $Var\left(n_{ij}\right) = q_{ij}^2$. We use the $K_i$ largest eigenvalues in the estimation of local genetic covariance to reduce noise.

#### 1.5.1 Theoretical and empirical variance of $\hat{\rho}_i$

From weighted linear regression, the weighted estimate of local genetic covariance is given by

$$\hat{\rho}_i = \frac{m_i}{\sqrt{n_1 n_2}} \frac{\sum_{j=1}^{K_i} \eta_{ij} w_{ij}^2 / q_{ij}^2}{\sum_{j=1}^{K_i} w_{ij}^4 / q_{ij}^2}. \tag{10}$$

So, the theoretical variance of $\hat{\rho}_i$ is given by

$$Var\left(\hat{\rho}_i\right) = \left(\frac{m_i^2}{n_1 n_2}\right) Var\left(\frac{\sum_{j=1}^{K_i} \eta_{ij} w_{ij}^2 / q_{ij}^2}{\sum_{j=1}^{K_i} w_{ij}^4 / q_{ij}^2}\right) = \left(\frac{m_i^2}{n_1 n_2}\right) \frac{\sum_{j=1}^{K_i} Var\left(\eta_{ij}\right) w_{ij}^4 / q_{ij}^4}{\left(\sum_{j=1}^{K_i} w_{ij}^4 / q_{ij}^2\right)^2} = \frac{m_i^2 / \left(n_1 n_2\right)}{\sum_{j=1}^{K_i} w_{ij}^4 / q_{ij}^2}.$$

We assume the variance of $\eta_{ij}$ is proportional to $q_{ij}$ and given by $q_{ij}\sigma_{\eta i}^2$ although we know the true value of $\sigma_{\eta i}^2$ is equal to 1. In other words, we treat the variance of $\eta_{ij}$ as unknown and use weighted linear regression procedure to estimate $\sigma_{\eta i}^2$. The estimate of $\sigma_{\eta i}^2$ is denoted by $\hat{\sigma}_{\eta i}^2$. Then, multiply the theoretical variance by $\hat{\sigma}_{\eta i}^2$, we have the empirical variance of $\hat{\rho}_i$

$$\widehat{Var}\left(\hat{\rho}_i\right) = Var\left(\hat{\rho}_i\right) \hat{\sigma}_{\eta i}^2. \tag{11}$$

We obtain $\hat{\sigma}_{\eta i}^2$ by the residues from the regression model. We have

$$\hat{\sigma}_{\eta i}^2 = \left[\sum_{j=1}^{K_i} \frac{\eta_{ij}^2}{q_{ij}^2} - \frac{\left(\sum_{j=1}^{K_i} \eta_{ij} w_{ij}^2 / q_{ij}^2\right)^2}{\sum_{j=1}^{K_i} w_{ij}^4 / q_{ij}^2}\right] \Bigg/ \left(K_i - 1\right). \tag{12}$$

Substituting (12) into (11), we have

$$\widehat{Var}\left(\hat{\rho}_i\right) = \frac{m_i^2 / \left(n_1 n_2\right)}{\sum_{j=1}^{K_i} w_{ij}^4 / q_{ij}^2} \cdot \left[\sum_{j=1}^{K_i} \frac{\eta_{ij}^2}{q_{ij}^2} - \frac{\left(\sum_{j=1}^{K_i} \eta_{ij} w_{ij}^2 / q_{ij}^2\right)^2}{\sum_{j=1}^{K_i} w_{ij}^4 / q_{ij}^2}\right] \Bigg/ \left(K_i - 1\right). \tag{13}$$

#### 1.5.2 Approximation of $\mathbf{Var}\left(\mathbb{E}\left[\hat{\rho}_i \middle| \left(\widehat{\mathbf{n_s}\rho_\mathbf{t}} / \sqrt{\mathbf{n_1 n_2}}\right)\right]\right)$

To compensate the noise introduced by the estimation of $\widehat{n_s\rho_t}/\sqrt{n_1 n_2}$ in LD score regression[1], we add $Var\left(\mathbb{E}\left[\hat{\rho}_i \middle| \left(\widehat{n_s\rho_t}/\sqrt{n_1 n_2}\right)\right]\right)$ to the total variance, $Var\left(\hat{\rho}_i\right)$. Then, the total variance of $\hat{\rho}_i$ is then given by

$$Var\left(\hat{\rho}_i\right) = Var\left(\mathbb{E}\left[\hat{\rho}_i \middle| \frac{\widehat{n_s\rho_t}}{\sqrt{n_1 n_2}}\right]\right) + \mathbb{E}\left(Var\left[\hat{\rho}_i \middle| \frac{\widehat{n_s\rho_t}}{\sqrt{n_1 n_2}}\right]\right).$$

The estimation of $\mathbb{E}\left(Var\left[\hat{\rho}_i|\left(\widehat{n_s\rho_t}/\sqrt{n_1n_2}\right)\right]\right)$ is given by the empirical variance in (13). In this section, we derive the expression of $Var\left(\mathbb{E}\left[\hat{\rho}_i|\left(\widehat{n_s\rho_t}/\sqrt{n_1n_2}\right)\right]\right)$. By (10), We have

$$\begin{aligned}
\mathbb{E}\left[\hat{\rho}_i\left|\frac{\widehat{n_s\rho_t}}{\sqrt{n_1n_2}}\right.\right] &= \frac{m_i}{\sqrt{n_1n_2}}\mathbb{E}\left[\frac{\sum_{j=1}^{K_i}\eta_{ij}w_{ij}^2/q_{ij}^2}{\sum_{j=1}^{K_i}w_{ij}^4/q_{ij}^2}\left|\frac{\widehat{n_s\rho_t}}{\sqrt{n_1n_2}}\right.\right] \\
&= \frac{m_i}{\sqrt{n_1n_2}}\frac{\sum_{j=1}^{K_i}\mathbb{E}\left[\eta_{ij}|\left(\widehat{n_s\rho_t}/\sqrt{n_1n_2}\right)\right]w_{ij}^2/q_{ij}^2}{\sum_{j=1}^{K_i}w_{ij}^4/q_{ij}^2} \\
&= \frac{m_i}{\sqrt{n_1n_2}}\frac{\sum_{j=1}^{K_i}\left(\mathbb{E}\left[\tilde{z}_{1ij}\tilde{z}_{2ij}\right]-\frac{\widehat{n_s\rho_t}}{\sqrt{n_1n_2}}w_{ij}\right)w_{ij}^2/q_{ij}^2}{\sum_{j=1}^{K_i}w_{ij}^4/q_{ij}^2} \\
&= \frac{m_i}{\sqrt{n_1n_2}}\frac{\sum_{j=1}^{K_i}\mathbb{E}\left[\tilde{z}_{1ij}\tilde{z}_{2ij}\right]w_{ij}^2/q_{ij}^2}{\sum_{j=1}^{K_i}w_{ij}^4/q_{ij}^2} - \frac{m_i}{\sqrt{n_1n_2}}\frac{\widehat{n_s\rho_t}}{\sqrt{n_1n_2}}\frac{\sum_{j=1}^{K_i}w_{ij}^3/q_{ij}^2}{\sum_{j=1}^{K_i}w_{ij}^4/q_{ij}^2}. \qquad (14)
\end{aligned}$$

The first term in (14) is constant. So, $Var\left(\mathbb{E}\left[\hat{\rho}_i|\left(\widehat{n_s\rho_t}/\sqrt{n_1n_2}\right)\right]\right)$ is given by

$$\begin{aligned}
Var\left(\mathbb{E}\left[\hat{\rho}_i\left|\frac{\widehat{n_s\rho_t}}{\sqrt{n_1n_2}}\right.\right]\right) &= Var\left(\frac{m_i}{\sqrt{n_1n_2}}\frac{\widehat{n_s\rho_t}}{\sqrt{n_1n_2}}\frac{\sum_{j=1}^{K_i}w_{ij}^3/q_{ij}^2}{\sum_{j=1}^{K_i}w_{ij}^4/q_{ij}^2}\right) \\
&= \frac{m_i^2}{n_1n_2}\left(\frac{\sum_{j=1}^{K_i}w_{ij}^3/q_{ij}^2}{\sum_{j=1}^{K_i}w_{ij}^4/q_{ij}^2}\right)^2 Var\left(\frac{\widehat{n_s\rho_t}}{\sqrt{n_1n_2}}\right).
\end{aligned}$$

$Var\left(\widehat{n_s\rho_t}/\sqrt{n_1n_2}\right)$ is the variance of the intercept estimated by LD score regression.

### 2 Consistency of SUPERGNOVA and GNOVA

The estimates of SUPERGNOVA is approximately consistent with the estimates of GNOVA[2] when the number of SNPs $m$ is large enough. The estimates of GNOVA corrected for sample overlap is given by

$$\hat{\rho}_{gnova} = \frac{m\left(\sqrt{n_1n_2}z_1^Tz_2 - mn_s\rho_t\right)}{n_1n_2\sum_{j=1}^{m}l_j}. \qquad (15)$$

Without loss of generality, we demonstrate the equivalence between SUPERGNOVA and GNOVA for global genetic covariance estimation and we use all the eigenvalues of LD matrix. By (10), the SUPERGNOVA estimator for global genetic covariance is given by

$$\hat{\rho} = \frac{m}{\sqrt{n_1n_2}}\frac{\sum_{j=1}^{m}\eta_jw_j^2/q_j^2}{\sum_{j=1}^{m}w_j^4/q_j^2},$$

with $q_j$, $w_j$ and $\eta_j$ are defined in the obvious way. We assume the number of SNPs is large enough so that we can approximate $q_j^2$ by $w_j^2$. So, the SUPERGNOVA estimator can be written as:

$$\begin{aligned}
\hat{\rho} &= \frac{m}{\sqrt{n_1 n_2}} \frac{\sum_{j=1}^{m}\left(\tilde{z}_{1j}\tilde{z}_{2j} - \frac{n_s \rho_t}{\sqrt{n_1 n_2}} w_j\right)}{\sum_{j=1}^{m} w_j^2} \\
&= \frac{m}{\sqrt{n_1 n_2}} \frac{\tilde{z}_1^T \tilde{z}_2 - \sum_{j=1}^{m} \frac{n_s \rho_t}{\sqrt{n_1 n_2}} w_j}{\sum_{j=1}^{m} w_j^2} \\
&= \frac{m}{\sqrt{n_1 n_2}} \frac{z_1^T U U^T z_2 - \frac{n_s \rho_t}{\sqrt{n_1 n_2}} tr\left(V\right)}{tr\left(V^2\right)} \\
&= \frac{m}{\sqrt{n_1 n_2}} \frac{z_1^T z_2 - \frac{m n_s \rho_t}{\sqrt{n_1 n_2}}}{\sum_{j=1}^{m} l_j} \\
&= \frac{m\left(\sqrt{n_1 n_2} z_1^T z_2 - m n_s \rho_t\right)}{n_1 n_2 \sum_{j=1}^{m} l_j},
\end{aligned}$$

which is exactly the GNOVA estimator given in (15).
