## Supplementary Figures for "Local genetic correlation analysis reveals heterogeneous etiologic sharing of complex traits"

**Supplementary Figure 1. Evaluation of global covariance estimation and type I error control on non-overlapping datasets (set 1 and set 2). (A)** Comparison of global genetic covariance estimates among LDSC, GNOVA and SUPERGNOVA **(B)** qq-plot of p value when the true global genetic covariance is set to be 0.

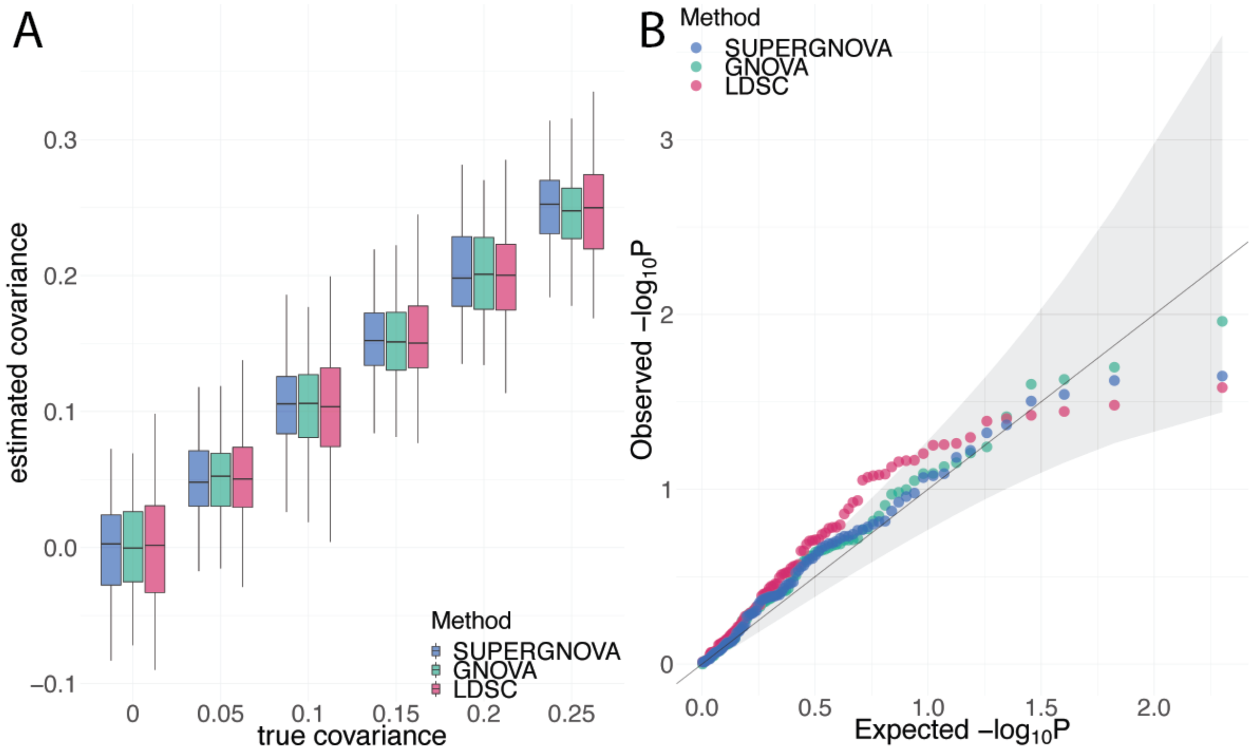

**Supplementary Figure 2. Evaluation of global covariance estimation and type I error control on partially-overlapping datasets (set 1 and set 3). (A)** Comparison of global genetic covariance estimates among LDSC, GNOVA and SUPERGNOVA **(B)** qq-plot of p value when the true global genetic covariance is set to be 0.

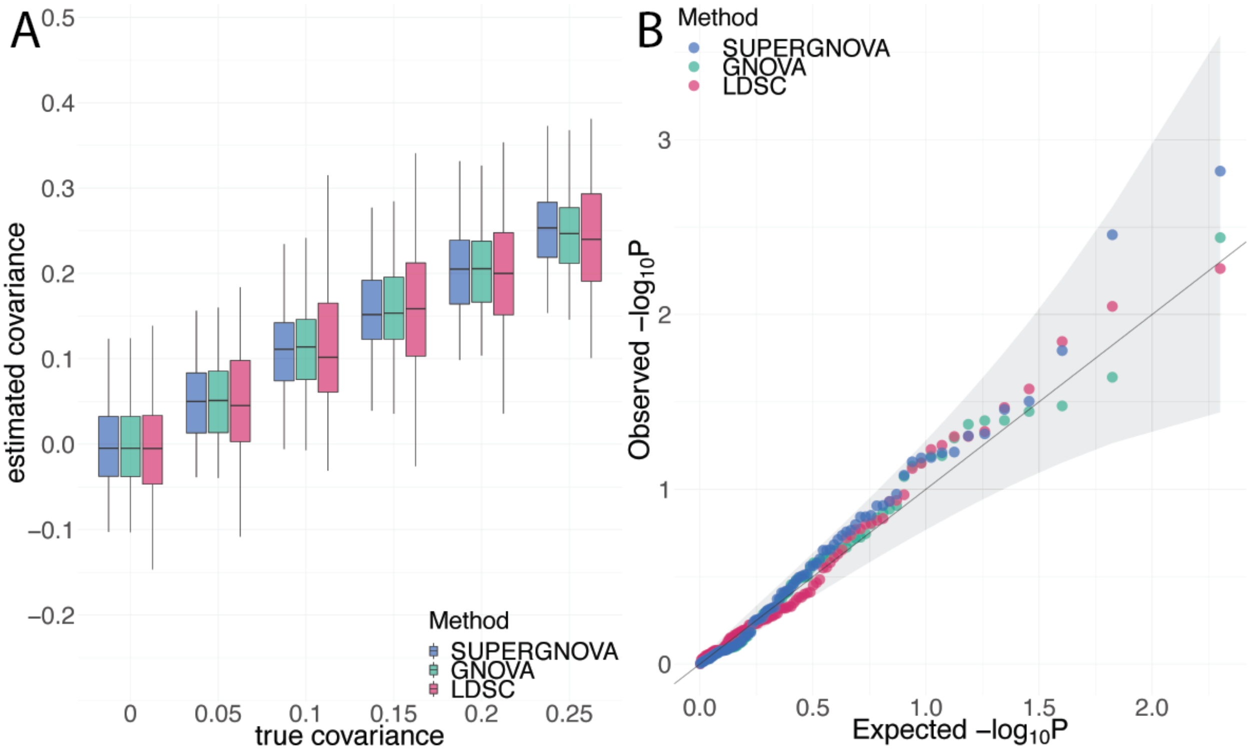

**Supplementary Figure 3. Evaluation of global covariance estimation and type I error control on a completely-overlapping dataset (set 1). (A)** Comparison of global genetic covariance estimates among LDSC, GNOVA and SUPERGNOVA **(B)** qq-plot of p value when the true global genetic covariance is set to be 0.

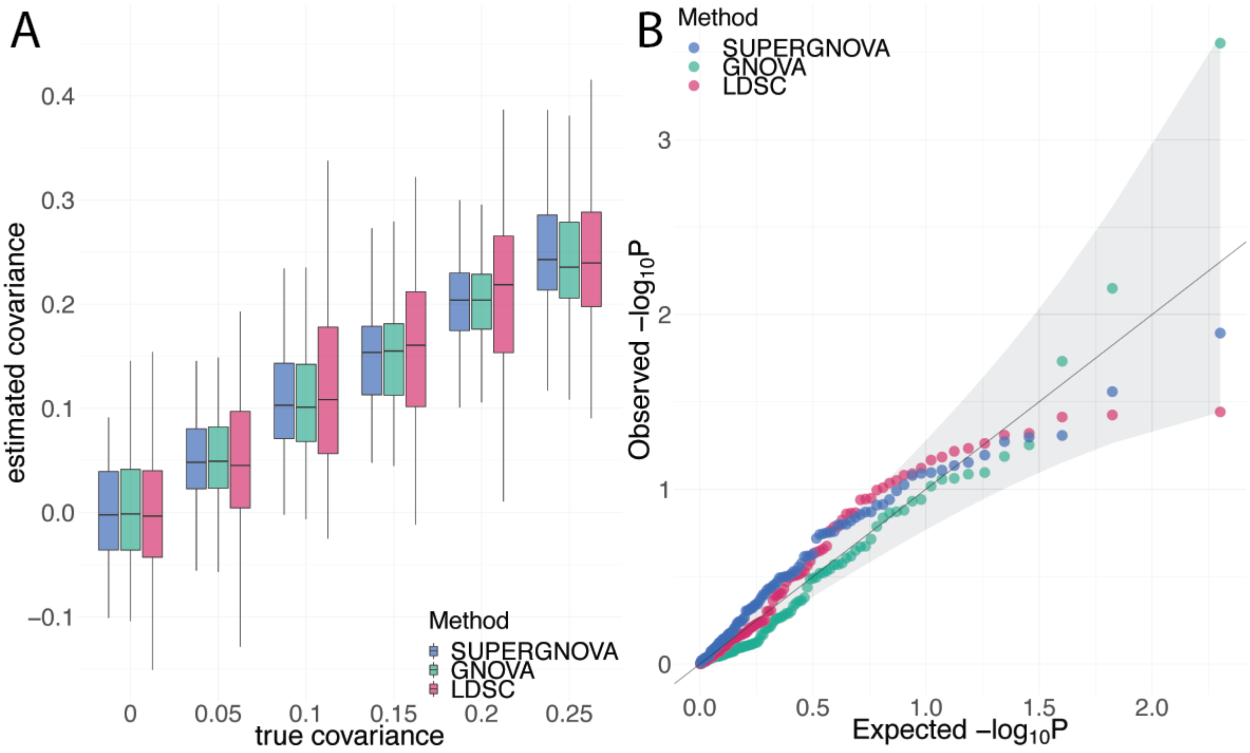

**Supplementary Figure 4. Evaluation of local covariance estimation and type I error control on non-overlapping datasets (set 1 and set 2). (A)** Comparison of local genetic covariance estimates between SUPERGNOVA and $\rho$-HESS. **(B)** qq-plot of p value when the true local genetic covariance is set to be 0.

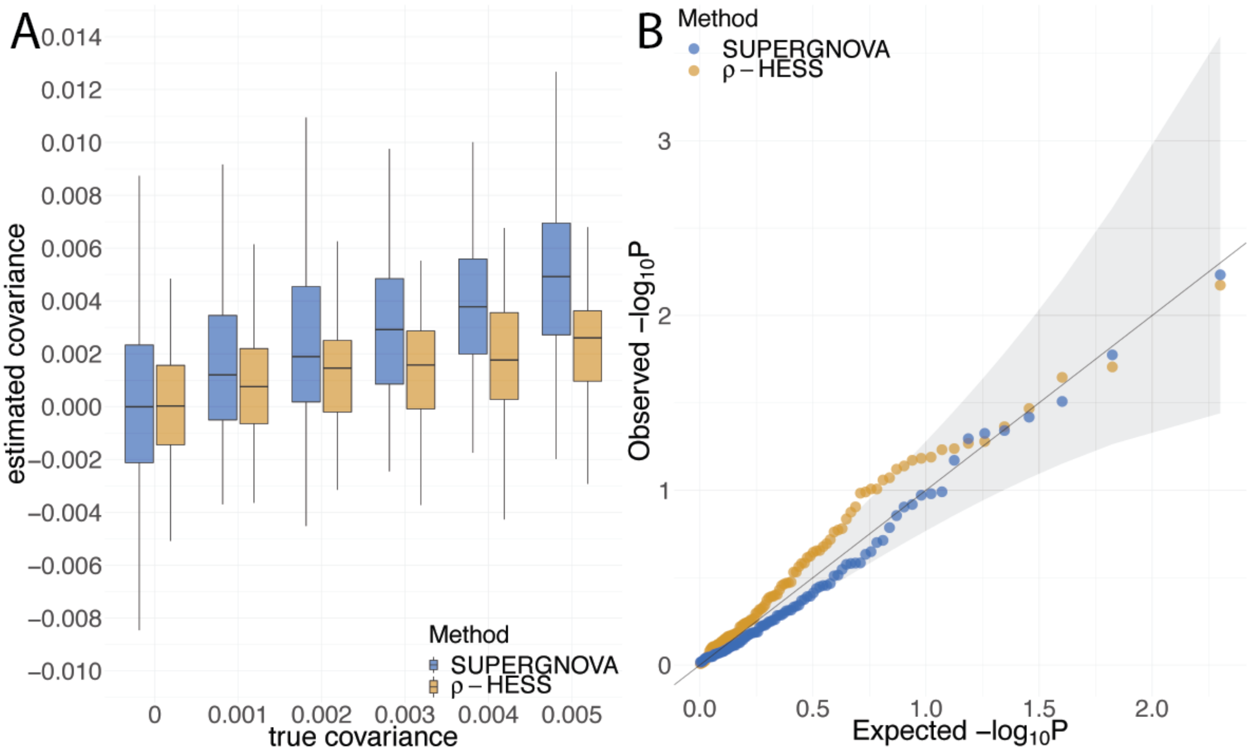

**Supplementary Figure 5. Evaluation of local covariance estimation and type I error control on partially-overlapping datasets (set 1 and set 3). (A)** Comparison of local genetic covariance estimates between SUPERGNOVA and $\rho$-HESS. **(B)** qq-plot of p value when the true local genetic covariance is set to be 0.

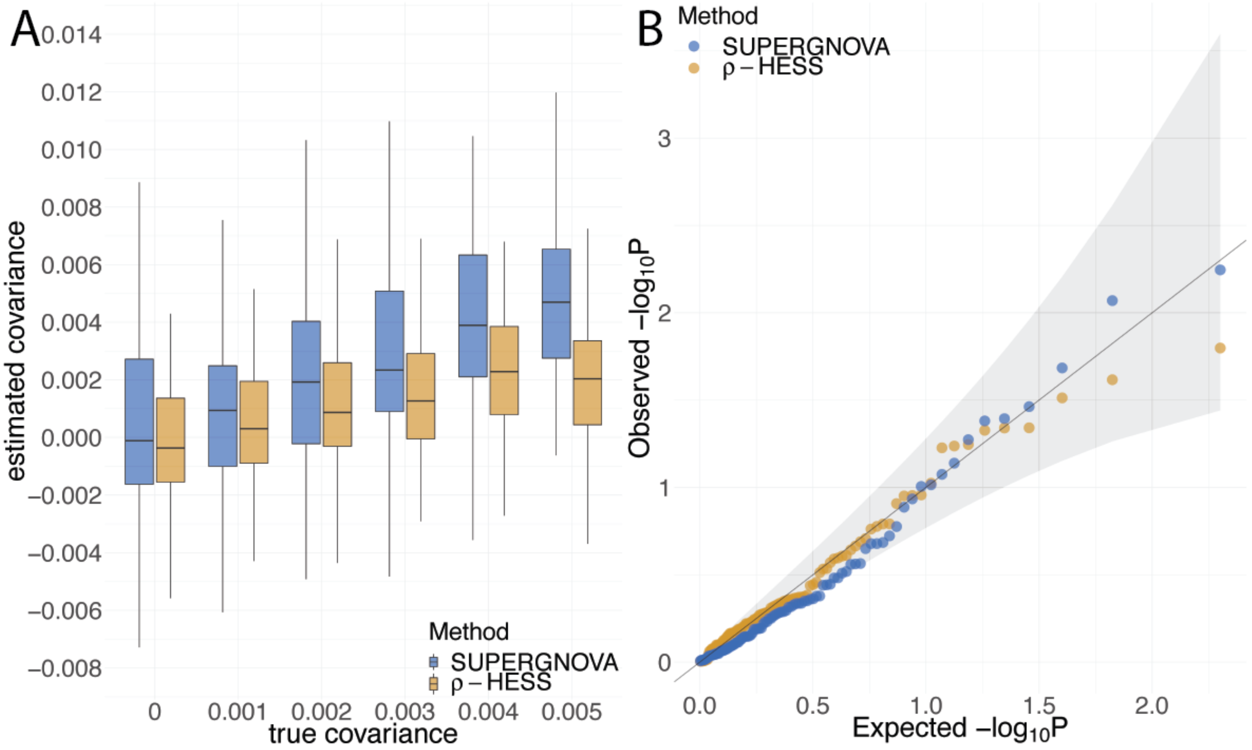

**Supplementary Figure 6. Evaluation of local covariance estimation and type I error control on a completely-overlapping dataset (set 1). (A)** Comparison of local genetic covariance estimates between SUPERGNOVA and $\rho$-HESS. **(B)** qq-plot of p value when the true local genetic covariance is set to be 0.

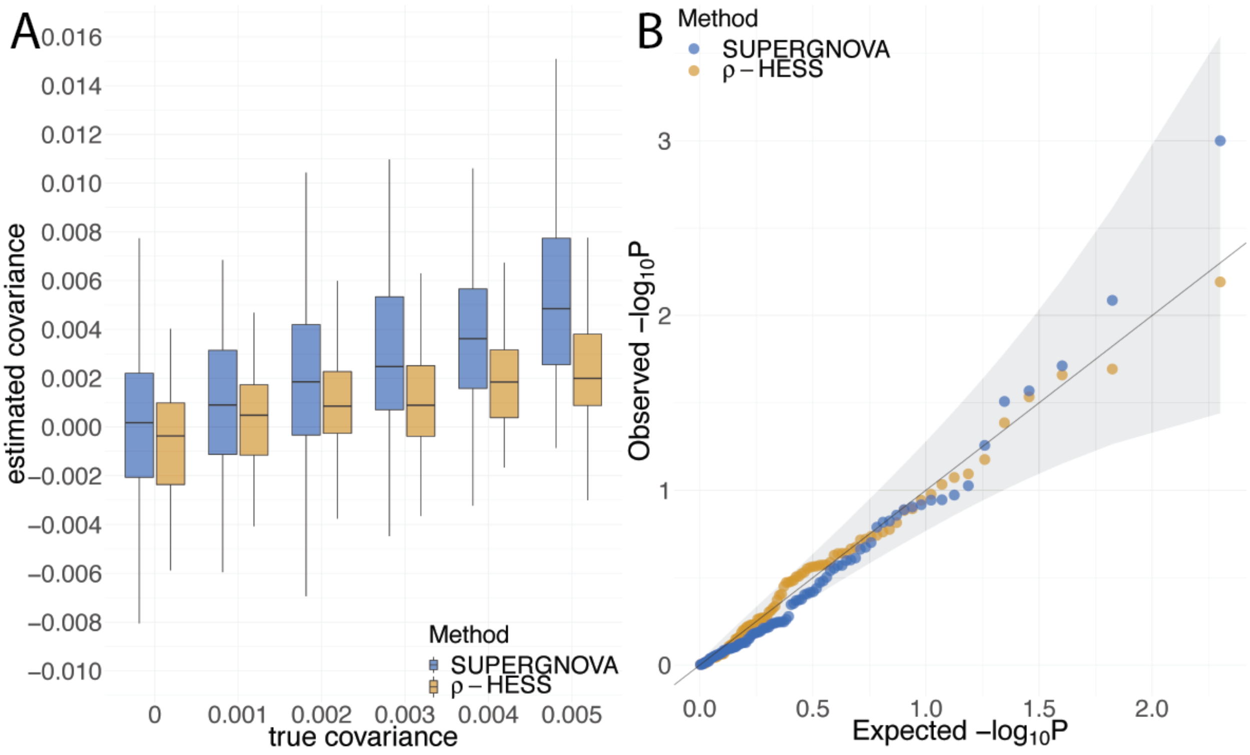

**Supplementary Figure 7. qq-plot for p value of $\rho$-HESS when provided with incorrect overlapping sample size and phenotypic correlation.** In both panels A and B, the true local genetic covariance was set to be 0. We input inaccurate overlapping sample size and phenotypic correlation in **(A)** partially-overlapping datasets (set 1 and set 3) and **(B)** a completely-overlapping dataset (set 1).

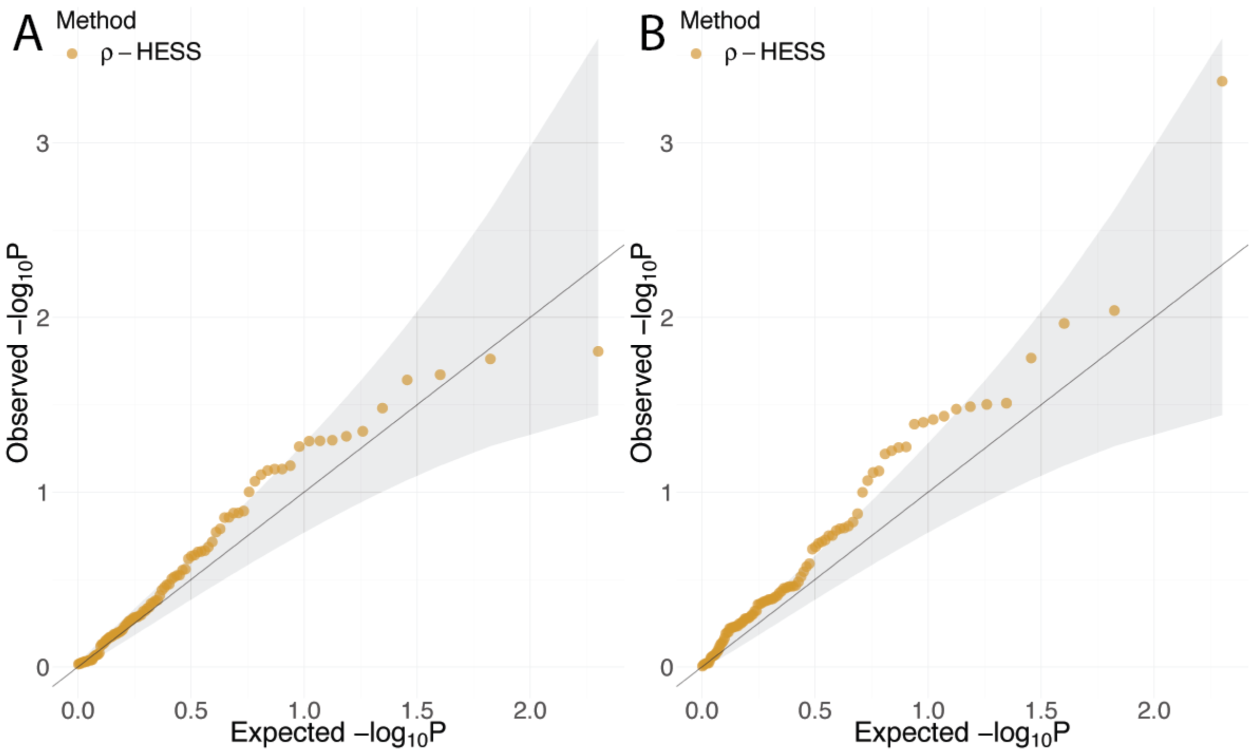

**Supplementary Figure 8. Type I error inflation of $\rho$-HESS under a mis-specified size for shared samples.** The bars show type I error of $\rho$-HESS when provided with incorrect overlapping sample size.

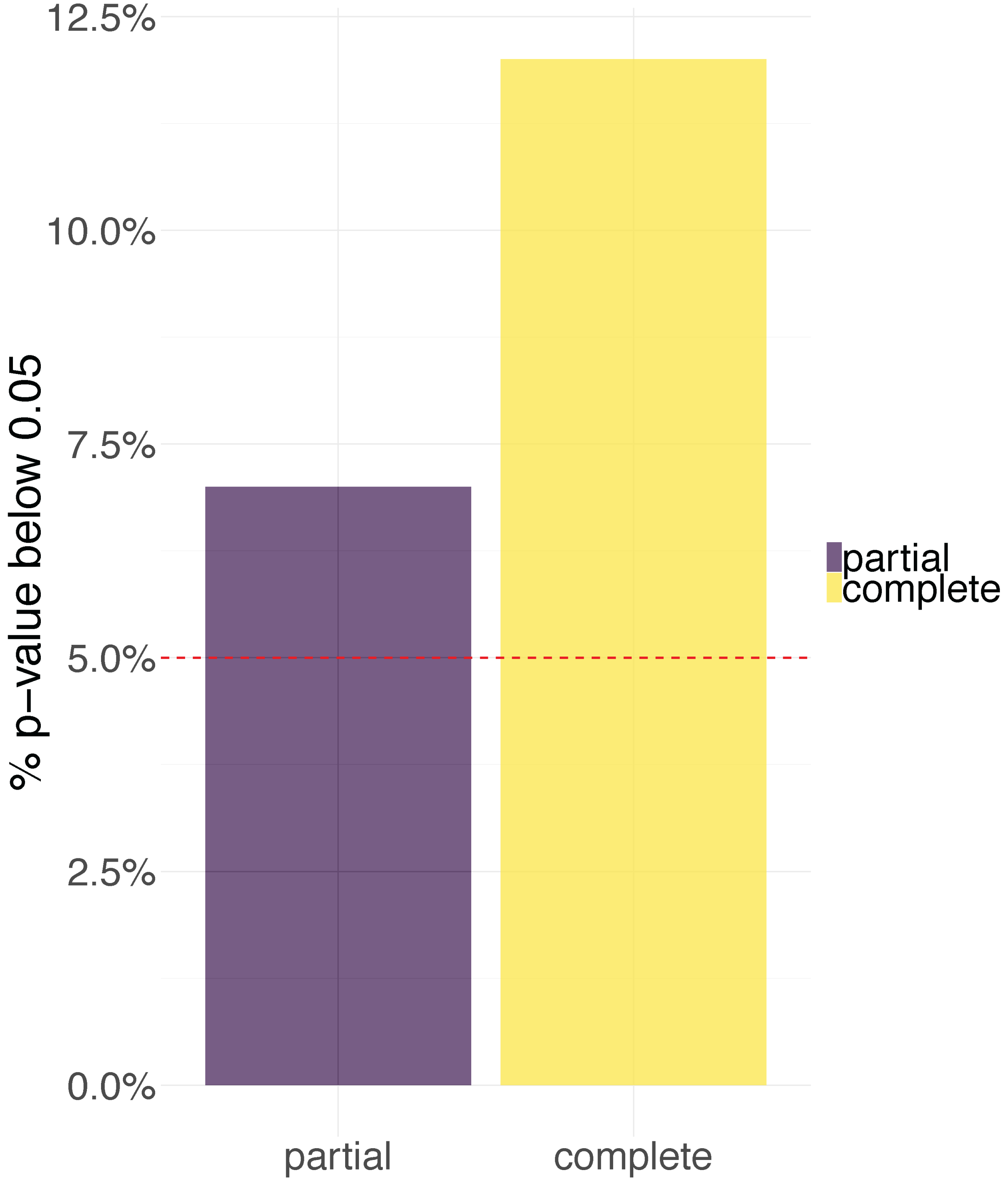

**Supplementary Figure 9. Volcano plot for global genetic correlation.** Each point represents a trait pair. Color of each data point represents the significance and direction of global correlation.

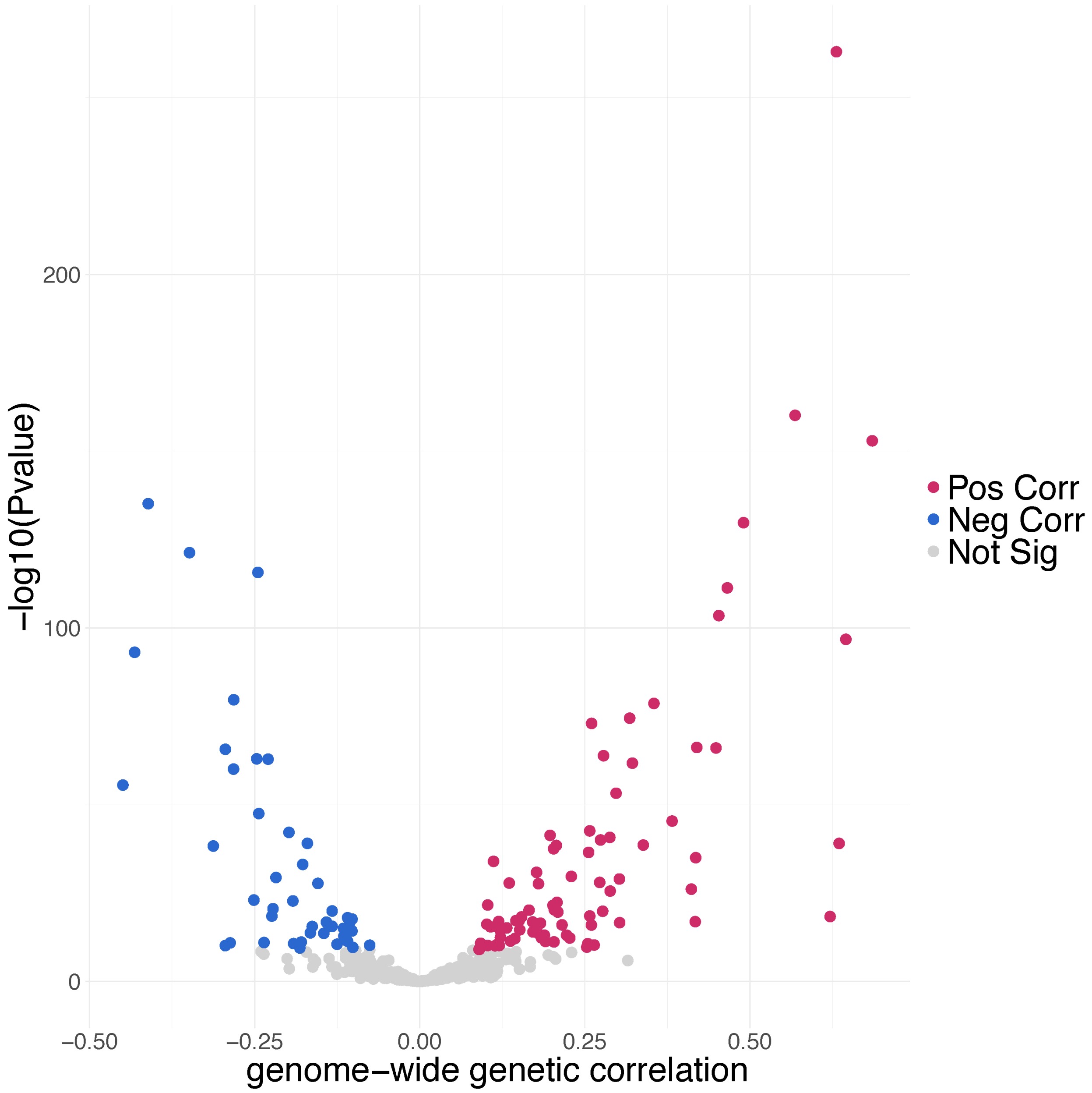

**Supplementary Figure 10. Histograms of z scores of local genetic covariance.** The red lines represent the density function of a standard normal distribution. The trait pairs are **(A)** Crohn-IBD, **(B)** IBD-UC, **(C)** Crohn-UC**, (D)** CP-EA, **(E)** BMI-HDL, and **(F)** ASD-CP.

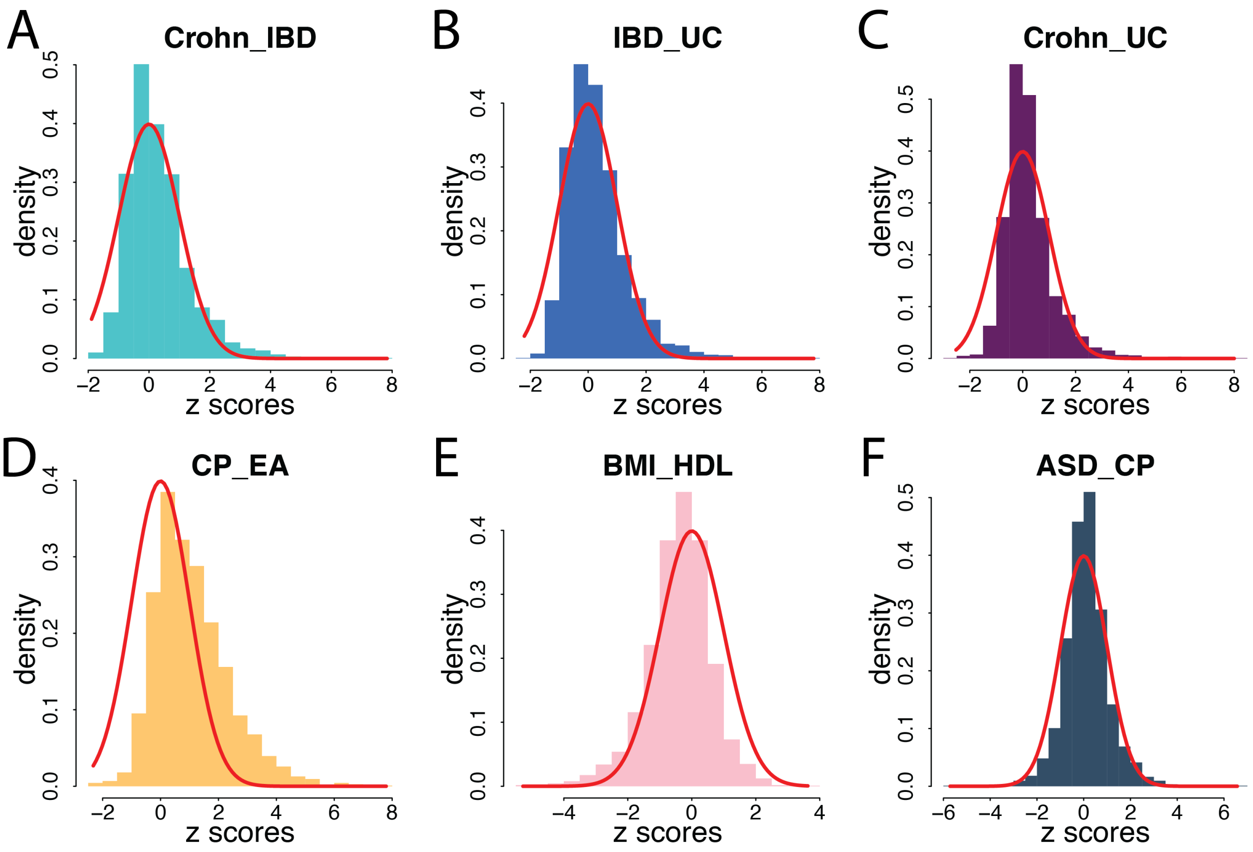

**Supplementary Figure 11. The relationship between the square root of the sample size product and proportion of correlated regions.** Each point represents a pair of traits. Color of each data point denotes the significance and direction of global genetic covariance. The sample size and the proportion of correlated regions showed a modest association (correlation=0.28).

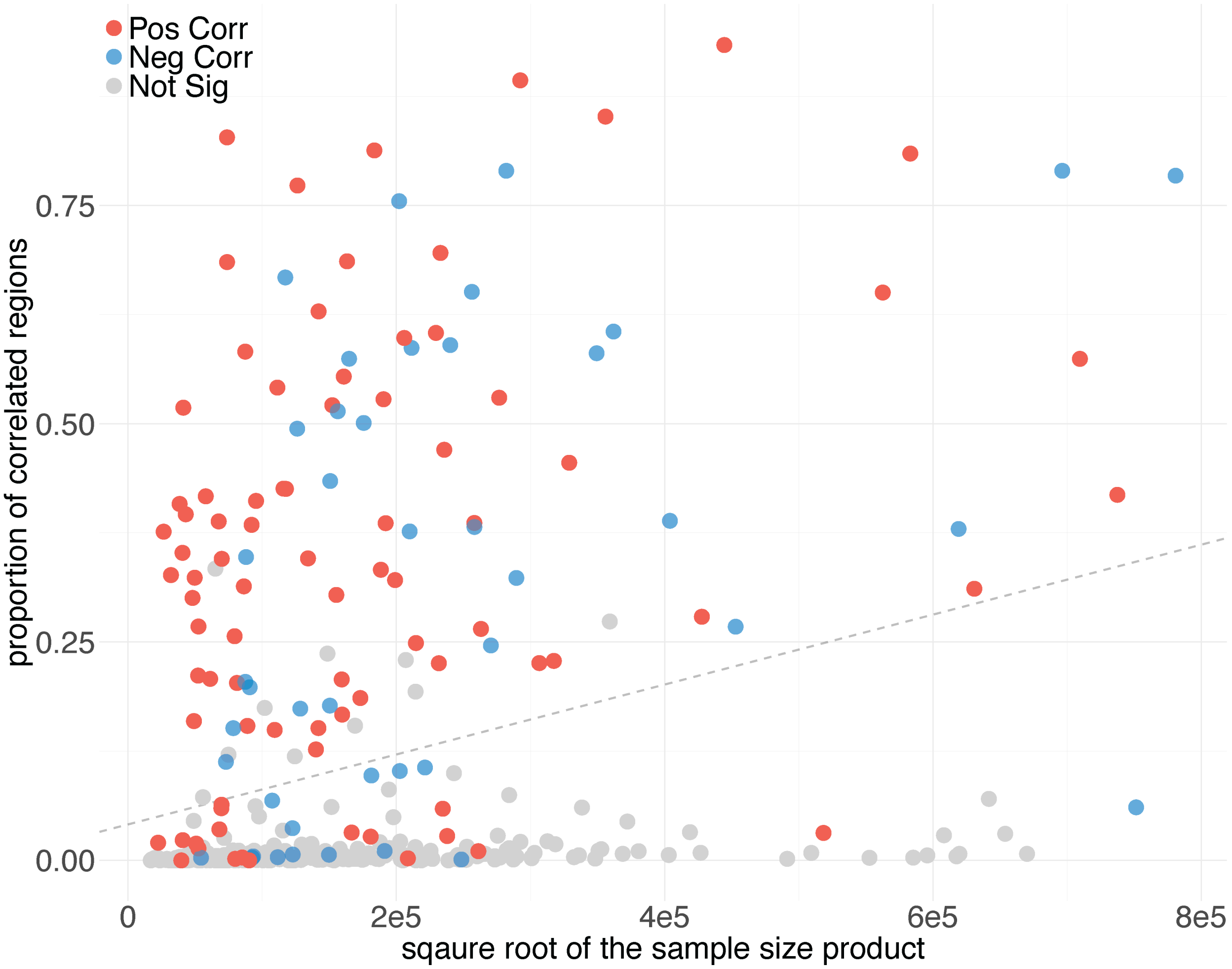

**Supplementary Figure 12. Histograms of z scores of local genetic covariance.** The red lines represent standard normal distribution. These trait pairs were significantly correlated in local regions but not identified as globally correlated, including **(A)** HDL-LDL, **(B)** CP-MDD, **(C)** AXD-OCD, and **(D)** ASD-BD.

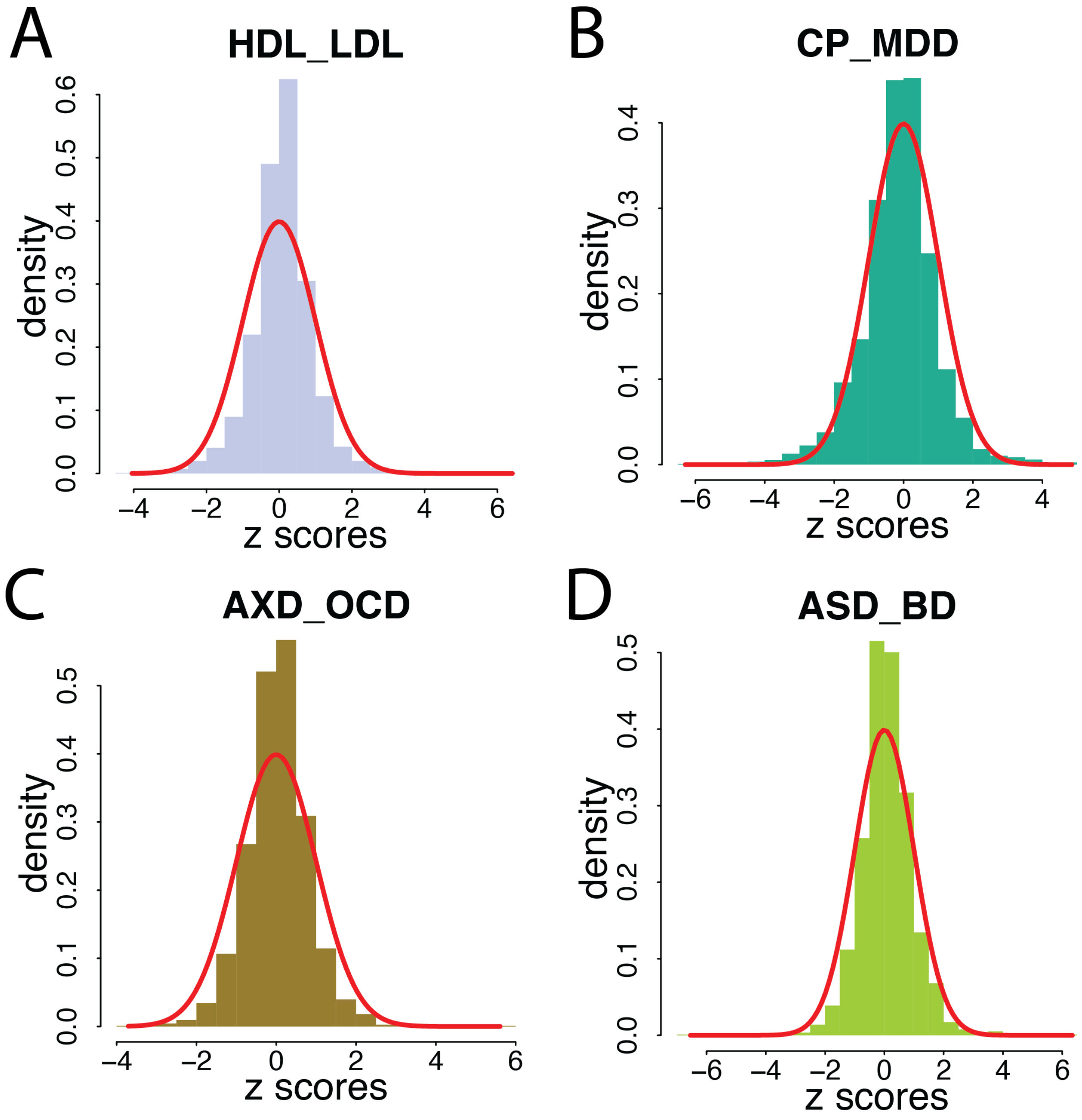

**Supplementary Figure 13. LocusZoom plots for SCZ, EA, SmkInit, DrnkWk, and ADHD at the *BDNF* locus.** ADHD-DrnkWk, ADHD-SCZ, ADHD-SmkInit, and DrnkWk-SCZ are positively correlated in the highlighted region and ADHD-EA and DrnkWk-EA are negatively correlated in the highlighted region.

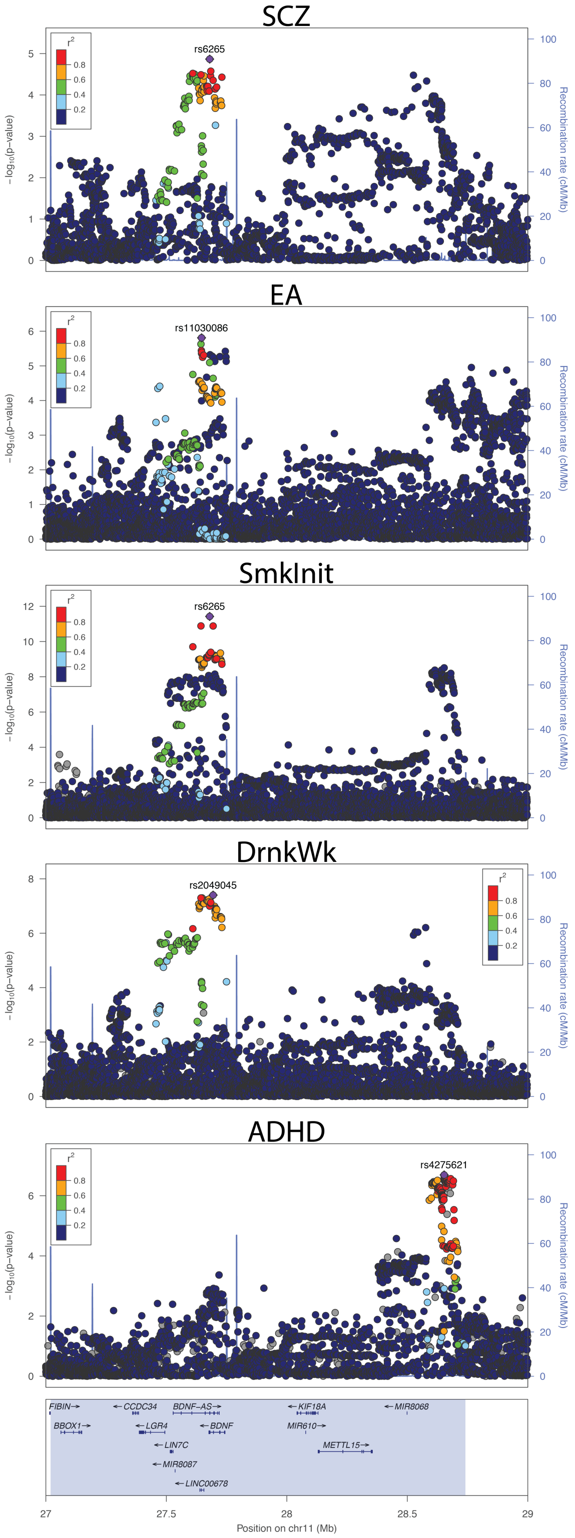

**Supplementary Figure 14. Estimates of local genetic covariance among EA, ADHD, SCZ, DrnkWk, and SmkInit at the *BDNF* locus.** Single asterisks highlight significant genetic covariances after Bonferroni correction for 435 pairs (among the 30 traits). Double asterisks highlight significant genetic covariances after Bonferroni correction for all 1,006,072 regions in 435 trait pairs.

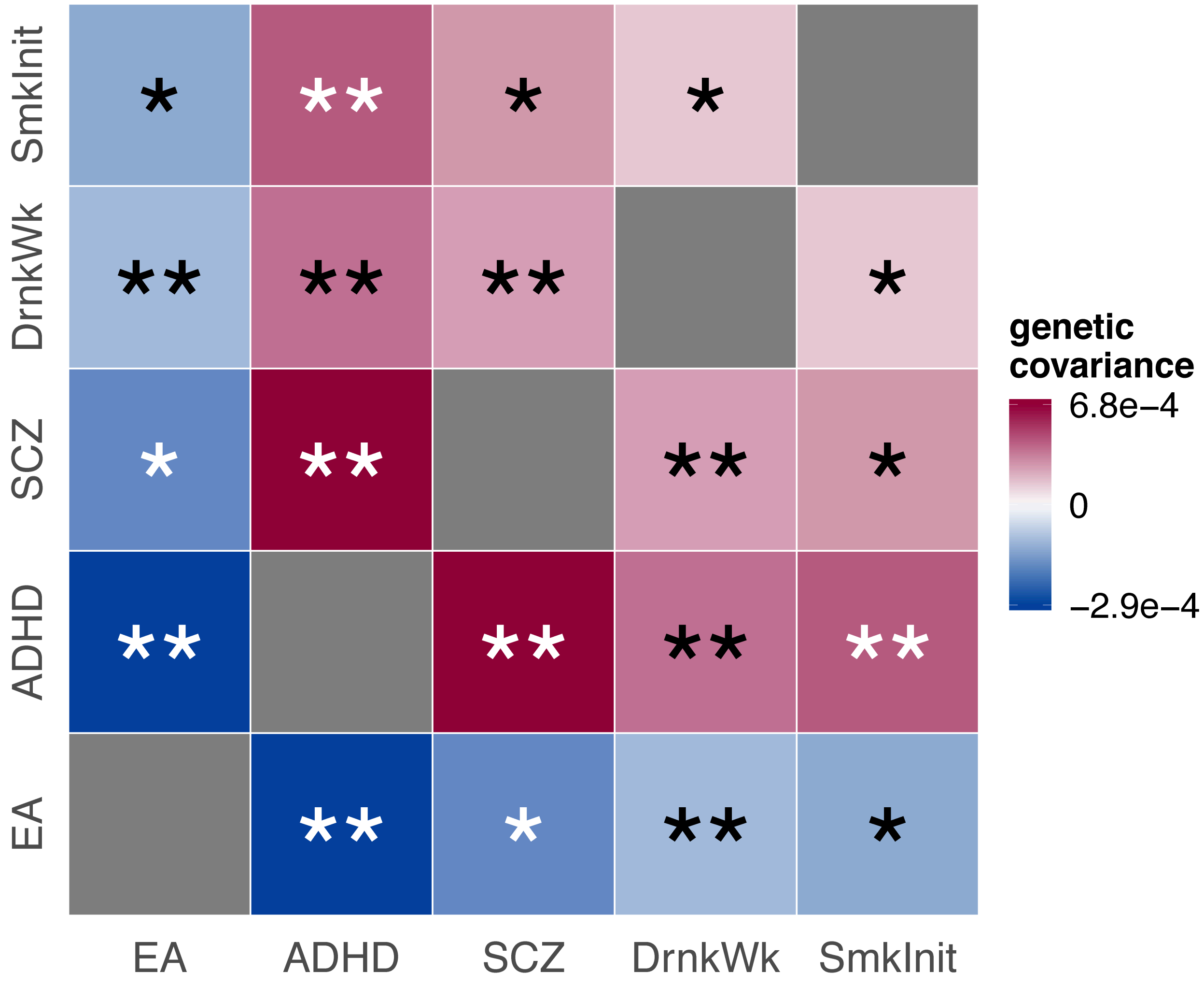

**Supplementary Figure 15. LocusZoom plots for AN, BD, MDD, CP, SCZ, SmkInit, and NSM at the *NCAM1* locus.** AN-SCZ, BD-SmkInit, MDD-SmkInit, NSM-SmkInit, and DrnkWk-SCZ are positively correlated in the highlighted region and AN-CP, CP-SCZ, and CP-SmkInit are negatively correlated in the highlighted region.

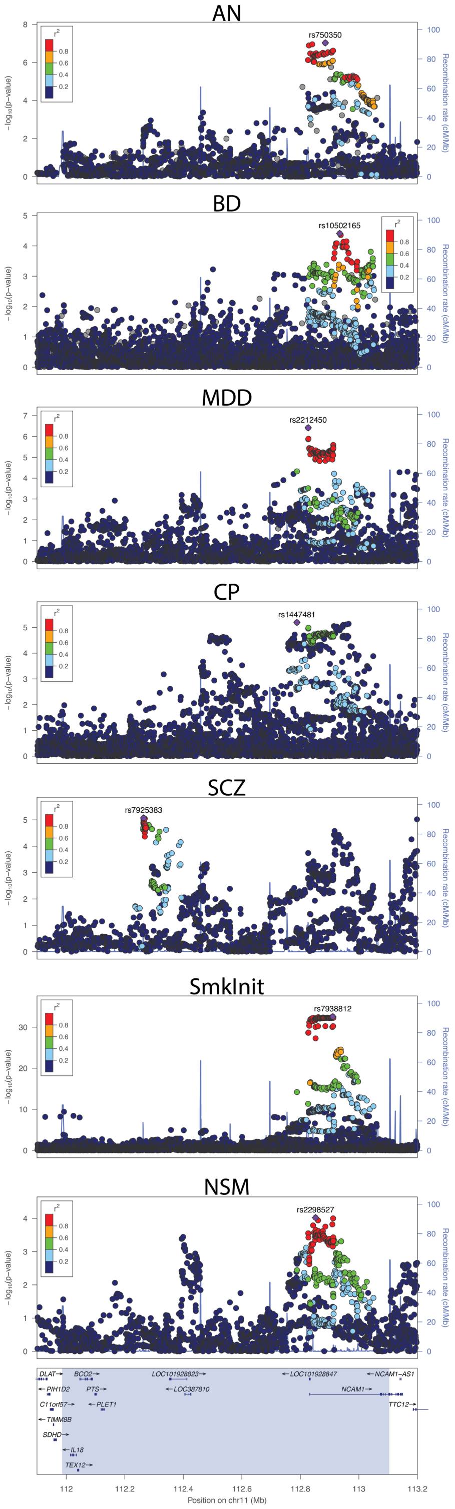

**Supplementary Figure 16. Estimates of local genetic covariance among CP, AN, MDD, NSM, BD, SmkInit, and SCZ at the *NCAM1* locus.** Single asterisks highlight significant genetic covariances after Bonferroni correction for 435 pairs (among the 30 traits). Double asterisks highlight significant genetic covariances after Bonferroni correction for all 1,006,072 regions in 435 trait pairs.

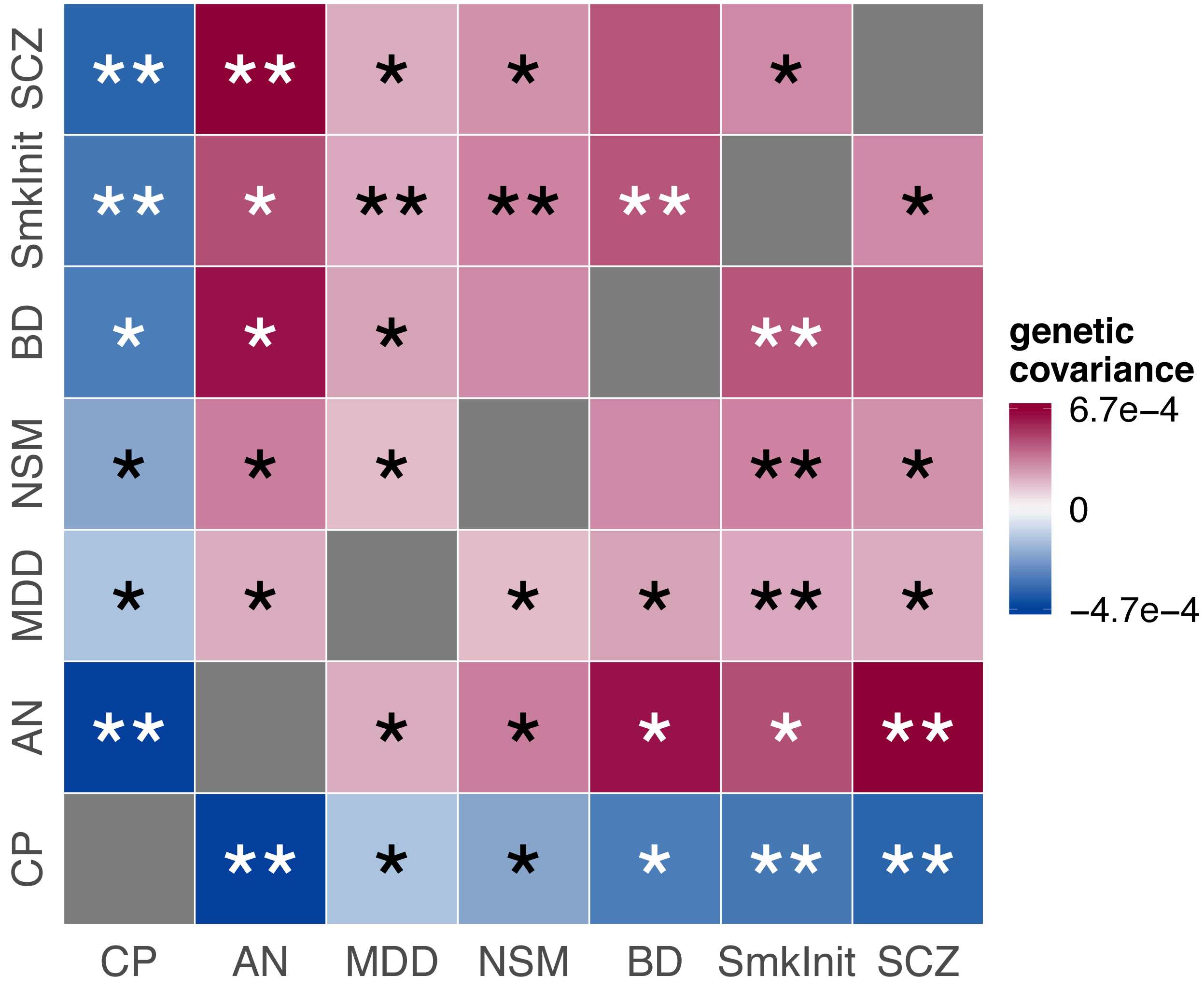

**Supplementary Figure 17. Local genetic covariance at the *SPI1* locus among AD, DrnkWk and NSM. (A)** replications of local genetic covariances estimates using UKBB AD-proxy summary data. The horizontal axis and the vertical axis represent local covariance estimates based on IGAP2019 AD and AD-proxy summary data, respectively. 95% confidence intervals are shown by the gray dashed lines. Panels B-C demonstrate overlap of genes identified by S-PrediXcan for AD, DrnkWk and NSM in **(B)** macrophages and **(C)** monocytes, respectively. **(D)** Genes and local Manhattan plots for AD, DrnkWk, and NSM at the *SPI1* loci where local genetic covariance was identified and successfully replicated among these traits.

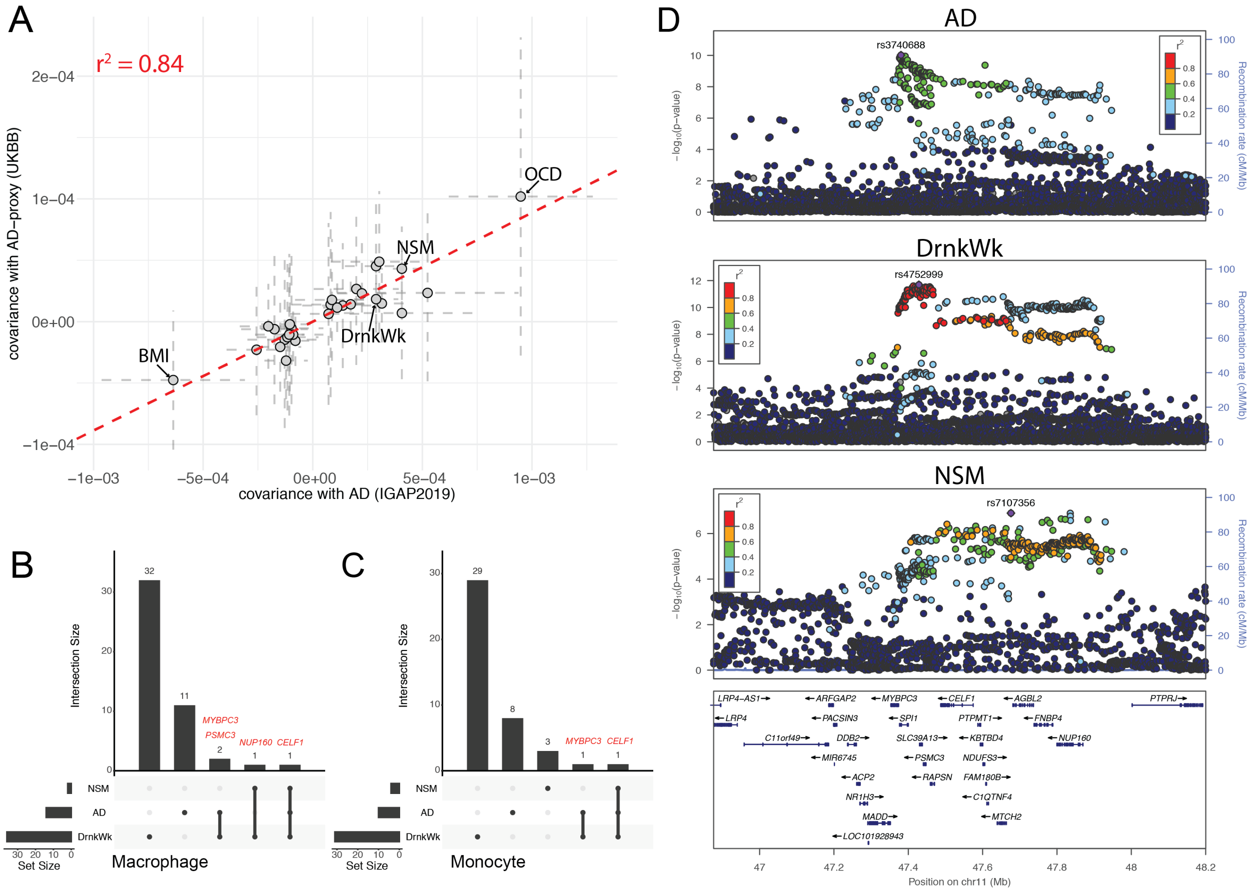

**Supplementary Figure 18. Mirrored Manhattan plots for ASD-CP, ADHD-ASD, and ADHD-CP.** Regions highlighted in red and blue are positively and negatively correlated, respectively, at an FDR cutoff of 0.1.

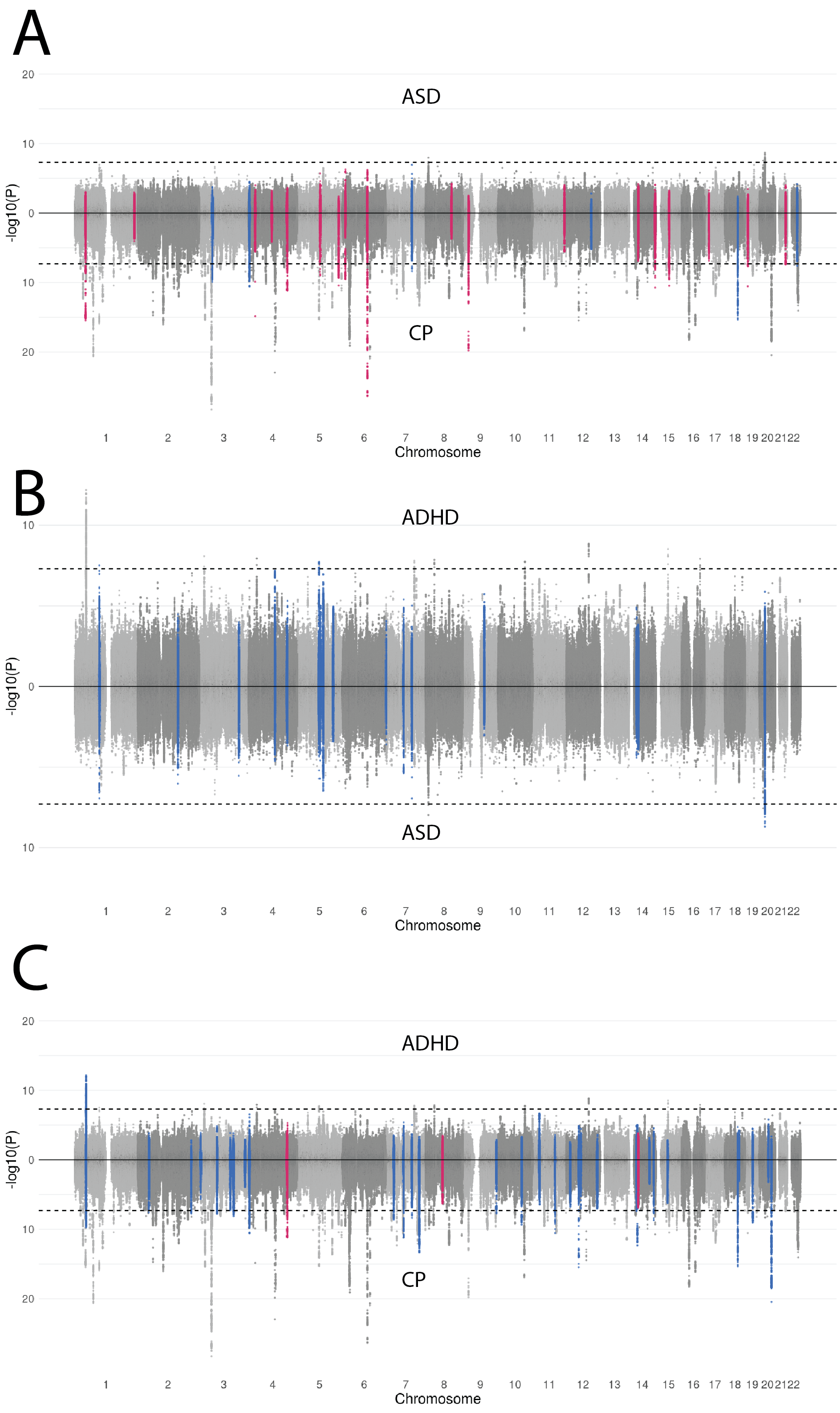

**Supplementary Figure 19. LocusZoom plots for ADHD, ASD, and CP at chromosome 4: 150.5M-153.3M.** ADHD, ASD, and CP are mutually positively corelated in the highlighted region.

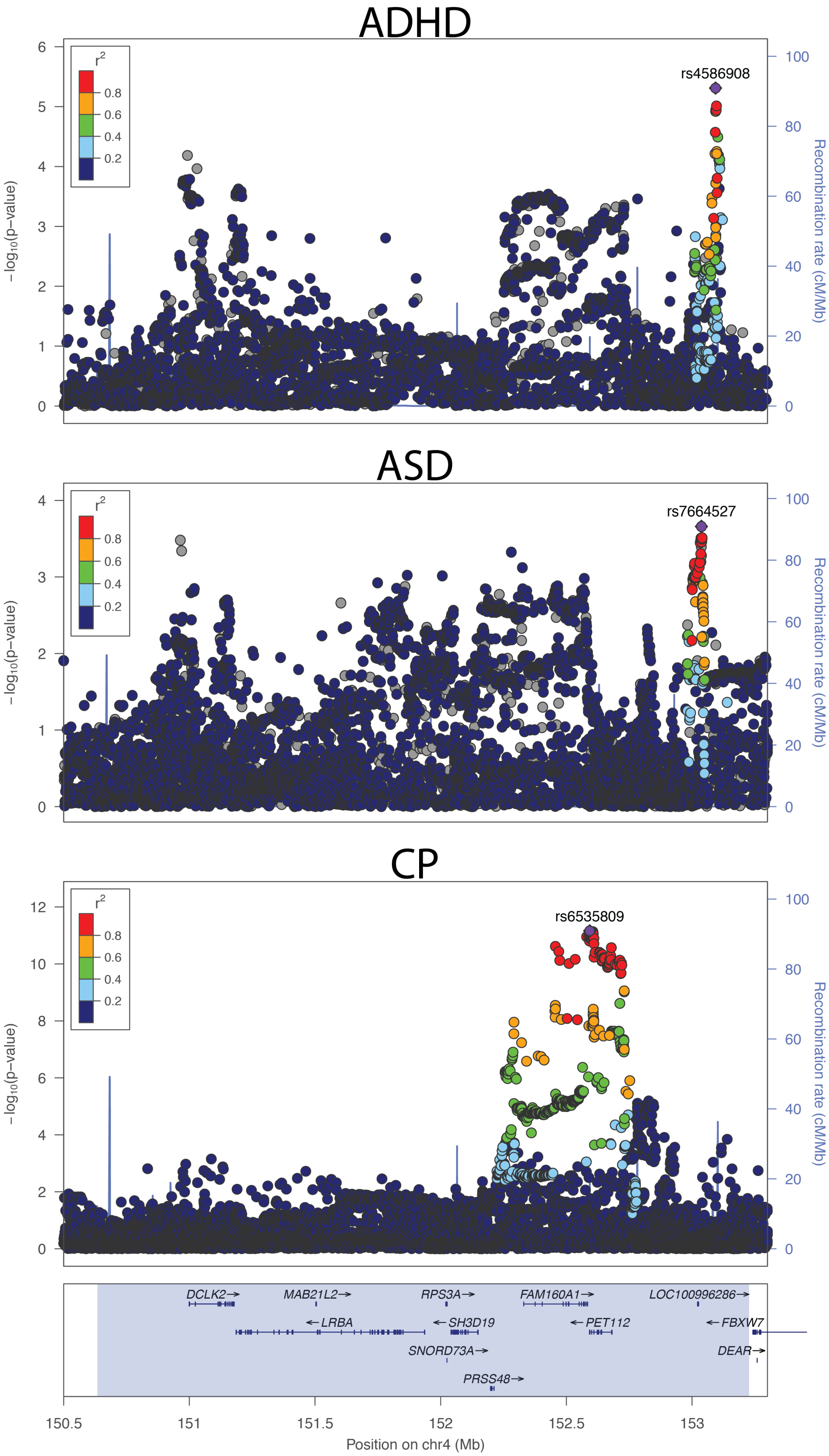

**Supplementary Figure 20. LocusZoom plots for ADHD, ASD, and CP at chromosome 14: 36.5M-38.5M.** ADHD, ASD, and CP are identified as mutually positively correlated at the highlighted region.

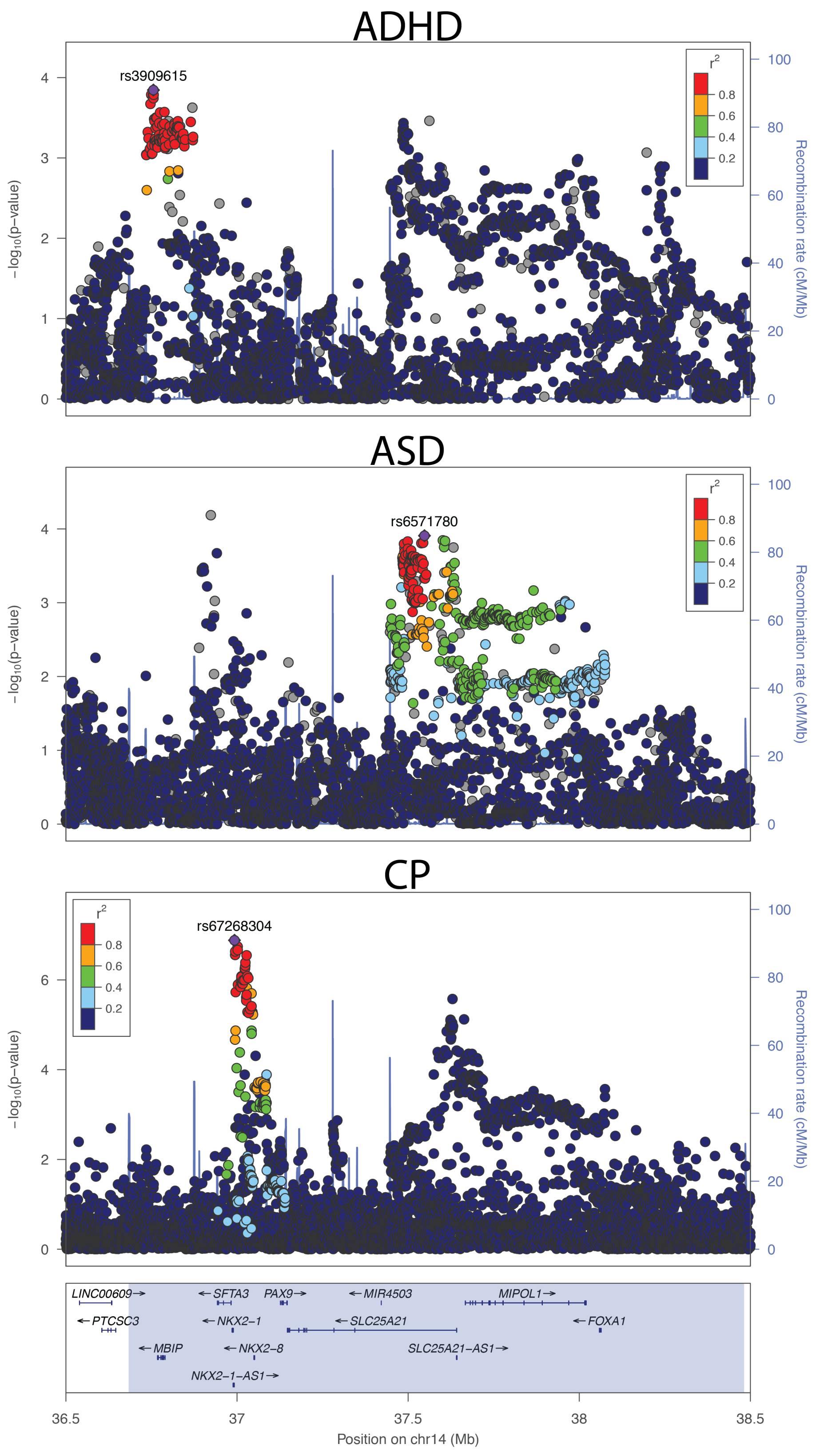

**Supplementary Figure 21. LocusZoom plots for ADHD and SCZ at the *KMT2E* locus.** The SNP in 1000 Genomes 2014 release with the lowest p value in GWAS was chosen to be the index SNP. Besides the correlations among ADHD, ASD and CP, SCZ was also identified to be negatively correlated with CP at the highlighted region.

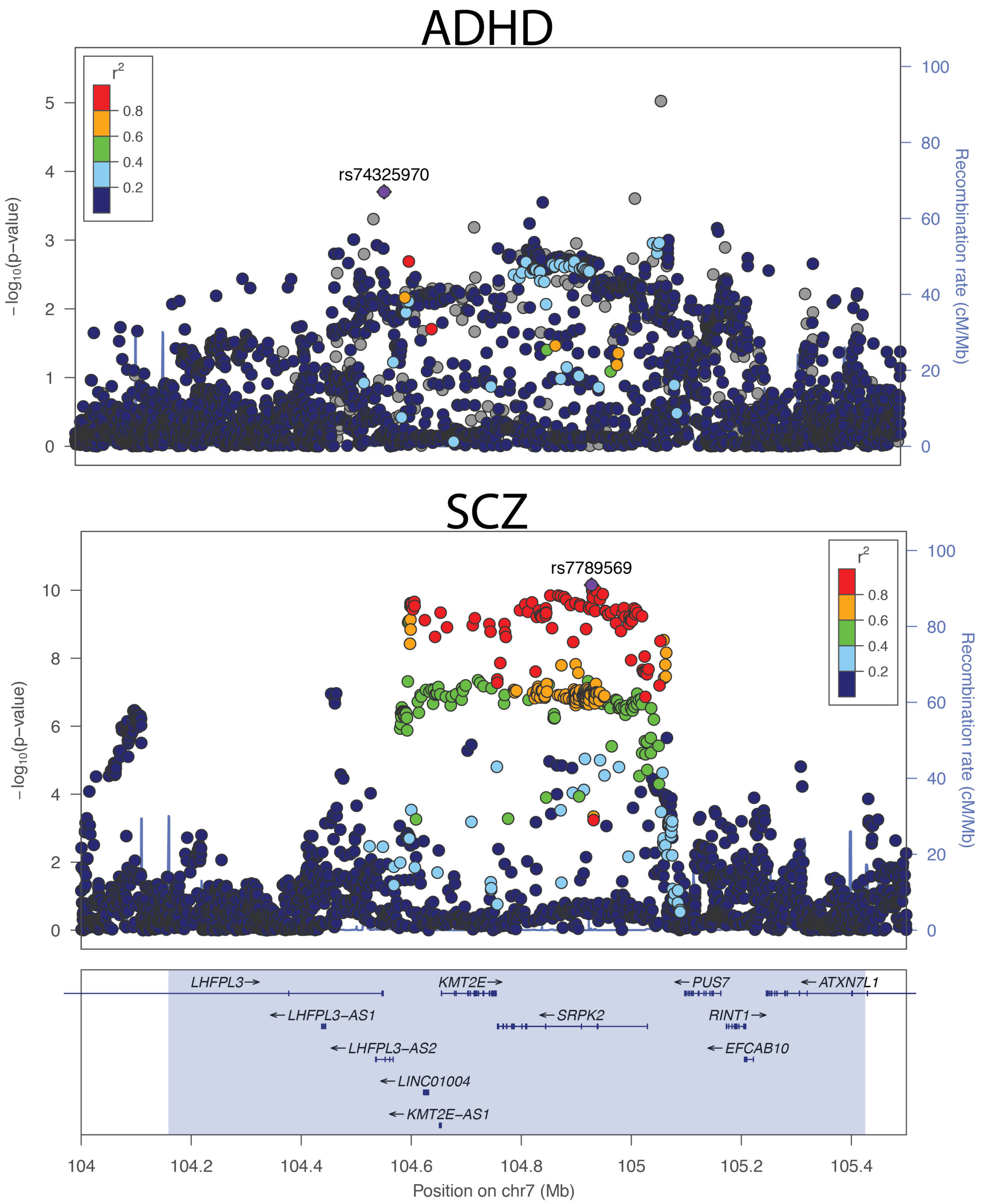

**Supplementary Figure 22. LocusZoom plots for AN, BD, DrnkWk, EA, and SmkInit at the *POU3F2* locus.** SNP in 2014 1000 Genome reference with the least p value for the association is chosen to be the index SNP. Positive correlations are identified among these neuropsychiatric phenotypes at the highlighted region.

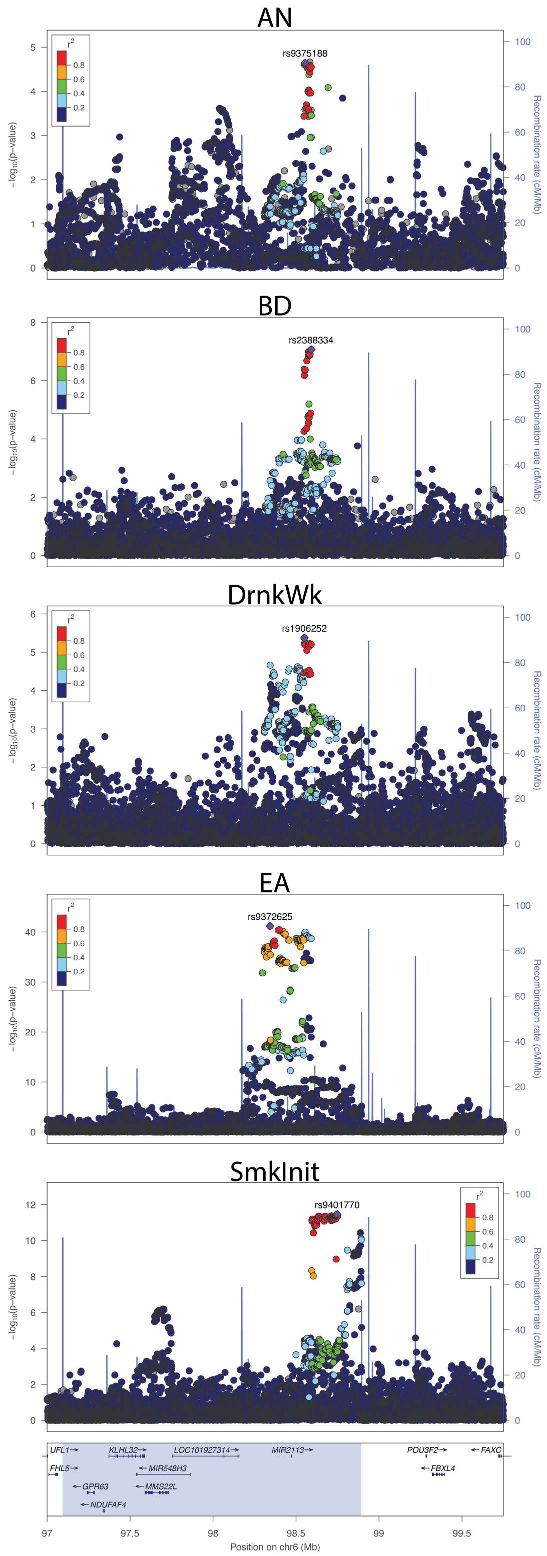

**Supplementary Figure 23. Estimates of local genetic covariance among the 7 correlated neuropsychiatric phenotypes at the *POU3F2* locus.** Asterisks highlight significant genetic covariances after Bonferroni correction for 435 trait pairs.

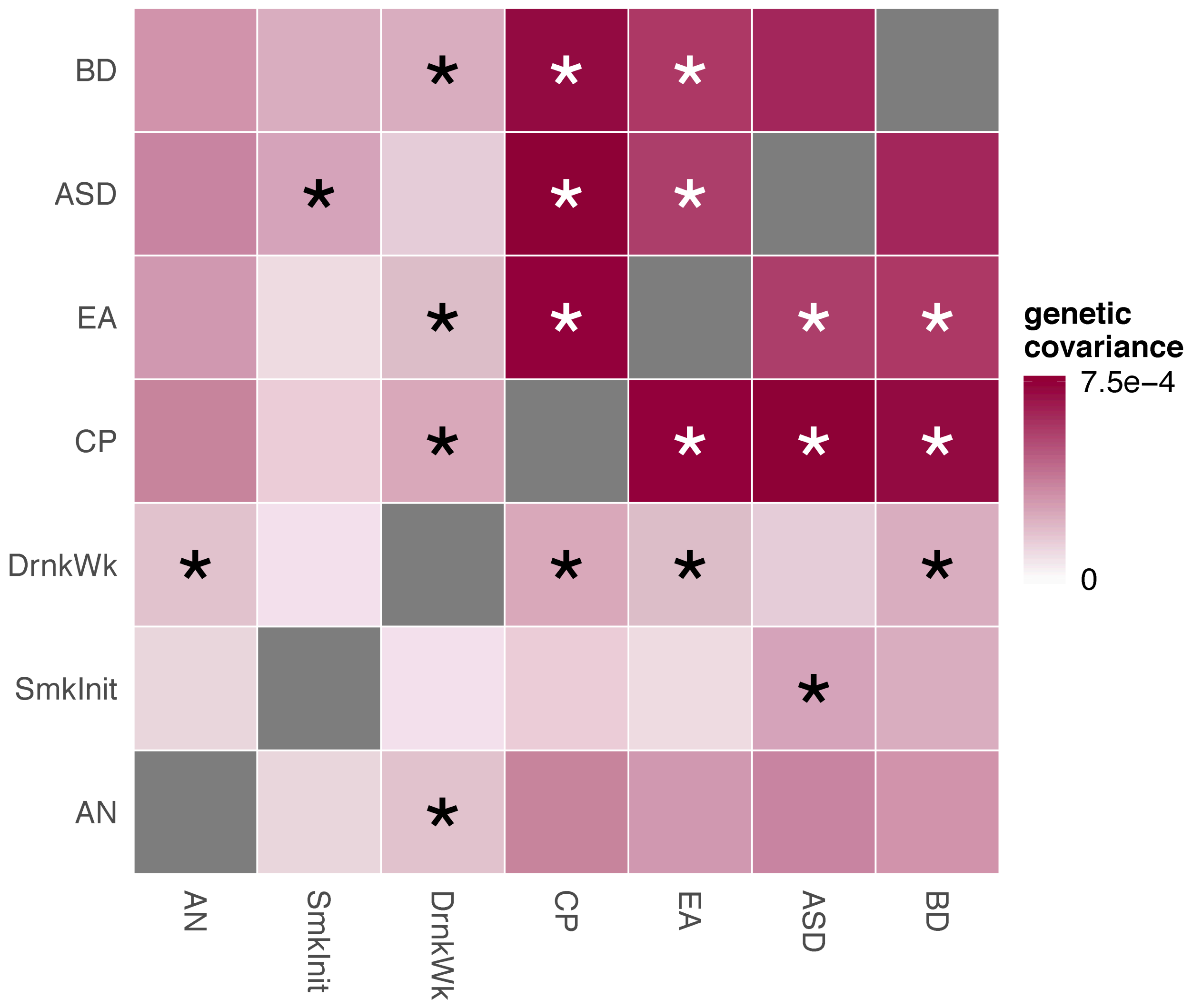

**Supplementary Figure 24. Enrichment for genetic associations with other complex traits in regions with opposite correlations between ASD and CP.** Odds ratio values are labeled next to each bar. The red dashed lines mark the p-value cutoff of 0.05 and the black dashed lines denote the p-value thresholds after Bonferroni correction (p=7.5e-5).

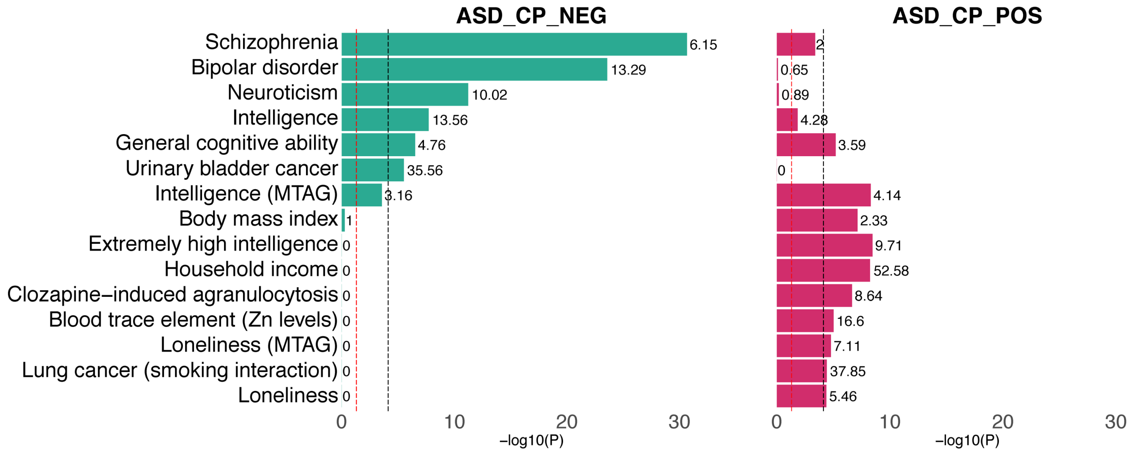

**Supplementary Figure 25.** Stratified genetic covariance of 28 complex traits with ASD and CP in regions identified with significant ASD-CP covariance. Red intervals denote genetic covariance in regions with a positive ASD-CP covariance. Blue intervals denote genetic covariance in regions with a negative ASD-CP covariance. Triangle dots denote genetic covariance between ASD and other traits. Circle dots denote genetic covariance between CP and other traits. Traits with a suggestive genetic covariance with either ASD or CP (p<0.05) are shown. Asterisks highlight significant covariances after Bonferroni correction. The interval length represents standard errors.

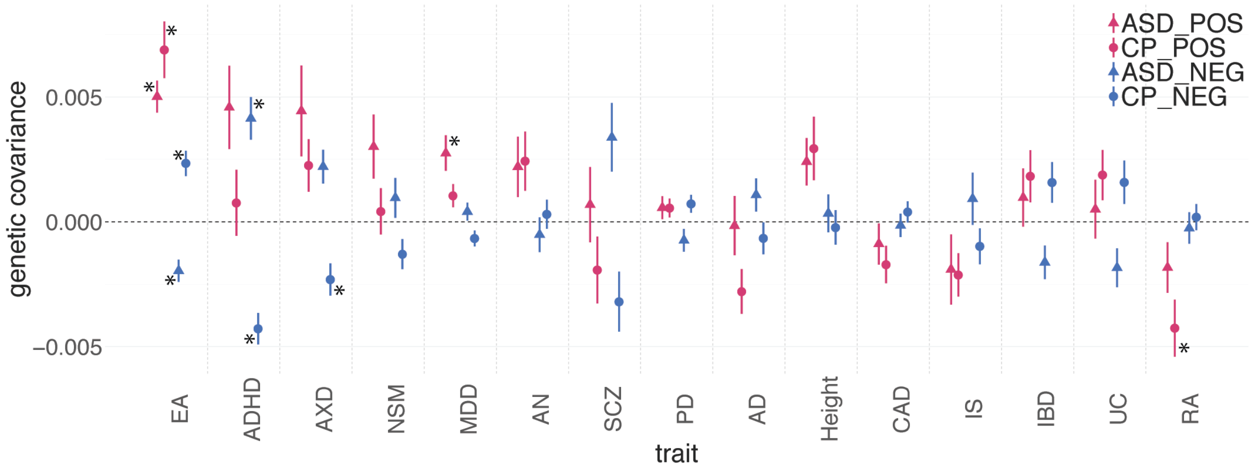

**Supplementary Figure 26. Cumulative gene expression rate in fetal brain of ASD-CP positively and negatively correlated genes and other genes across developmental stages.** Standard error of gene expression rate estimation for ASD-CP positively and negatively correlated genes and other genes is shown.

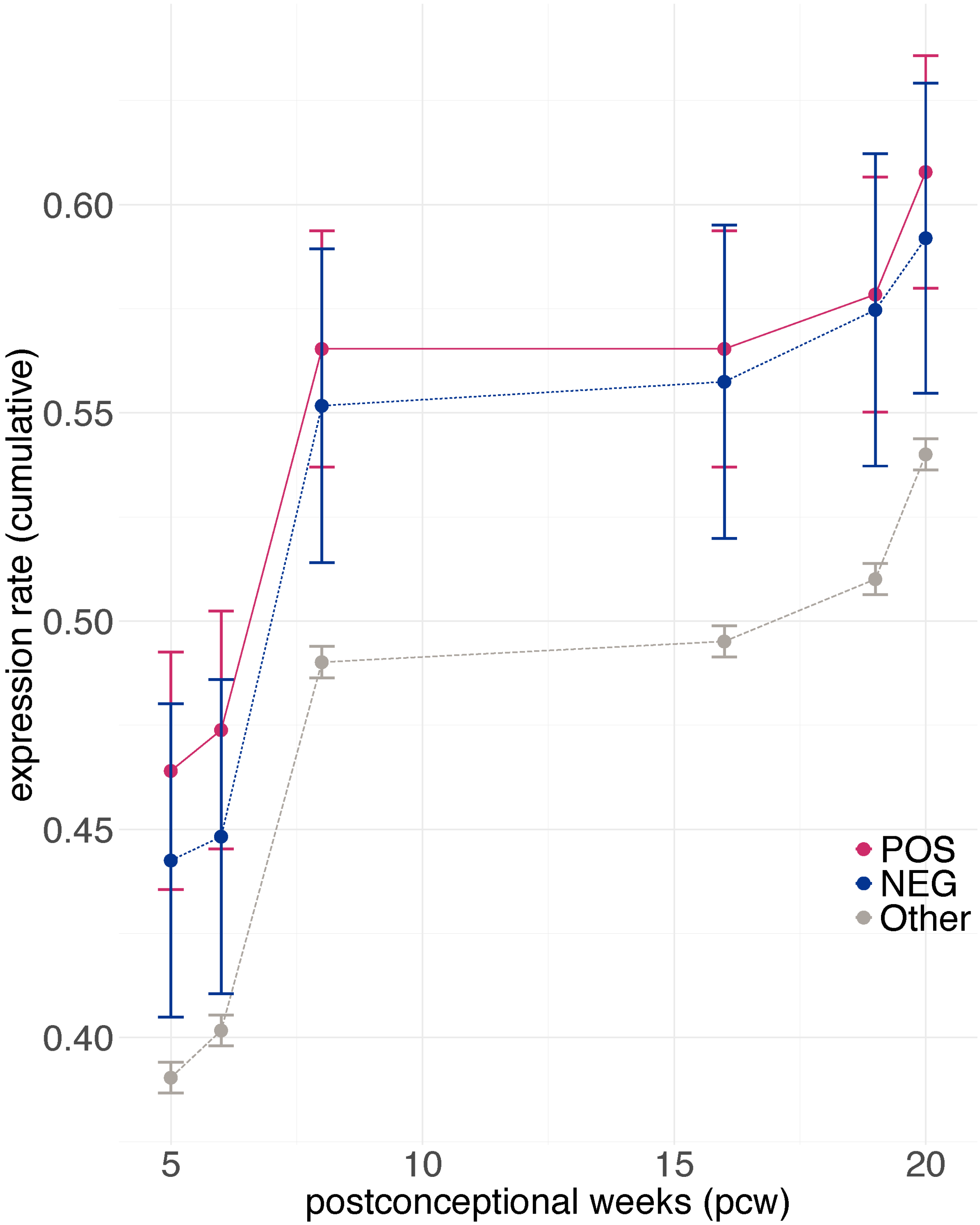

**Supplementary Figure 27. Gene expression in brain across developmental stages.** Means of log-transformed RPKM and standard error are shown. The black dashed line represents neonatal time. The red line, blue line and gray line represents expression levels of ASD-CP positively correlated, negatively correlated and other background genes, respectively.

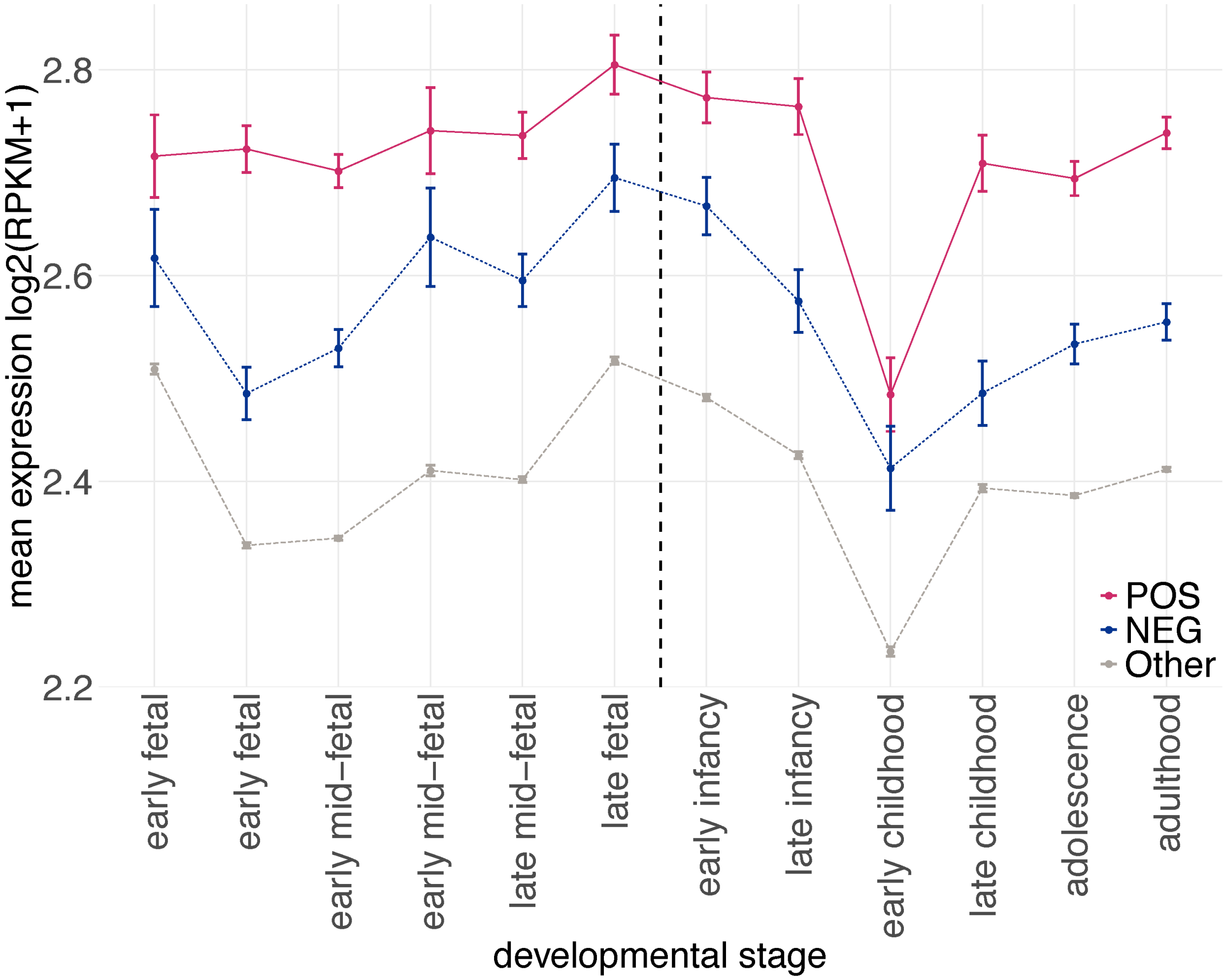

**Supplementary Figure 28. Normality of PRSs.** We normalized PRS+ and PRS- of ASD probands with sample mean and standard deviation. In panel A-B, we used qq-plot to compare the distributions of normalized **(A)** PRS+ and **(B)** PRS- with standard normal distribution.

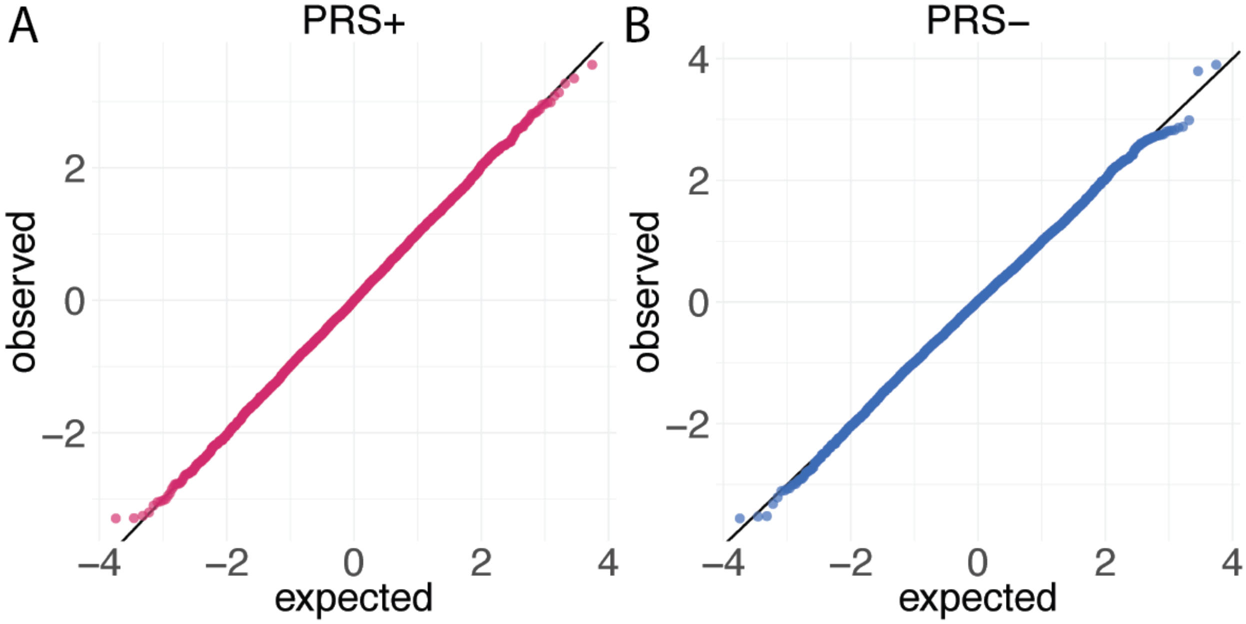

**Supplementary Figure 29. Distribution of IQ in ASD probands with extreme PRS+ and PRS- values.** The distribution was estimated by the Gaussian kernel approach.

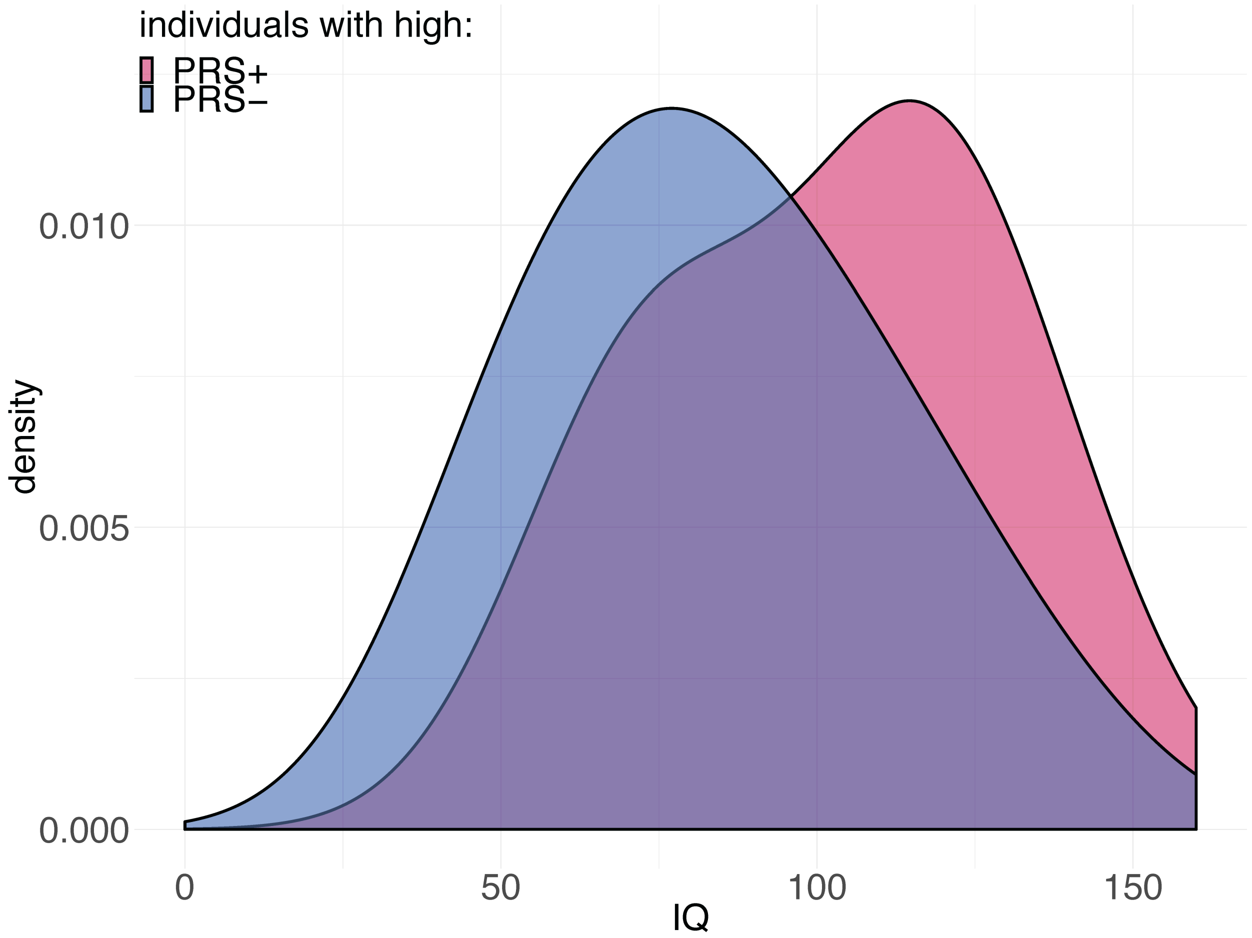

**Supplementary Figure 30. Enrichment of ASD subtypes in extreme PRS+ and PRS- groups.** Each bar indicates the fold enrichment of a subtype in ASD probands with extreme PRS. The p values of enrichment are annotated to the top of each bar. The dashed line represents enrichment=1. Autism spectrum disorder represents Rett disorder and childhood disintegrative disorder. PPD-NOS represents pervasive developmental disorder – not otherwise specified.

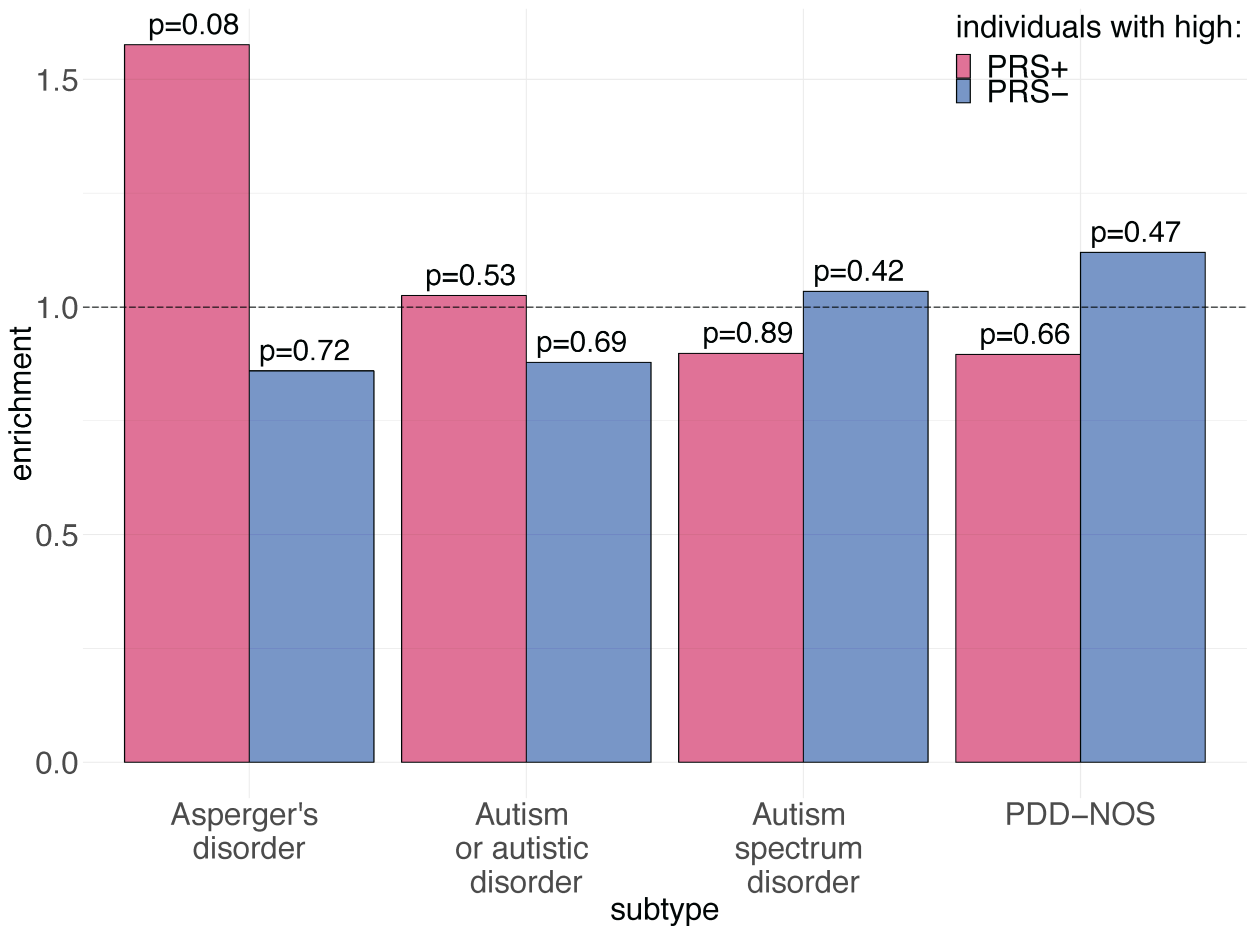

**Supplementary Figure 31. Theoretical variance and empirical variance of estimates of local genetic covariance.** The x-axis is the cutoff for eigenvalues included in the estimation of local genetic covariance. Theoretical variance decreases with the increase of $K_i$ and empirical variance increase rapidly when the cutoff is small enough. We used the data from one example of simulation to plot this figure.

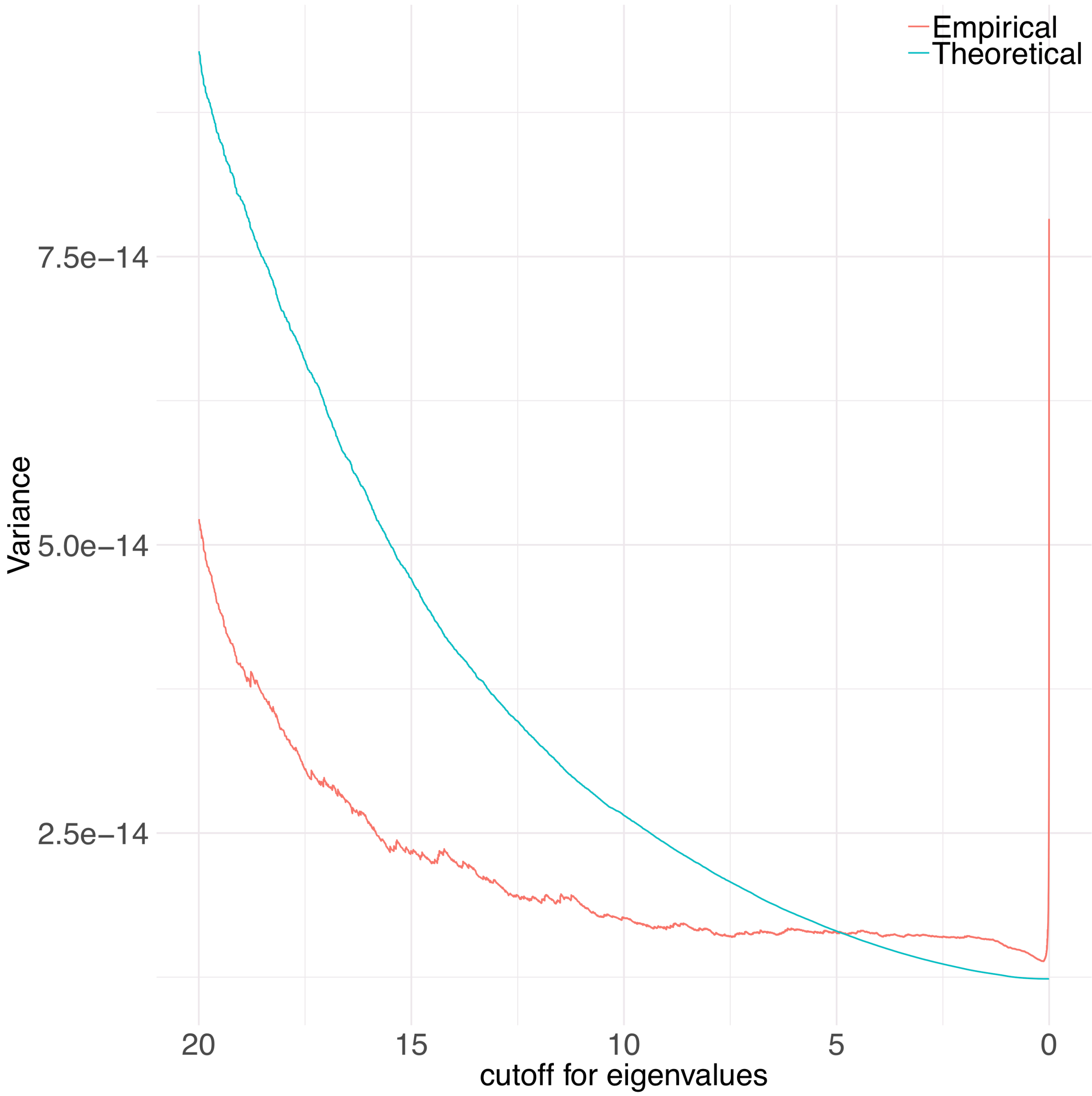
